# Integrated structural dynamics uncover new modes of B_12_ photoreceptor activation

**DOI:** 10.1101/2025.08.13.669063

**Authors:** Ronald Rios-Santacruz, Harshwardhan Poddar, Kevin Pounot, Derren J. Heyes, Nicolas Coquelle, Megan J. Mackintosh, Linus O. Johannissen, Sara Schianchi, Laura N. Jeffreys, Elke De Zitter, Rory Munro, Martin Appleby, Danny Axford, Emma Beale, Matthew J. Cliff, Maria Davila, Sylvain Engilberge, Guillaume Gotthard, Kyprianos Hadjidemetriou, Samantha J. O. Hardman, Sam Horrell, Jochen S. Hub, Kotone Ishihara, Sofia Jaho, Gabriel Karras, Machika Kataoka, Ryohei Kawakami, Thomas Mason, Hideo Okumura, Shigeki Owada, Robin L. Owen, Antoine Royant, Annica Saaret, Michiyo Sakuma, Muralidharan Shanmugam, Hiroshi Sugimoto, Kensuke Tono, Ninon Zala, John H. Beale, Takehiko Tosha, Jacques-Philippe Colletier, Matteo Levantino, Sam Hay, Pawel M. Kozlowski, David Leys, Nigel S. Scrutton, Martin Weik, Giorgio Schirò

**Author notes:** Contributed equally (Ronald Rios-Santacruz, Harshwardhan Poddar, Kevin Pounot, Derren J. Heyes, Nicolas Coquelle, Megan J. Mackintosh).

## Abstract

Photoreceptor proteins initiate, regulate and control fundamental biological processes such as vision, photosynthesis and circadian rhythms^1^. A large photoreceptor subfamily uses vitamin B_12_ derivatives for light sensing^2^, contrasting with the well-established mode of action of these organometallic derivatives in thermally activated enzymatic reactions^3^. The molecular mechanism of B_12_ photoreception and how this differs to the thermal pathways remain unknown. Here we provide a detailed spatio-temporal description of photoactivation in the prototypical tetrameric B_12_ photoreceptor CarH^4,5^ from nanoseconds to seconds by using an integrative approach, combining time- and temperature-resolved structural and spectroscopic methods with quantum chemical calculations. High resolution structural snapshots of key intermediates illustrate how photocleavage of a Co–C bond triggers a pathway of structural changes that propagate throughout CarH from the B_12_ chromophore, via a previously unknown adduct, to finally cause tetramer dissociation. These unique intermediates, which differentiate CarH from thermally-activated B_12_ enzymes, steer the photoactivation pathway and act as the molecular bridge between photochemical and photobiological timescales. Our results offer a spatio-temporal understanding of CarH photoactivation and pave the way for designing and optimising B_12_-dependent photoreceptors for future optogenetic applications.

## Main

Photoreceptor proteins absorb light to elicit a biological response, controlling processes such as vision, circadian rhythms and plant development. Several photoreceptor families have been identified harbouring a range of chromophores [e.g. retinal, flavin adenine dinucleotide (FAD), bilin, p-coumaric acid, ketocarotenoids]^1^. How the initial photochemical events at the chromophore translate into the desired biological outcome is a current focus of intense research^6^. In particular, it is unclear how photoreceptors follow a specific reaction pathway over multiple time scales and prevent premature energy dissipation^7^. Recently, a new photoreceptor family was discovered that repurposes the ubiquitous organometallic enzyme cofactor vitamin B_12_ for light sensing^2^ and is estimated to include several thousand members^8,9^. This family is sensitive to the green light region of the visible spectrum. Its photoresponse can be extended to the red either naturally in photocobilins, a recently discovered subfamily that uses red-light absorbing linear tetrapyrrole biliverdin in close proximity to the B_12_ cofactor^10^, or artificially^11,12^. B_12_-dependent photoreceptors have become an exciting target for a number of light-dependent biotechnological and optogenetic applications, such as in the formation of light-responsive hydrogels for drug delivery, light-activated technology devices and the regulation of mammalian gene expression^13-18^. A detailed spatio-temporal description of the photoresponse in B_12_-photoreceptors is, therefore, of fundamental and applied interest.

CarH from *Thermus thermophilus* is a model B_12_ photoreceptor, regulating carotenoid gene expression to mitigate photooxidative stress under light exposure^4^. Monomeric CarH binds coenzyme B_12_ or adenosylcobalamin (AdoCbl, Extended Data Fig. 1), triggering formation of a homotetramer in the dark that binds to DNA and suppresses gene transcription^5^. Illumination with green (or blue) light cleaves the photolabile Co–C5′ bond and leads to the release of the final photoproduct 4′, 5′ anhydroadenosine^19^ by an, as yet, unknown mechanism^20^. The ensuing O_2_-dependent Co redox state changes drive protein ligand switches that culminate in the formation of the light-adapted state, a stable bis-histidine cobalamin adduct^5,21^ (Supplementary Fig. 1), and tetramer dissociation, thereby enabling transcription^4^. What distinguishes CarH from the thermally-activated B_12_ enzymes, allowing it to harness the inherent photochemical properties of the B_12_ chromophore, remains unclear, and likewise for the structural changes underpinning the photochemical mechanism.

Here we used an integrated experimental and computational approach to study light-induced structural changes spanning over nine orders of magnitude in time, from nanoseconds to seconds. We show that homolytic cleavage of the Co–C5′ bond leads to formation of a previously unknown Co–C4′ adduct, followed by downstream, slower structural changes that result in tetramer dissociation. By revealing the structural changes connecting photochemical and photobiological events our results on CarH yield a holistic spatio-temporal understanding of B_12_ photoreceptor activation.

### An integrated experimental and computational approach sheds light on photoactivation mechanism from photolysis to downstream structural changes

To elucidate the structural and chemical changes following photon absorption and formation of the light-adapted state, we combined time- and temperature-resolved structural and spectroscopic methods with quantum chemical calculations. We studied the tetrameric cobalamin binding domains of *Thermus thermophilus* CarH (*Tt*CBD), each consisting of a Rossmann fold and a four-helix bundle domain, i.e. without the N-terminal DNA-binding domain^4,21^. Time-resolved UV-vis absorption spectral changes, monitored following ns excitation with green light (530 nm) of *Tt*CBD in solution (see Supplementary Note 1 and Extended Data Fig. 2), reveal similar kinetics to those reported for full-length CarH^22^, validating the use of *Tt*CBD as our model.

Formation of the light-adapted state occurs via a series of spectroscopically-distinct intermediates formed with respective time constants of < 10 ns^22,23^, ∼ 500 ns^22^, ∼ 7 ms and ∼ 600 ms (Supplementary Note 1 and Extended Data Fig. 2). Importantly, UV-vis spectral changes on the ns-ms time scale of crystalline samples are not impacted by crystal packing (Supplementary Fig. 2). Furthermore, liquid chromatography mass spectrometry (LCMS) and NMR spectroscopy demonstrate that an identical final 4′, 5′ anhydroadenosine photoproduct is formed *in crystallo* and in solution^19^ upon illumination (Supplementary Figs. 3 and 4).

To capture light-induced structural-changes, time-resolved serial femtosecond crystallography (TR-SFX) experiments^24,25^ were carried out on the ns - ms time scale at the SPring-8 Angstrom Compact free electron LAser (SACLA)^26^ and SwissFEL^27^ X-ray free electron lasers (XFELs) using *Tt*CBD microcrystals (Supplementary Fig. 5). Guided by the spectroscopic kinetics, TR-SFX experiments were carried out according to an optical-laser pump - X-ray probe scheme at 10 ns, 300 ns, 3 µs, 100 µs, and 3 ms (SACLA) and 3 µs and 10 ms (SwissFEL). The fluence (30 mJ/cm^2^) of the nanosecond pump laser pulses (530 nm) was chosen based on crystallographic and spectroscopic power titrations (see Supplementary Note 2, Extended Data Figs. 3 and 4 and Supplementary Fig. 6) so as to maximize the photo-converted fraction while limiting the risk of non-linear effects^28^.

Structural changes were visualized in Fourier difference maps calculated between light and dark (no laser) data sets (see Supplementary Note 3; SACLA, Extended Data Fig. 5 and Supplementary Figs. 7 and 8; SwissFEL, Supplementary Figs. 9-11). Intermediate-state structures were determined following structure factor amplitude extrapolation (see Supplementary Note 3, Supplementary Figs. 12-18) and were used as a basis for QM/MM simulations to determine the likely corresponding chemical species, while Density Functional Theory (DFT) cluster models were used to explore the chemical transformations between the cobalamin species along the reaction profile. Cryogenic UV-vis absorption measurements in vitrified solution (Supplementary Fig. 19) guided crystallography experiments on macrocrystals at the European Synchrotron Radiation Facility (ESRF) that allowed cryo-trapping of an intermediate-state structurally similar to that captured by TR-SFX at 3 µs (Supplementary Figs. 20 and 21), and whose Co redox state was assessed by cryo-trapping EPR spectroscopy in solution (Supplementary Fig. 22). To corroborate the biological relevance of the intermediates captured by TR-SFX and inform on large-scale structural changes leading up to tetramer dissociation, time-resolved X-ray solution scattering (TR-XSS) experiments^29^ were carried out on the µs to s time scale according to an optical-laser pump (530 nm) - X-ray probe scheme at the ESRF synchrotron. TR-XSS experiments were also carried out on full-length CarH and in the presence of DNA. Our integrated experimental and computational approach yields a holistic understanding of the CarH photoreaction mechanism across a hierarchy of timescales, revealing how fast photochemical changes drive protein structural changes via a unique rearrangement of the adenosyl group leading to the biological output (Fig. 1).

**Fig. 1:**
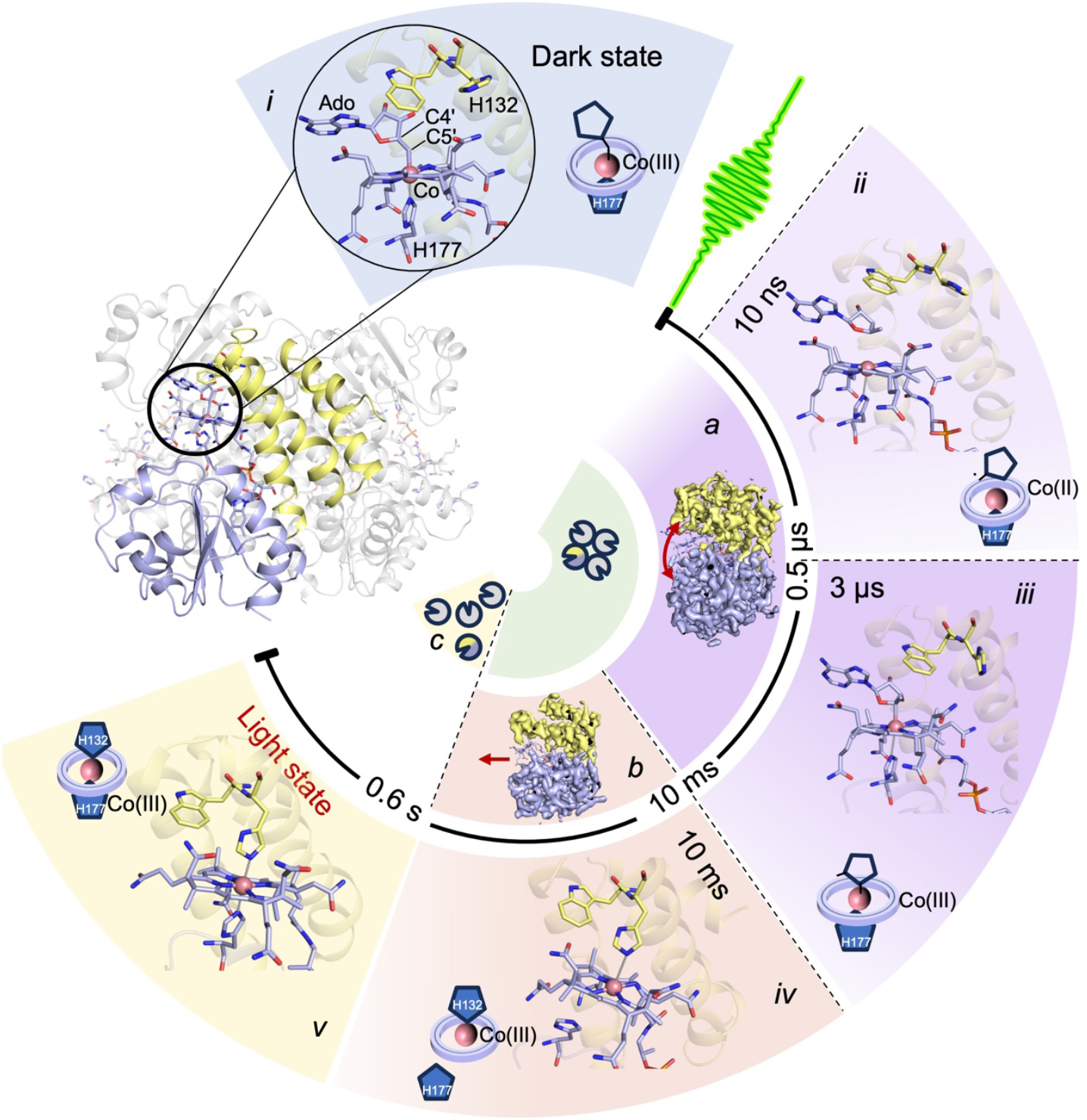
Spatio-temporal photoactivation mechanism of CarH based on integrated application of structural, spectroscopic, and computational methods. In the tetrameric dark-state of *Tt*CBD, each monomer comprising a Rossmann fold (light blue) and a four-helix bundle (yellow) domain binds an adenosylcobalamin (AdoCbl) chromophore with the adenosyl group coordinating the Co atom within the cobalamin corrin ring through its C5′ atom (*i*). Following green laser photoexcitation, sequential photointermediates (*ii - iv*) are transiently populated to yield the monomeric light-adapted state (*v*^5^). These include a bi-radical state with a broken Co–C5′ bond (*ii*), a Co–C4′ adduct (*iii*) and mono-histidine state (*iv*^21^). Monomer B is featured in *i* - *v* (only monomer B is considered hereafter; see Supplementary Note 3 for inspection of all monomers). Tertiary and quaternary structural changes, emanating from active site changes, feature cleft opening (*a*), corrin ring shifting (*b*) and finally tetramer dissociation (*c*).

### Diradical formation after Co–C5′ bond photolysis

In the earliest photointermediate captured by TR-SFX (10 ns, Fig. 1ii), the Co–C5′ bond is broken as evidenced in a Fourier difference electron density map (F_o_^10ns_30mJ/cm2_SACLA^ - F_o_^dark_ref_SACLA^, Fig. 2a) calculated at 2.3 Å resolution and in the 10 ns intermediate-state structure (orange model in Fig. 2b; Co–C5′ distance of 4.5 ± 0.1 Å in monomer B). Further structural changes in the active site with respect to the dark-state structure (Fig. 2b) include a shift in the W131 side chain and a tilt in the ribose moiety of the adenosyl group with its O3′ atom shifting by 1.7 ± 0.1 Å. Further structural changes can be identified from a distance difference matrix calculated between the 10 ns intermediate-state model (*10ns_30mJ/cm2_SACLA*) and the dark-state structure (*dark_ref_SACLA*; Fig. 2d). This reveals part of the four-helix bundle moving away from part of the Rossmann fold domain in a clamshell-like motion (inset in Fig. 2d, Fig. 1a), a structural change that persists at all TR-SFX time delays up to 3 ms (Extended Data Fig. 6). Our TR-SFX evidence indicates that the Co–C5′ bond is cleaved on a timescale faster than 10 ns, which contradicts the suggestion by Miller *et al*^23^ that this process occurs on a µs timescale, simultaneously with electronically excited triplet-state decay.

**Fig. 2:**
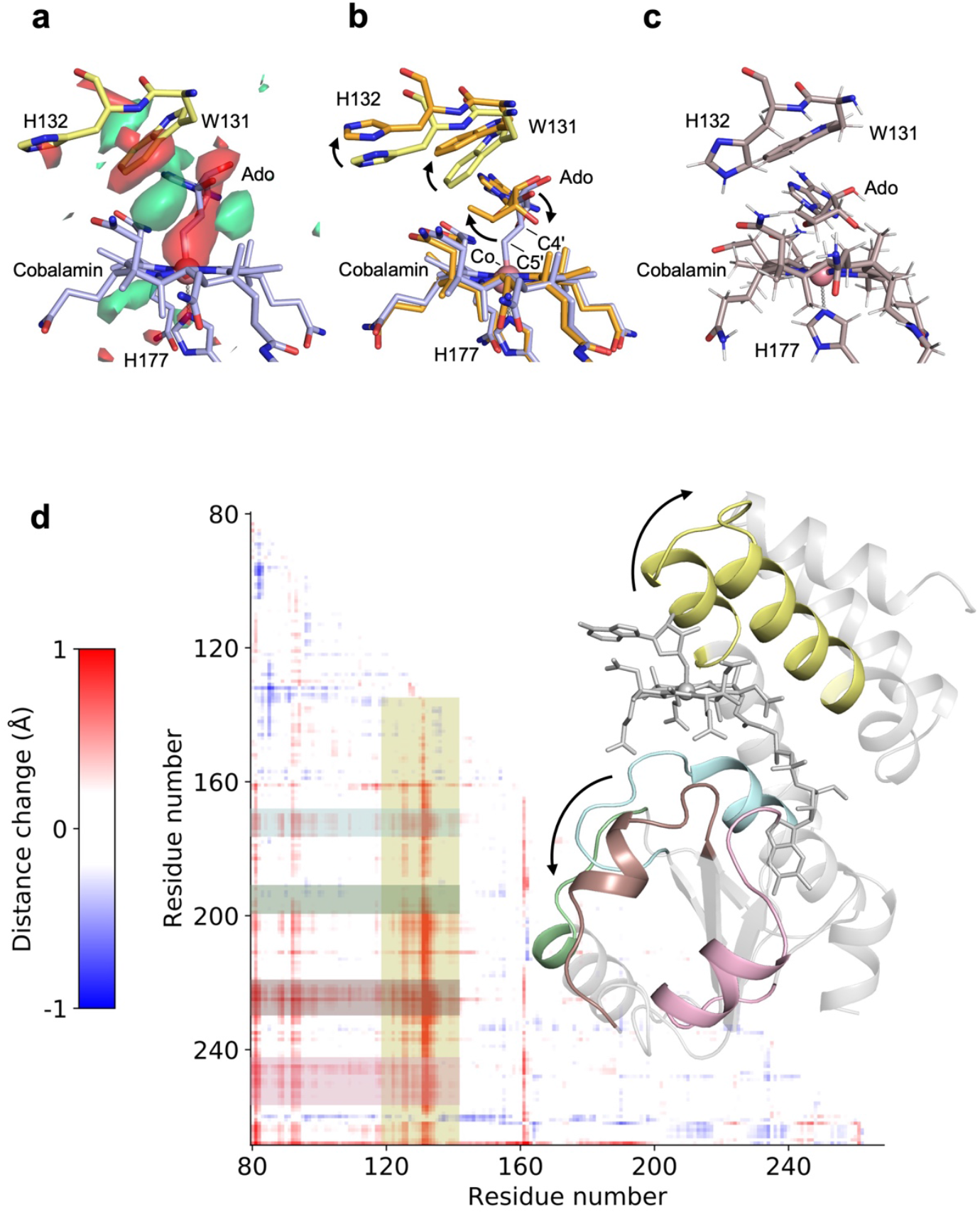
Co–C5′ bond photolysis leads to radical formation. (***a***) Time-resolved crystallography (TR-SFX) demonstrates Co–C5′ bond photolysis having occurred 10 ns after photon absorption as evidenced in a Fourier difference electron density map (Fo^10ns_30mJ/cm2_SACLA^ - Fo^dark_ref_SACLA^) of monomer B, computed between SFX data collected 10 ns after pump laser (530 nm) excitation and without excitation, respectively, contoured at -3.5 σ (red) and +3.5 σ (green) and displayed for the chromophore-binding pocket. The dark-state structure (*dark_ref_SACLA*) is superimposed in yellow (protein) and blue (chromophore) sticks. The map drawn over the entire tetramer is shown in Extended Data Fig. 5. (***b***) Overlay of the dark-state structure (*dark_ref_SACLA*) and the 10-ns intermediate-state structure (*10ns_30mJ/cm2_SACLA*) refined against extrapolated structure factor amplitudes (monomer B), with the protein in yellow and orange sticks, respectively and the chromophore in blue and orange sticks, respectively. The Co–C5′ distance increases from 2.0 ± 0.1 Å in the dark-state structure to 4.5 ± 0.1 Å in the 10-ns intermediate-state structure (errors estimated by the bootstrap technique^31^, see materials-and-methods section in SI). (***c***) Singlet diradical structure, based on the 10-ns intermediate-state structure and optimized by QM/MM. (***d***) Distance difference matrix (DDM) calculated between the Cα atoms of the 10 ns intermediate-state and the dark-state structures of monomer B. Red and blue pixels indicate intra-monomer distances becoming larger and shorter in the 10 ns intermediate-state compared to the dark-state structure, respectively. The DDM information content is transcribed structurally in the inset, with the distances between part of the four-helix bundle (yellow, residues L121 to T146) and part of the helices in the Rossmann fold domain (T171 - to L183, light blue; P200 - to D206, light green; A223 - to D234, brown; Q249 - to E262, light magenta) increasing in 10-ns intermediate-state structure, representing the clamshell-like movement represented by a curved arrow in Fig. 1a.

We investigated the chemical nature of the 10-ns intermediate-state structure by the QM/MM method, following the same protocol as previously applied by Toda *et al*^30^. A model corresponding to a diradical species with a Co(II)/C5′• radical pair (Fig. 2c) was more consistent with the experimental 10-ns intermediate-state structure (Fig. 2b, orange model) than a photoproduct model (hydridocobalamin and 4′,5′-anhydroadenosine, see Supplementary Note 6 for details), as judged from key distances, i.e. Co–C5′, Co–C4′ and Co–NH177 (Supplementary Table 1). Therefore, we assign the experimental 10 ns intermediate-state structure to a diradical, indicating that Co–C5′ bond photolysis is homolytic in nature, as proposed earlier^19,23^.

### Formation of an unexpected transient Co–C4′ adduct

Following Co–C5′ bond breakage the next intermediate in the photoactivation process is formed with a spectroscopic time constant of ∼500 ns (Fig. 1iii; Extended Data Fig. 2). This intermediate is characterized by further structural changes as evidenced by TR-SFX experiments with a 3 µs time delay between pump and probe pulses (Fig. 1iii). The Fourier difference electron density map (F_o_^3μs_30mJ/cm2_SACLA^ - F_o_^dark_ref_SACLA^, Fig. 3a, calculated at 2.25 Å resolution) and the 3 µs intermediate-state structure (orange model in Fig. 3b) indicate re-binding of the adenosyl group to the Co, surprisingly via the C4′ atom (Co–C4′ distance of 2.2

**Fig. 3:**
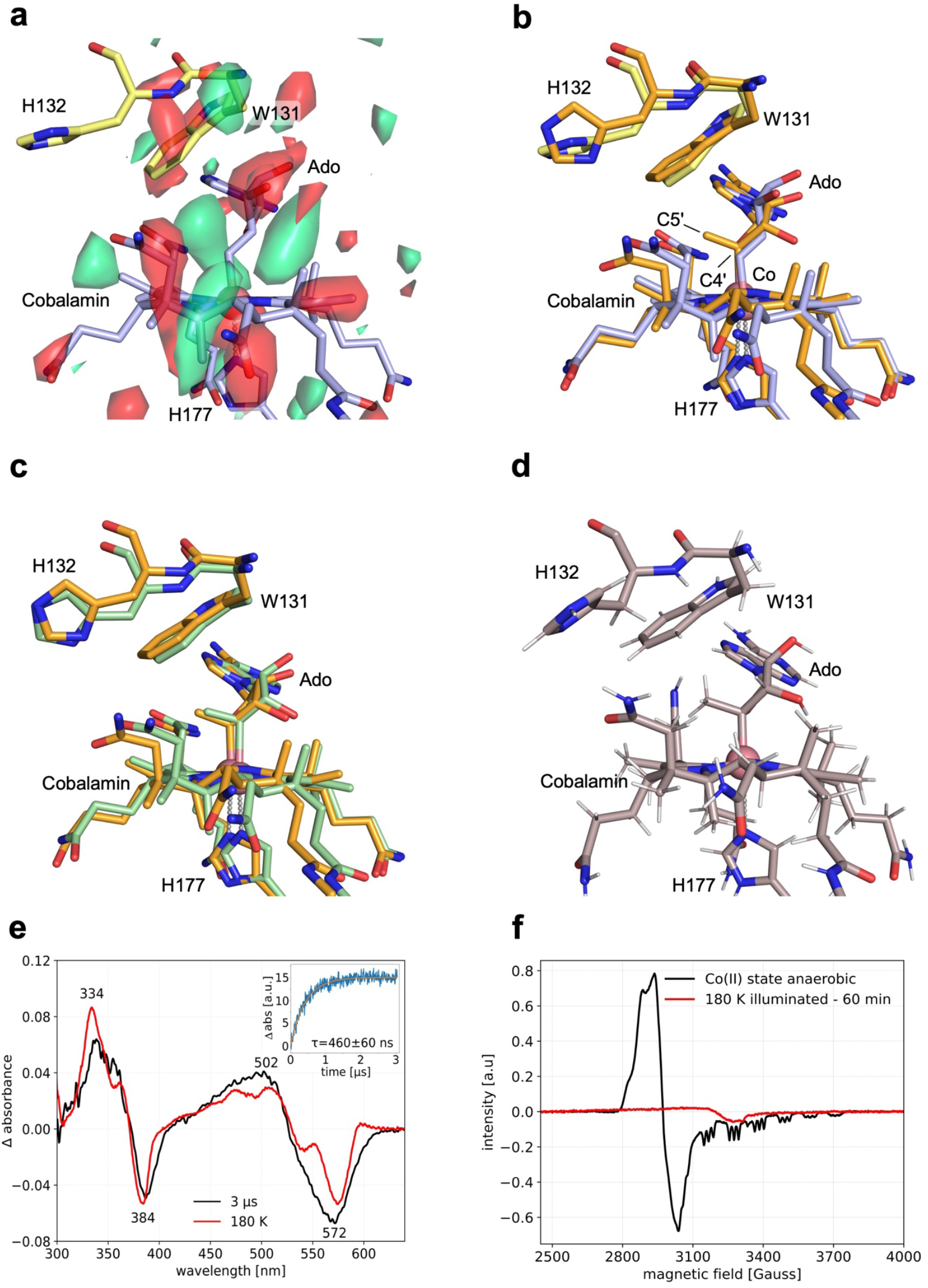
Co–C4′ adduct formed transiently. (***a***,***b***) Structural changes in the chromophore-binding pocket having occurred 3 µs after photon absorption as observed by TR-SFX. (***a***) Fourier difference electron density map (Fo^3μs_30mJ/cm2_SACLA^ - Fo^dark_ref_SACLA^) of monomer B, computed between SFX data collected 3 µs after pump laser (530 nm) excitation and without excitation, respectively, contoured at - 3.5 σ (red) and + 3.5 σ (green) and displayed for the chromophore-binding pocket. The dark-state structure (*dark_ref_SACLA*) is superimposed in yellow (protein) and blue (chromophore) sticks. The map drawn over the entire tetramer is shown in Extended Data Fig. 5. (***b***) Overlay of the dark-state structure (*dark_ref_SACLA*) and the 3 µs intermediate-state structure (*3µs_30mJ/cm2_SACLA*) refined against extrapolated structure factor amplitudes (monomer B), with the protein in yellow and orange sticks, respectively and the chromophore in blue and orange sticks, respectively. (***c***) Comparison of structural models obtained by cryo-trapping *Tt*CBD crystals after illumination at 180 K (green) and by TR-SFX 3 µs after photon absorption (orange). (***d***) Co–C4′ adduct structure, based on the 3 µs intermediate-state structure and optimized by QM/MM. (***e***) Time- and temperature-resolved difference absorption spectra of *Tt*CBD in solution (50 µM). Black: spectrum measured 3 µs after pump laser (530 nm) excitation. Red: spectrum measured at 77 K after illumination (530 nm LED, 15 mins) at 180 K. The inset shows a kinetic transient at 500 nm over 3 µs upon excitation with the pump laser. Data were fitted to a single exponential equation (orange line) to obtain time constants. (***f***) EPR silent spectrum of *Tt*CBD in solution recorded at 20 K, after steady state illumination (530 nm) at 180 K (red). As a reference, the signal of a Co(II) state is shown in black.

± 0.1 Å, monomer B). This experimentally determined structure was optimized in a QM/MM approach according to different chemical species (Extended Data Fig. 7, Supplementary Note 6), among which a covalent Co–C4′ adduct (Fig. 3d) whose Co–C4′ distance of 2.1 Å matches the experimentally determined value (Supplementary Table 2). Furthermore, in the QM/MM-optimized adduct (Fig. 3d) there was a 0.7 Å elongation of the Co–H177NE2 distance which aligns with the 3 µs intermediate-state structure (2.9 and 2.6 ± 0.1 Å, respectively, compared with 2.3 ± 0.1 Å in the dark-adapted monomer B). The experimental 3 µs intermediate-state is thus likely to represent a covalent Co–C4′ adduct. Prior to the TR-SFX experiments, a comparable Co–C4′ adduct was identified in a H132A mutant by serial synchrotron crystallography after laser excitation at Diamond Light Source (Supplementary Note 4).

A structure similar to the 3 µs intermediate-state structure was obtained by temperature-controlled synchrotron cryo-crystallography^32,33^ (see quantitative comparison in Supplementary Note 5). A *Tt*CBD macrocrystal was illuminated at 180 K (Fig. 3c) until no more spectral changes could be observed by microspectrophotometry^34^ (6 min; Supplementary Fig. 20). The crystal was then cooled to 100 K and X-ray diffraction data collected. The similarity between the UV-vis difference absorption spectra measured on *Tt*CBD in solution 3 µs after pump laser excitation and during steady state illumination of a crystal at 180 K (Fig. 3e) suggests that the same reaction intermediate is formed at 3 µs and 180 K. Therefore, the redox state of the Co atom of the 3 µs intermediate-state structure can be inferred by cryo-trapping EPR measurements. EPR spectra of a flash-cooled *Tt*CBD solution following illumination at 180 K reveals that no Co(II) species are present at this stage of the reaction (Fig. 3f). As the UV-vis absorbance features (red in Fig. 3e) do not match those of a Co(I) species^21^ it is most likely that the intermediate-state present at 180 K and at 3 µs represents a Co(III)– C4′ adduct.

To the best of our knowledge, the Co–C4′ adduct has not been observed previously in B_12_-dependent systems. Hence, the ability of CarH to form the Co–C4′ ligated species distinguishes it from thermally-activated B_12_ enzymes and is linked to its unique ability to harness the inherent photoreactivity of AdoCbl. Importantly, our DFT cluster calculations (see Supplementary Note 6) suggest that, after photochemical dissociation, formation of the Co– C4′ adduct is kinetically favoured over a return to the dark state (barriers of 6.9 and 14 kcal mol^-1^, respectively; energies and structural parameters are shown in Supplementary Table 3). The DFT calculations (Supplementary Fig. 23) show a significantly lower bond dissociation energy in the Co–C4′ adduct (∼17 kcal mol^-1^ *vs* ∼35 kcal mol^-1^ in the Co–C5′ adduct), which suggests that Co–C4′ bond breakage may occur thermally. This is not surprising since breaking the Co–C5′ bond forms a primary radical while breaking the Co–C4′ bond forms a tertiary radical. Taken together, the data illustrate how the formation of the Co–C4′ adduct may protect the system from returning to the dark state before the adenosyl group has been able to escape the binding pocket. This proposed mechanism, along with a computed potential energy profile, is shown in Supplementary Fig. 23.

### Exit of adenosyl group, corrin ring displacement and Co ligation switch

The putative Co–C4′ adduct species decays into a new intermediate-state with a spectroscopic time constant of ∼7 ms (Extended Data Fig. 2), in an O_2_-dependent process^21^. Structural changes that accompany the decay of the Co–C4′ adduct have been observed on the millisecond time scale by TR-SFX at the SwissFEL (Fig. 4). At 10 ms the Fourier difference electron density map (F_o_^10ms_30mJ/cm2_SwissFEL^ - F_o_^dark_ref_SwissFEL^, Fig. 4a, calculated at 2.9 Å resolution) points towards extensive changes at both the adenosyl and the cobalamin groups and at H132. Indeed, the 10-ms intermediate-state structure of monomer B, refined against extrapolated structure factor amplitudes (orange model in Fig. 4b), lacks the adenosyl group and features a corrin ring shifted by 4.3 ± 0.2 Å, with the Co now ligated in the upper position by H132 rather than the original, lower H177 coordination. The clamshell-like opening of the chromophore-binding pocket, observed between 10 ns and 3 ms (Fig. 2d), is reversed at 10 ms (Extended Data Fig. 6). The 10 ms intermediate-state structure is almost identical [root mean square deviation (RMSD) protein Cα and cobalamin atoms in monomer B of 0.6 and 1.1 Å, respectively] to the one solved by synchrotron cryo-crystallography after illuminating a *Tt*CBD macrocrystal for 5 s at RT followed by flash-cooling^21^; Fig. 4c; see Supplementary Note 3). The same spectroscopic species forming with a time constant of ∼7 ms has also been identified by cryogenic UV-vis absorbance spectra collected on flash-cooled *Tt*CBD solutions illuminated at 180 K before progressive warming to higher temperatures (230-260 K) in the dark (Supplementary Fig. 19). Cryogenic UV-vis spectra (Supplementary Fig. 24) suggest that this intermediate represents a water/OH–ligated Co(III)–H132 species (see Supplementary Note 5). Hence, we conclude that the Co–C4’ adduct decays to a new state that is characterized by a shifted corrin ring structure with a Co(III) atom ligated to a single histidine residue (H132) at the upper position.

**Fig. 4:**
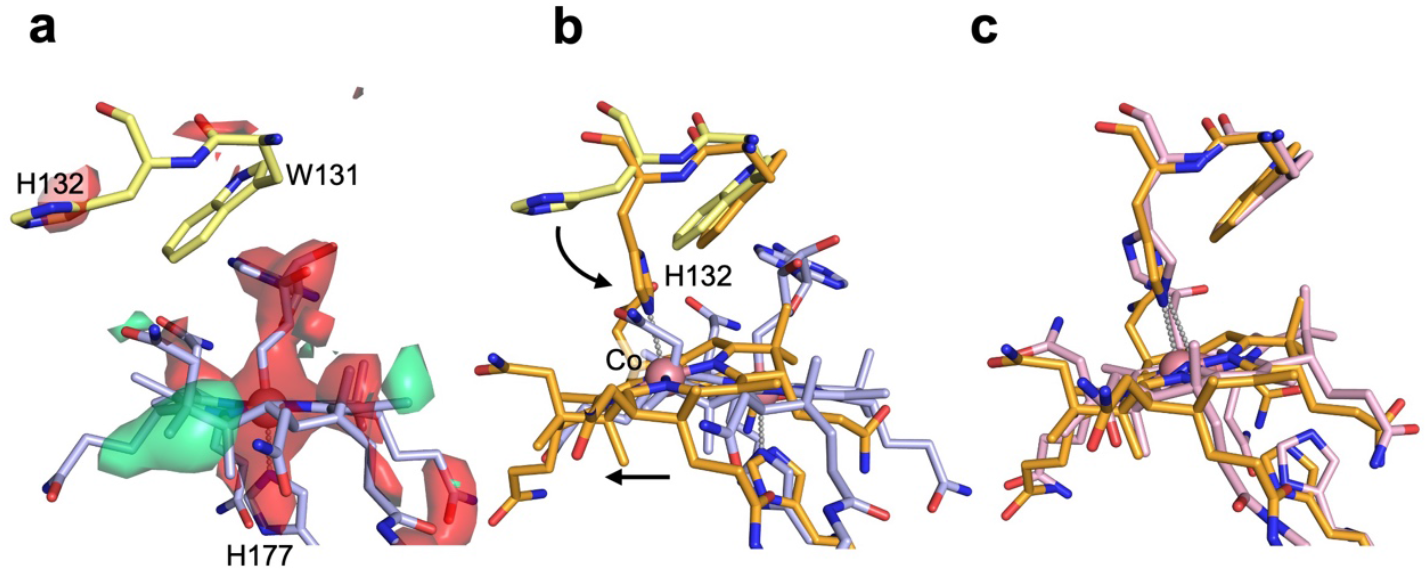
Exit of adenosyl moiety is followed by corrin ring displacement and Co ligation switch. (***a***) Fourier difference electron density map (Fo^10ms_30mJ/cm2_SwissFEL^ - Fo^dark_ref_SwissFEL^) of monomer B, computed between SwissFEL SFX data collected at 10 ms after pump laser (530 nm) excitation and without excitation, respectively, contoured at - 3.5 σ (red) and + 3.5 σ (green) and displayed for the chromophore-binding pocket. The dark-state structure (*dark_ref_SwissFEL*) is superimposed. The map drawn over the entire tetramer is shown in Supplementary Fig. 11. (*B, C*) Overlay of the 10 ms intermediate-state structure (*10ms_30mJ/cm2_SwissFEL*) refined against extrapolated structure factor amplitudes (monomer B) and (***b***) the dark-state structure (*dark_ref_SwissFEL*) and (***c***) a synchrotron cryo-crystallography structure of *Tt*CBD obtained after illuminating a macrocrystal for 5 s at RT followed by flash-cooling (PDB entry code: 8C76)^21^. In the 10 ms intermediate-state structure, the dark-state structure and the synchrotron cryo-crystallography structure, the protein is in orange, yellow and pink sticks, respectively and the chromophore in orange, blue and pink sticks, respectively.

### Large scale movements leading to tetramer dissociation and light-adapted state formation

Structural changes leading to the monomeric light-adapted state were measured in solution by using TR-XSS experiments covering the µs-to-s time scale (Fig. 5a). Data from the short time delays (≤ 100 ms) can be entirely reconstructed as a linear combination of only two species (Fig. 5a), with a time constant for the transition between these of 10.0 ± 0.4 ms (Fig. 5e), as revealed by a Singular Value Decomposition (SVD) analysis^35^ (Extended Data Fig. 8). The experimental 100 µs difference signal (representative of the first species) is in good agreement up to ∼0.9 Å^-1^ (see peaks at ∼0.2, 0.3, 0.6 and 0.8 Å^-1^) with the difference signals calculated using the atomic coordinates of all the TR-SFX intermediate-state structures up to 3 ms (Fig. 5d). The experimental 100 ms difference signal (representative of the second species) is in excellent agreement with the difference signal (Fig. 5c) calculated using the atomic coordinates of the corrin-ring-shifted *Tt*CBD (PDB entry code: 8C76 and 10 ms intermediate-state, Fig. 4c and Supplementary Fig. 25). Importantly, a difference signal calculated based on a modified dark-state *Tt*CBD (8C73^*^), generated using the dark-state protein scaffold (PDB entry code: 8C73) but with the cobalamin atomic coordinates from the corrin-ring-shifted *Tt*CBD structure (PDB entry code: 8C76), is also largely compatible with the 100 ms TR-XSS data (Fig. 5c). Therefore, most of the TR-XSS difference signal originates from the corrin ring shift (Fig. 4b). Figs 5C-E show that: (i) the clamshell-like structural change revealed by SFX (Fig. 2d) occurs also in solution, without additional structural changes that would be inhibited in crystals; (ii) the clamshell-like change gives way to the corrin ring shift with a 10-ms time constant, in agreement with TR UV-Vis kinetics (green trace in Fig. 5f).

**Fig. 5:**
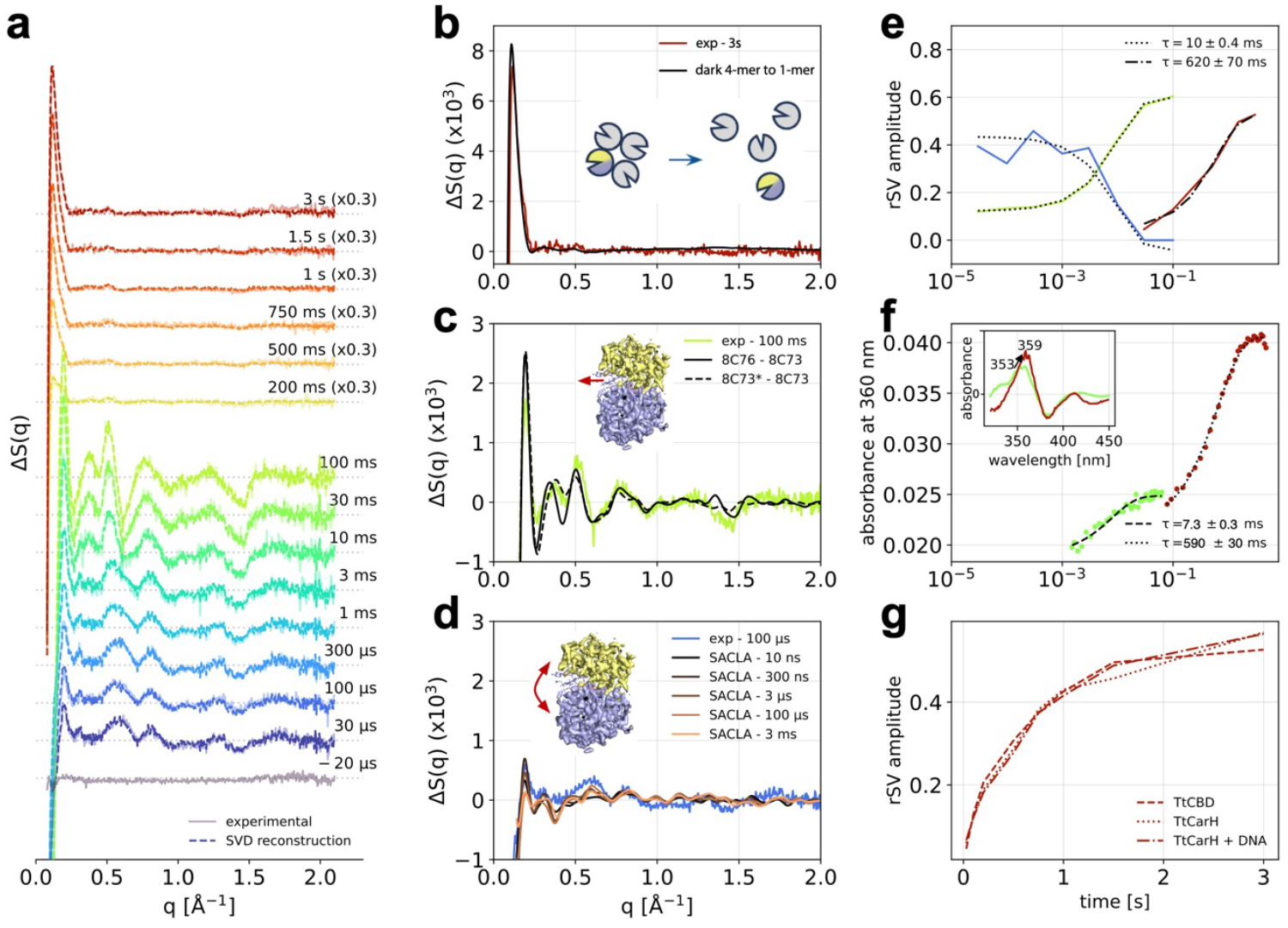
Photoinduced intermediates of *Tt*CBD in solution eventually lead to tetramer dissociation. (**a**) Time-resolved X-ray solution scattering signals of *Tt*CBD in solution from 30 µs to 3 s after removal of the solvent heating signal (Supplementary Fig. 29) and correction for the photoconversion accumulation (Supplementary Fig. 30) (coloured continuous lines) overlaid on their SVD reconstructions (coloured dashed lines). The protein concentration in the 200 ms – 3 s range was reduced 4 × to reach a CO/dissolved-O_2_ ratio that allowed for the photoreaction to be completed (see materials-and-methods section in SI). The intensity of signals in the 200 ms - 3 s range is multiplied by 0.3. (**b**) Comparison between the 3 s TR-XSS signal (red line) and the calculated difference obtained using the tetrameric dark structure (4-mer, PDB entry code: 8C73) and a monomer (1-mer) (black line). (**c**) Comparison between the 100 ms TR-XSS signal and calculated differences obtained using the cryo-trapped corrin-ring-shifted structure reported by Poddar *et al*.^21^ (black continuous line) and a modified tetrameric dark-state structure (8C73^*^) obtained using the PDB 8C73 with the corrin ring coordinates from the cryo-trapped structure PDB 8C76 (black dashed line). (**d**) Comparison between the 100 µs experimental TR-XSS signal and calculated differences obtained using TR-SFX intermediate-state structures at 10 ns, 300 ns, 3 µs, 100 µs and 3 ms. The tetrameric dark structure determined by SFX (Fig. 1) was used as a reference. (**e**) Time course of the SVD right-singular vectors (rSV) amplitude from the short (≤ 100 ms; green, blue) and the longer time-delay TR-XSS data (> 100 ms; red). (**f**) Time course of the optical absorbance change at 360 nm with (inset) the spectral differential signal at 50 ms (green continuous line) and 5 s (red continuous line). (**g**) Time course of the SVD rSV amplitude from the ms-to-s TR-XSS data sets in *Tt*CBD, *Tt*CarH and *Tt*CarH-DNA.

TR-XSS data at longer time delays (> 100 ms) were collected at a Co/dissolved-O_2_ ratio that allows photoreaction completion^19,21^ (Supplementary Fig. 26) and reveal a new species characterized by an intense peak at ∼0.11 Å^-1^ (Fig. 5a). These data are described by a single basis pattern (Fig. 5a) building up with a time constant of 620 ± 70 ms (Fig. 5e), as determined by SVD analysis. The near perfect agreement between the TR-XSS signal at 3 s and the calculated signal expected for the *Tt*CBD tetramer-monomer transition (Fig. 5b) clearly assigns the signal change captured at 620 ms to the tetramer dissociation. This dissociation seems to occur in a single step (Fig. 5b, Supplementary Fig. 27). To confirm the biological relevance of the results obtained on *Tt*CBD, analogous TR-XSS data were collected on both the full-length *Tt*CarH and a *Tt*CarH-DNA complex. These data sets are in overall agreement with that on *Tt*CBD (Supplementary Fig. 28), showing comparable albeit slightly longer time constants (710 ± 50 and 760 ± 30 ms, respectively) (Fig. 5g and Supplementary Fig. 28). The time constant for tetramer dissociation in *Tt*CBD (620 ms) is very similar to the TR-UV-Vis spectral change of 590 ± 30 ms associated with formation of the bis-histidine ligation that characterizes the light-adapted state (Fig. 5f)^5^. Taken together, we therefore assign the build-up of the final *Tt*CBD light-adapted state (i.e. bis-histidine ligated)^5^ as concomitant with tetramer dissociation (Fig.1).

### CarH photoactivation mechanism

Our integrated experimental and computational approach combining time-resolved, temperature-resolved and quantum chemical calculations provides a comprehensive picture of the photoactivated reaction in the *Tt*CarH CBD (Fig. 1, Extended Data Fig. 9). Following photon absorption by the B_12_ chromophore, the light-sensitive bond connecting the Co atom of the cobalamin to the C5′ atom of the adenosyl group breaks homolytically, yielding a singlet diradical Co(II) species (Fig. 1ii), the 3D structure of which was captured at 10 ns (Fig. 1ii). The diradical species (Fig. 1ii) is then converted on the microsecond timescale to a previously unidentified adduct between Co and the C4′ atom of the adenosyl group (Fig. 1iii). The formation of the adduct is only likely to be possible due to the unique orientation of the H–C4′ bond with respect to the Co atom in CarH, as compared to thermally-activated B_12_ enzymes^36^, enabling Co–assisted H transfer from C4′ to C5′. Calculated bond dissociation energies indicate that the Co–C4′ bond can be broken thermally, unlike the initial Co–C5′ bond which requires photon absorption. We speculate that formation of the Co–C4′ adduct prevents reformation of the Co–C5′ bond that would return the diradical species to the initial dark-state. The Co–C4′ adduct then decays on the millisecond timescale to yield an intermediate in which the adenosyl group has disappeared from the binding site, the corrin ring has moved by ∼5 Å, and the Co ligation has switched from H177 to H132 (Fig. 1*iv*). The associated movement of H132 has been suggested to cause larger-scale motions beyond the chromophore-binding pocket that lead to tetramer dissociation^5^. These cannot be captured by time-resolved crystallography as the diffraction quality degrades at longer timescales, presumably because of the larger-scale movements being incompatible with crystal packing. Time-resolved X-ray solution scattering confirmed the formation of the corrin ring-shifted intermediate in solution on the millisecond timescale and established that it is followed by tetramer-to-monomer transition occurring in a single process with a time constant of ∼0.6 s (Fig. 1c) concomitant with formation of the final light-adapted state (e.g. bis-histidine cobalamin adduct, Fig. 1v^5^). Importantly for the biological relevance of the results presented here, similar kinetics of monomer formation is observed in the full-length *Tt*CarH, even when bound to DNA.

The elucidated photoactivation mechanism of the *Tt*CarH CBD spans nine orders of magnitude in time, leading from B_12_ photolysis to tetramer dissociation. Surprisingly few intermediates appear to be involved, which we suggest confers robustness towards energy dissipation that would otherwise abort the photoactivation process without biological output. A Co–C4′ adduct, not observed previously in thermally-activated B_12_ enzymes, is key to photoactivation, maintaining the photoreceptor on path from nano-to milliseconds, thus bridging the photochemical and photobiological timescales. Formation of this crucial intermediate is facilitated by the unique orientation of the B_12_ cofactor in CarH, highlighting potential routes for harnessing B_12_ photochemistry for future biotechnological innovations. Importantly, the discovered intermediates and their associated reaction pathways advances our understanding of how light-driven processes in B_12_-dependent photoreceptors differ from thermal pathways and provides crucial insights for the design and optimization of B_12_-dependent proteins in potential optogenetics and enzyme photocatalysis applications.

## Methods

### Protein expression, purification and crystallization

The expression and purification protocols for full-length CarH, *Tt*CBD and the variant *Tt*CBD-H132A were adapted from those previously published for full-length CarH^37^. Proteins with a C-terminal 6×His tag were expressed in *E. coli* BL21(DE3) cells. The cells were lysed by sonication, then centrifuged and the supernatant containing the apo*-*form of the expressed protein (i.e. full-length apo-CarH, apo-*Tt*CBD the variant apo-*Tt*CBD-H132A) was retained for purification. The crude extract was subjected to affinity chromatography and partially purified protein obtained after this step was incubated with adenosylcobalamin (AdoCbl) to generate holo-forms of the respective proteins in the dark. A final size exclusion chromatography step with 50 mM Tris-HCl pH 7.5, 100 mM NaCl as eluent was performed to achieve high purity and removal of free AdoCbl. Detailed protocols for protein expression and purification are provided in the Supplementary Methods.

For TR-SFX experiments at SACLA and SwissFEL, macrocrystals of *Tt*CBD were produced by the batch method, at room temperature (20 °C) and under red light conditions as described recently^21^. Those large crystals were used to generate seeds under red-light conditions according to a recently reported protocol^38^. Microcrystals were grown by the seeded batch method using 20% PEG 10000, 0.1 M HEPES pH 7.5. For SSX experiments at the Diamond Light Source, *Tt*CBD-H132A was microcrystallised in batches using a previously reported^21^ crystallization condition from LMB screen (Molecular Dimension) containing 0.1 M ammonium sulphate, 0.1 M sodium citrate, pH 5.8, 16% w/v PEG 4000, and 20% v/v glycerol. For MX experiments at the ESRF, macrocrystals of *Tt*CBD were obtained using a previously reported protocol^21^. Detailed crystallization procedures are provided in the Supplementary Methods.

### Time-resolved absorption spectroscopy

Time-resolved absorption spectroscopy experiments after laser photoexcitation were carried out on *Tt*CBD samples in solution using an LP980-KS flash photolysis instrument (Edinburgh Instruments Ltd). Photoactivation was initiated by ns laser excitation at 530 nm, using an optical parametric oscillator of a Q-switched Nd-YAG laser (NT432, EKSPLA) in a 1-cm pathlength cuvette. Laser pulses were between 6-8 ns in duration and varied in energy up to 20 mJ by using an attenuator. Difference absorbance spectra were recorded between 300 and 700 nm at selected time points using an image intensified CCD camera (Andor Technologies). Single-wavelength kinetic absorption transients were recorded at a range of wavelengths between 300 and 700 nm with the detection system (comprising probe light, sample, monochromator and photomultiplier) at right angles to the incident laser beam. Single-wavelength kinetic absorption transients were also collected on *Tt*CBD microcrystals embedded in sodium carboxymethyl cellulose (CMC, Sigma) in a 1-mm pathlength cuvette at a 45° angle to the incident laser beam and the probe beam. An extended version of time-resolved absorption spectroscopy methods are provided as Supplementary Methods.

### Temperature-resolved absorption spectroscopy

Static absorbance spectra were measured using a Cary 50 spectrophotometer (Agilent Technologies). For temperature-resolved absorption spectroscopy measurements, *Tt*CBD samples were cooled down to 77 K at a rate of ∼10 K per minute in an Optistat DN liquid nitrogen cryostat (Oxford Instruments Inc.) to record spectra. Samples were then warmed to the desired temperature at a rate of ∼10 K per minute to initiate the reaction by illumination (approx. 1000 µmol/m^2^ /s) with a 530 nm LED (Thorlabs Inc.) for 15 min, before cooling again to 77 K to record spectra. Room temperature absorbance spectra were also collected on *Tt*CBD microcrystals embedded in a sodium carboxymethyl cellulose (CMC) matrix in a 1-mm pathlength cuvette. An extended version of temperature-resolved absorption spectroscopy methods are provided as Supplementary Methods.

### Determination of photoproduct formed when illuminating *Tt*CBD crystals

Using the method utilized in Zhang *et al*^10^ we sought to establish whether the same photoproduct is formed in the *Tt*CBD microcrystals produced for TR-SFX. Microcrystals were either kept in the dark or placed in ambient light for 30 min and subsequently resuspended in deuterated methanol (Eurisotop) to be used for liquid chromatography mass spectrometry (LCMS) and for NMR elucidation.

LCMS was undertaken on an Agilent 1100 LC-MSD instrument with an Agilent 150 × 3.0 mm Poroshell 120 SB-C18 (2.7 µm pore size) reversed phase column. An in-line ESI-TOF single quadrupole mass spectrometer (Agilent Technologies) in positive ion mode was used to determine mass-to-charge ratios (m/z).

NMR spectra were recorded on a Bruker AVIII 500 MHz spectrophotometer with 1H/19F/13C-15N QCI-F cryoprobe equipped with z-gradients. Initially, all NMR spectra were recorded using the same parameters as in Zhang *et al*^10^. Briefly, 1D 1H NMR spectra were collected at 298 K using a 1D 1H NMR method with presaturation water suppression. More extended descriptions of LCMS and NMR methods used are given as Supplementary Methods.

### Low temperature cw-EPR

For electron paramagnetic resonance (EPR) spectroscopy, samples containing 200 μL of ∼ 250 μM *Tt*CBD were prepared under aerobic and anaerobic conditions. The anaerobic and aerobic samples were photoactivated with a Thorlabs Mounted LED (emitting at 530 nm with a nominal output power of 370 mW) at room temperature and at 180 K, respectively. Samples were measured on a Bruker EMXplus EPR spectrometer equipped with a Bruker ER 4112SHQ/ X-band resonator at 20 K (measurements at higher temperatures are stated accordingly). Data processing and analysis were performed using the EasySpin toolbox (5.2.36) for the Matlab program package^39^. Sample preparation and illumination protocols, as well as EPR data collection and processing details are specified in the Supplementary Information.

### TR-SFX data collection and processing and structure refinement

TR-SFX experiments involving nanosecond pump-laser excitation were carried out at SACLA^26^ and SwissFEL^27^ using *Tt*CBD microcrystals whose size (∼ 15 × 5 × 2 μm (Supplementary Fig. 5) did not exceed in any dimension the 1/e penetration depth at 530 nm (17 μm; Supplementary Fig. 40) to ensure efficient photoexcitation at low pump-laser fluences. Spectroscopic and crystallographic pump-laser power titrations were carried out to choose the appropriate fluence (Supplementary Note 2).

For data collection at SACLA, *Tt*CBD microcrystals were embedded in CMC (detailed protocol provided as Supplementary Information) and extruded into a helium-filled *Diverse Application Platform for hard X-ray Diffraction in SACLA* (DAPHNIS)^40^ chamber by a high-viscosity extrusion (HVE)^41^ injector^42^. The SFX experiments were carried out in a time-resolved mode (TR-SFX)^24^ according to an optical pump – X-ray probe scheme, using XFEL pulses at a repetition rate of 30 Hz (< 10 fs in length, nominal photon energy 7.996 keV (FWHM 42 eV), photon flux of ∼2 × 10^11^ photons/pulse, pulse energy of ∼300 μJ at the sample position) focused to 1.6 μm (h) × 1.4 μm (v) (FWHM). The pump laser (EKSPLA NT230, 530 nm, 5 ns pulse length (FWHM), circularly polarized) was aligned perpendicular to both the X-ray beam and the HVE jet and focused to a Gaussian spot with size of 105.0 (h) × 98.2 (v) µm^2^ (FWHM). TR-SFX data were collected at pump-probe delays of 10 ns, 300 ns, 3 μs, 100 μs, and 3 ms and at 3.5 μJ laser pulse energy (fluence at the Gaussian peak of 30 mJ/cm^2^ ; power density at the Gaussian peak of 6 MW/cm^2^ ; nominally 2.4 absorbed photons on average per chromophore). Diffraction data with (*light*) and without (*dark*) pump-laser excitation were collected in an interleaved way.

For data collection at the Cristallina experiment station^43^ of SwissFEL, *Tt*CBD microcrystals were mounted on micro-structured polymer fixed targets (MISP chips^43^, 30 × 30 mm^2^, detailed protocol provided as Supplementary Information). Interleaved *light-dark* TR-SFX data were collected using X-ray pulses at a repetition rate of 100 Hz (25 - 45 fs in length, mean photon energy 11 keV, photon flux of 1.59 × 10^11^ photons/pulse, pulse energy of ∼280 μJ at the sample position) focused to 3.7 μm (h) × 4.2 μm (v) (FWHM). The pump laser (EKSPLA NT230, 532 nm, 3.5 ns pulse length (FWHM)) was fibre-coupled collinear to the X-ray beam and perpendicular to the chip plane. The 1/e^2^ width of the laser spot at the sample position was determined by a beam profiler (55 µm) and assumed to be the diameter of a top-hat circular beam. TR-SFX data were collected at pump-probe delays of 3 μs and 10 ms and at 0.75 μJ laser pulse energy (30 mJ/cm^2^; 9 MW/cm^2^; nominally 2.5 absorbed photons on average per chromophore). A more detailed description of TR-SFX data collections at SACLA and SwissFEL is given as Supplementary Information.

Offline data processing was performed using CrystFEL^44^ versions 0.10.1 (SACLA) and 0.10.2 (SwissFEL). For *dark_ref*_*SACLA* structure refinement, the previously reported *Tt*CBD tetramer structure solved from form II crystals (PDB entry code: 8C73^21^) was used as a search model for molecular replacement with *PHASER*^45^. Iterative cycles of reciprocal space refinement performed using *REFMAC5*^*46*^ from the CCP4 software suite ^47^ were interspersed with local real-space refinements and model building using *COOT*^48^. *Dark_ref*_*SwissFEL* structure refinement was carried out similarly, with the refined *dark_ref_SACLA* model as a search model for molecular replacement. Q-weighted Fourier difference electron density maps (F_o_^*light*^ – F_o_^*dark*^)^49^ were computed and scalar structure factor extrapolation^50^ was carried out with *Xtrapol8* ^51^ to determine the structures of intermediate states at all time delays from experiments at both SACLA and SwissFEL. Models of all intermediate-state structures were refined against q-weighted extrapolated structure factor amplitudes using *REFMAC5* and *phenix*.*refine*^52^. Coordinate errors were estimated using the bootstrapping approach^31^. Detailed descriptions of data processing, refinement and error estimation are available as Supplementary Information. Data collection and refinement statistics are presented in Supplementary Tables 4-6.

### SSX data collection and processing and structure refinement

Serial synchrotron crystallography (SSX) after laser excitation was performed on the *Tt*CBD-H132A mutant at beamline I24 of Diamond Light Source (Didcot, UK)^53^. Microcrystals of the *Tt*CBD H132A mutant (∼15 × 8 × 5 μm^3^) were mounted on a glow-discharged silicon chip (Southampton Nanofabrication Centre, University of Southampton) with aperture size of ∼10-12 µm (detailed protocol provided as Supplementary Information). Two datasets were collected at room-temperature, i) without optical-pump (*dark_DLS*) and ii) according to an optical pump – X-ray probe scheme. Using an X-ray beam of 12.4 keV focused to 8 μm (h) × 8 μm (v) (FWHM), each crystal was exposed for 10 ms with a photon-flux of 1.6 × 10^12^ photons/second and diffraction data were recorded. For the SSX data collection involving illumination, a portable pulsed laser system from Light Conversion (PORTO) was used to photoactivate the *Tt*CBD H132A microcrystals. Mirrors and an achromatic focusing lens were used to focus the laser beam at the sample position, in such a geometry that the laser beam was ∼15° off-axis from the X-ray beam with a laser beam size of 50 µm (FWHM) in both directions. The laser was operated at a wavelength of 515 nm and a repetition rate of 50 kHz, providing individual pulses of 300 fs. The laser intensity at the sample position was attenuated using a neutral density filter, yielding a laser power of ∼0.8 mW; i.e. 0.016 μJ per 300-fs pulse. With this setup, pump laser excitation was performed during 10 ms (i.e. 500 pulses), corresponding to a pump laser energy of 8 µJ, a fluence of 320 mJ/cm^2^ and to nominally ∼12 absorbed photons per chromophore assuming an average crystal size of 8 μm. Diffraction images were collected during 10 ms immediately after laser exposure. This dataset was collected with the pump-laser on for all apertures of the chip, i.e. no *dark-interleaved* was collected. This dataset is referred to as *illuminated_DLS*.

For *dark_DLS* structure refinement, molecular replacement was performed with *PHASER*^*45*^ using as a search model the structure of the dark-adapted *Tt*CBD-H132A mutant (PDB entry code: 8C35), followed by a refinement procedure similar to the one described above for *dark_ref_SACLA* structure refinement. Refinement of an *illuminated_DLS* model against extrapolated structure factor amplitudes was not possible, as Fourier difference maps were uninterpretable due to an unexpectedly high R_iso_ value (19.3%). Instead, composite-model refinement was carried out as described in the Supplementary Information. Data collection and refinement statistics are presented in Supplementary Table 7.

### Cryo-temperature dependent macromolecular crystallography and in crystallo UV-vis spectroscopy

Cryo-temperature dependent macromolecular crystallography (MX) in combination with *in crystallo* UV-vis absorption microspectrophotometry^34^ on the BM07-FIP2 beamline of the ESRF^54^ (Supplementary Fig. 37) was used to characterize structural and spectroscopic changes in crystalline *Tt*CBD after light illumination at various cryo-temperatures, including at 180 K. Details on the illumination and temperature-cycling protocols, as well as on X-ray data collection and processing and structure refinement are given as Supplementary Information. Data collection and refinement statistics are reported in Supplementary Table 8.

### TR-XSS: data collection and reduction

TR-XSS data were collected at the ID09 beamline of the ESRF^55,56^. The protein solutions (*Tt*CBD, *Tt*CarH and a *Tt*CarH-DNA complex) were photoexcited with a 5 ns pulsed laser (EKSPLA NT342B) at 532 nm. The laser was focused with cylindrical lenses to an elliptical spot approximately 2.0 × 0.2 mm^2^ (FWHM) corresponding to a fluence of ∼50 mJ/cm^2^ and ∼2.5 absorbed photons/chromophore. X-rays (14.7 keV) probed the sample at 90° with respect to the laser incoming direction. Scattering patterns were recorded on a Rayonix MX170-HS and azimuthally integrated to obtain 1-dimensional scattering signals S(q). TR-XSS difference signals ΔS(q) were obtained as the difference between the signals measured after photoexcitation minus the signal measured before photoexcitation. Data reduction was performed using the txs python package developed at ID09 (https://gitlab.esrf.fr/levantin/txs). Two different TR-XSS datasets were collected on *Tt*CBD: the first dataset (30 µs - 100 ms) was obtained by flowing a high concentration sample through a quartz capillary in a closed loop with a peristaltic pump (Gilson, Minipuls); the second dataset (100 ms - 3 s) was obtained by flowing a low concentration sample through a quartz capillary with a syringe pump (Chemyx, Fusion 4000X). The lower concentration in the second dataset ensured a protein chromophore concentration approximately half of the oxygen concentration in solution, with fully aerobic conditions allowing completion of the photoreaction up to tetramer dissociation^19,21^. A protocol analogous to that used for the second dataset was also used to collect data on *Tt*CarH and a *Tt*CarH-DNA complex in the time interval 100 ms - 3 s. An extended description of TR-XSS methods is given as Supplementary Information.

### Cluster model calculations

We used Density Functional Theory (DFT) cluster models to examine the mechanism for the formation of a Co–C4′ adduct and explore the inherent differences in the chemistry of the Co– C5′ and Co–C4′ adducts. These calculations were performed in Gaussian16 rev. D^57^ using the BP86 functional^58^ which has been shown to perform well for B_12_ systems^59,60^, with the Def2-TZVP basis set on cobalt and the 6-31G(d,p) basis sets on the remaining atoms, an implicit solvation model (CPCM) with dielectric constant of 5.0 and empirical dispersion (GD3BJ^61,62^). The cluster model was constructed from the *dark_DLS* model of the H132A mutant (monomer A) and consisted of a truncated Cbl, adenosyl, and binding-site residues W131 and H177 truncated as –CH_3_ at the Cβ and V138, E141 and H142 truncated as CHO and NH_2_ C– and N– termini, respectively (Supplementary Fig. 38) for a total of 211 atoms, with a net charge of 0. Six atoms were kept fixed during the calculations: the Cβ of W131 and H177, the Cα of V138, E141 and H142 and the Co. Comparison with the WT crystal structure (*dark_ref_SACLA*) revealed that the fixed atoms could be overlayed with the RMSD of 0.1 Å. The potential energy profile for the interconversion between Co–C4′ adduct, photoproduct and Co–C5′ adduct (Supplementary Fig. 23) was calculated using relaxed potential energy scans, and all transition states were fully optimized and frequency calculations confirmed the presence of a single imaginary frequency. Results of cluster model calculations are presented and discussed in detail in Supplementary Note 6.

### QM/MM setup and simulation

Computational models of dark and intermediate state structures were created based on a monomer of the corresponding crystal structures (*dark_ref_SACLA, 10ns_30mJ/cm*^*2*^*_SACLA*, and *3µs_30mJ/cm*^*2*^*_SACLA*) and minimized using the ff14sb AMBER force field^63^. Hydrogens were added with Amber16^64^. The AMBER parameters for coenzyme-B_12_ were obtained from Marques, *et al*.^65^. In preparation for quantum mechanics/molecular mechanics (QM/MM) simulations, the models were divided into a higher (QM) and a lower (MM) layer (Supplementary Fig. 39). The QM layer contained 170 atoms and comprised the chromophore’s adenosyl group, corrin ring (including cobalt ion and corrin side chains) and the imidazole portion of H177. The MM layer comprised the chromophore’s nucleotide loop (including the dimethylbenzimidazole), the H177 backbone atoms and all atoms of residues other than H177. The QM region was treated with density functional theory (DFT) using the BP86 functional along with the TZVP basis set for H and the TZVPP basis set for Co, C, N, and O^66-70^. Any residue which had at least one atom within 10 Å of the cobalt was kept unfrozen, meaning it was allowed to freely move throughout the course of the optimisations. The QM/MM models were optimized using Gaussian09^71^.

## Data availability

Crystal structures have been deposited at the Protein Data Bank (PDB), accession codes: 9S0A, 9S0B, 9S0C, 9S0D, 9S0E, 9S0F, 9S0G, 9S0H, 9S0I, 9S0J, 9S06, 9S07, 9S08, 9S09.

## Acknowledgments

M.W., G.S., and J.-P.C. warmly thank Ilme Schlichting, Robert Shoeman, Bruce Doak and Thomas Barends for long-standing support, advice and collaboration related to all aspects of serial crystallography experiments. G.S. and M.L are grateful to Marco Cammarata for the invaluable long-term advice and collaboration and the very helpful discussions on several aspects of the time-resolved X-ray solution scattering experiments. M.W. acknowledges a stimulating discussion with David Le Bolloc’h on energy dissipation in photoreceptors. The authors thank Elena A. Andreeva for advice on viscous matrix preparation for microcrystal injection. The study was supported by ANR grants to G.S. and M.L. (PhotoGene) and to M.W. (BioXFEL), by CNRS grants to G.S. (XFEL Instruments) and to M.W. (GoToXFEL) and a grant to M.W. by GRAL, a project of the University Grenoble Alpes graduate school (Ecoles Universitaires de Recherche) CBH-EUR-GS (ANR-17-EURE-0003). MW and TT acknowledge financial support from the Partenariat Hubert Curien (PHC) Sakura program, jointly funded by the Ministère de l’Europe et des Affaires étrangères (MEAE) and the Ministère de l’Enseignement supérieur et de la Recherche (MESR) of France, and the Japan Society for the Promotion of Science (JSPS) of Japan. R.M. is also supported by a GRAL PhD Fellowship to J.P.C. We thank the EPSRC for access to the National Research Facility for Electron Paramagnetic Resonance Spectroscopy (NS/A000055/1, EP/W014521/1). Funding from an EPSRC International Centre-to-Centre grant no. EP/S030336/1 was awarded to D.J.H., D.L. and N.S.S. R.R.-S. acknowledges a CEA CFR PhD fellowship and travel grants from EMBO (scientific exchange grant number 9591) and the Université Grenoble Alpes for a one-month stay at SACLA (Research Support Program for Graduate Students). XFEL experiments were carried out at BL2-EH3 of SACLA with the approval of the Japan Synchrotron Radiation Research Institute (JASRI; Proposal No. 2022B8815), at the SwissMX instrument at the Cristallina experiment station of SwissFEL (proposal 20231018). This project has received funding from the European Union’s Horizon 2020 research and innovation programme under the Marie Skłodowska-Curie grant agreement No 884104 (PSI-FELLOW-III-3i). The SwissFEL Cristallina experimental station at the Paul Scherrer Institut was realized with financial support from the Swiss National Science Foundation and the University of Zurich under Project Nr 206021_183330. We warmly thank the SACLA and SwissFEL staff for assistance. M.W., G.S., and R.R.-S. warmly thank Ilme Schlichting and Robert L. Shoeman, and Sofia M. Kapetanaki for their time invested during the screening beamtimes at the LCLS (P240) and the EuXFEL (PCS proposal N°3007), respectively. The ESRF is acknowledged for providing beamtimes (LS-3091, BLC-14546, LS-3226, LS-3303, LS-3364) at the ID09 beamline. We acknowledge the French Biology/Health Panel Review Committee for provision of synchrotron radiation beamtime at the ESRF on beamline BM07-FIP2, supported by the French ANR PIA3 (France 2030) EquipEx+ project MAGNIFIX under grant agreement ANR-21-ESRE-0011. SSX experiments were carried out at the I24 beamline at Diamond Light Source under proposal numbers MX24447 and MX31850. The PORTO laser at Diamond was funded by the EPSRC (EP/P001548/1). This work was partially carried out at the platforms of the Grenoble Instruct-ERIC center (IBS and ISBG; UMS 3518 CNRS-CEA-UGA-EMBL) within the Grenoble Partnership for Structural Biology (PSB). Platform access was supported by FRISBI (ANR-10-INBS-05-02) and GRAL. The IBS acknowledges integration into the Interdisciplinary Research Institute of Grenoble (IRIG, CEA). M.J.M. and P.M.K. wish to acknowledge the Cardinal Research Cluster at the University of Louisville for access to high-performance computing resources.

## Author contributions

D.J.H. and G.S. conceived the project. D.J.H., M.L., N.S.S., M.W., and G.S. designed and organized the overall research. The main manuscript was written by D.J.H., M.J.M., M.W., and G.S., with input from R.R.-S., H.P., N.C., L.O.J., L.N.J., E.D.Z., K.H., S.H., S.J., R.O., J.B., J.-P.C., T.T., M.L., S. Hay, P.M.K., D.L., and N.S.S., while the Supplementary Information was written by R.R.-S., H.P., K.P., D.J.H., N.C., M.J.M., L.O.J., S.S., L.N.J., M.L., S. Hay, P.M.K., M.W., and G.S. Protein expression and purification were carried out by R.R.-S., H.P., S.S., M.D., and N.Z., for WT-*Tt*CBD, H.P., and M. Sakuma for *Tt*CBD-H132A and full-length *Tt*CarH. Microcrystals for SFX were produced by R.R.-S., S.S., and K.H., and for SSX by H.P. Macrocrystals for MX were grown by R.R.-S., H.P., and S.S. Microcrystal injection conditions were established by R.R.-S., and G.S. Microcrystal diffraction quality was tested at ESRF and SPring-8 by R.R.-S., T.T., and G.S. SFX experiments at SACLA and SwissFEL were performed by R.R.-S., H.P., D.J.H., N.C., S.S., L.N.J., E.D.Z., M.A., E.B., M.D., G.G., S.J.O.H., K.I., M.K., R.K., T.M., R.M., H.O., S.O., A.S., M. Sakuma, H.S., K.T., J.B., J.-P.C., T.T., M.W., and G.S. SSX experiments at DLS were performed by H.P., D.A., S.H., S.J., G.K., and R.O. Off-line data processing, analysis, and structure refinement were conducted by R.R.-S., N.C., E.D.Z., and J.-P.C., for SFX and by H.P., N.C. for SSX. Cryo-temperature dependent MX experiments were carried out by R.R.-S., H.P., D.J.H., S.S., L.N.J., M.D., S.E., A.R., M.W., and G.S., with data analysis and structure refinement by R.R.-S., N.C., and S.S. TR-XSS experiments at ID09/ESRF were performed by R.R.-S., H.P., K.P., D.J.H., L.N.J., M.L., and G.S., with data reduction by K.P. and M.L. and analysis and interpretation by K.P., J.S.H., M.L., and G.S. NMR spectra were collected and interpreted by L.N.J. and M.J.C. LC-MS data were collected and interpreted by L.N.J. QM/MM calculations were performed, analyzed, and interpreted by M.J.M., and P.M.K. DFT cluster calculations were performed, analysed, and interpreted by L.O.J., and S. Hay. Time- and temperature-resolved UV-Vis spectroscopy was carried out and interpreted by H.P., D.J.H., and S.J.O.H., and EPR data were collected and interpreted by H.P., D.J.H. and M. Shanmugam.

