## Extended Data Figures for "Integrated structural dynamics uncover new modes of B_12_ photoreceptor activation"

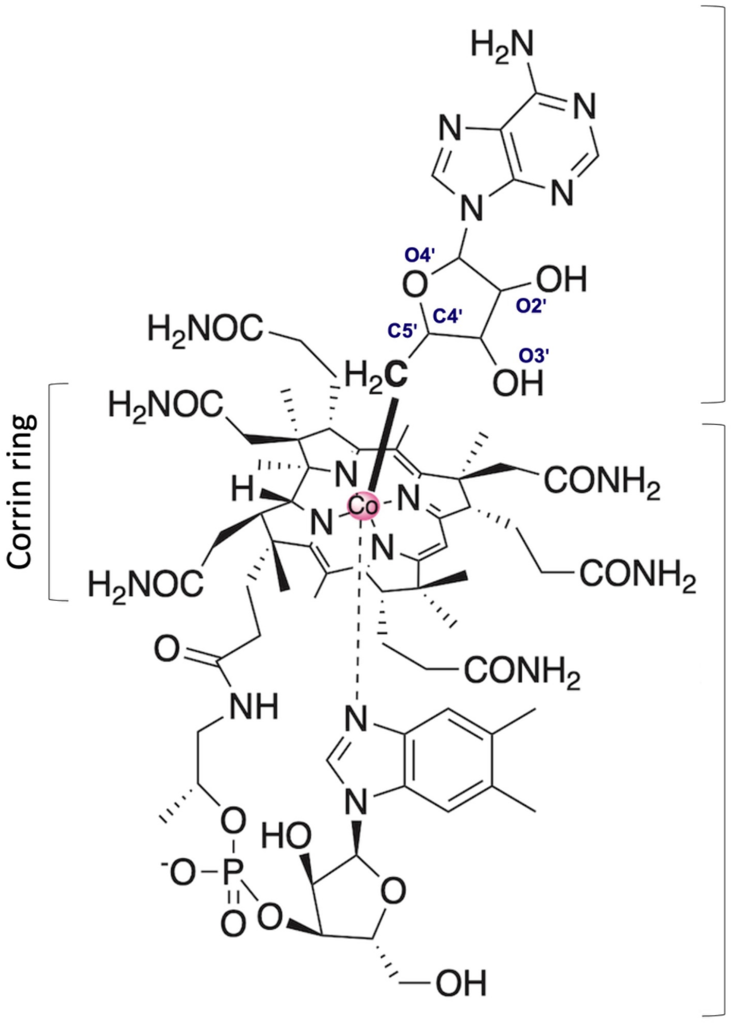


**Extended Data Fig. 1. Chemical structure of adenosylcobalamin (AdoCbl).**

The cobalt atom is shown as a pink sphere and the photolabile Co-C5′ bond depicted in bold.

.

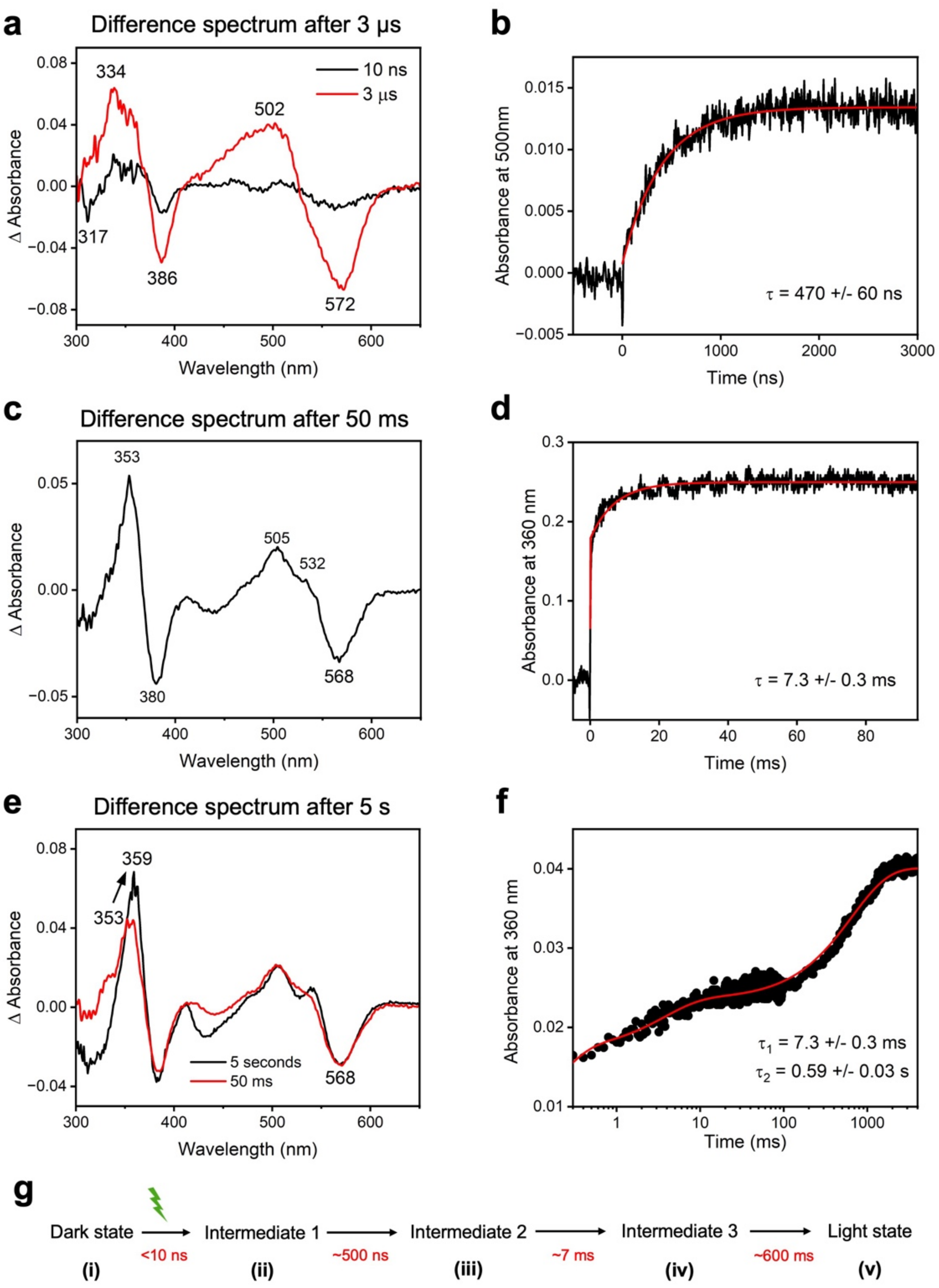


**Extended Data Fig. 2. Formation of intermediate states in the photochemical reaction pathway of *Tt*CBD in solution monitored by TR-absorption spectroscopy after excitation at 530 nm.**
Time-dependent difference absorbance spectra of samples containing 50 µM *Tt*CBD are shown 10 ns, 3 µs (**a**), 50 ms (**c**) and 5 s (**e**) after excitation with a ns laser pulse at 530 nm. The wavelengths of the main spectral changes are labelled accordingly. (**b**) Kinetic transients at 500 nm on the µs timescale of samples containing 50 µM *Tt*CBD. (**d**) Kinetic transients at 360 nm on the ms timescale of samples containing 50 µM *Tt*CBD. (**f**) Kinetic transients at 360 nm on the ms-s timescale of samples containing 50 µM *Tt*CBD. Data were fitted to a single exponential equation (red lines) to obtain time constants. All transients were measured at room temperature and data shown are the average of at least five traces. (**g**) The time-resolved absorption spectroscopy data yield the reaction scheme shown here, indicating the time constants for the formation of each of the intermediate states. The *i*, *ii*, *iii*, *iv* and *v* labels illustrate how these species relate to those depicted in Fig. 1 of the main text.

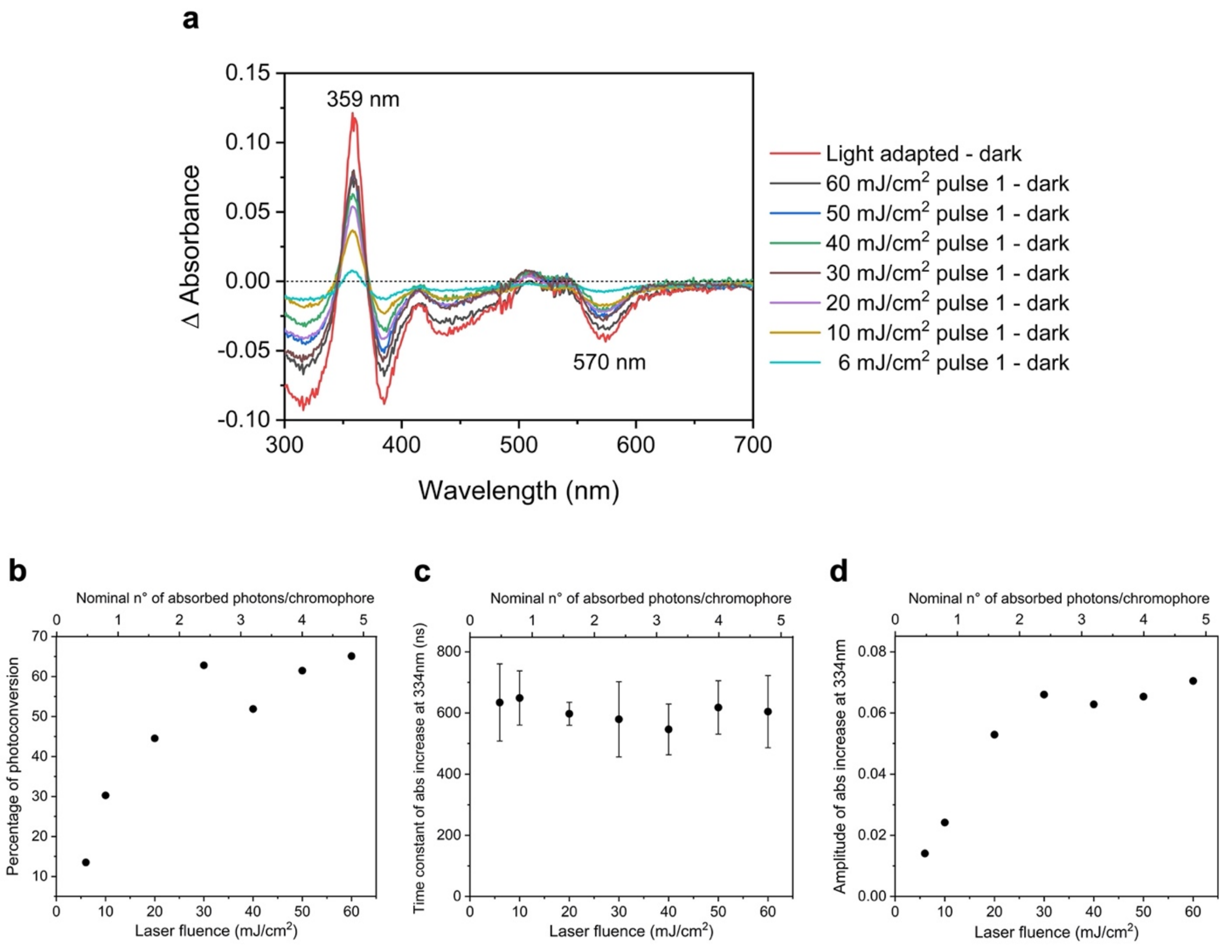


**Extended Data Fig. 3. Spectroscopic pump-laser power titration of photoconversion in *Tt*CBD microcrystals and in solution.**

(**a**) Absorbance difference spectra of *Tt*CBD microcrystals in 4% CMC before and after excitation with a single ns laser pulse of varying laser fluence at 530 nm from 6 to 60 mJ/cm². The non-illuminated sample was used as blank. (**b**) The amount of light state formed in *Tt*CBD microcrystals in 4% CMC as a function of laser fluence (converted to the corresponding number of absorbed photons per chromophore). The amount of light-adapted state was determined by measuring the absorbance at 359 nm, which is characteristic of the bis-histidine light-adapted state. The fully converted light-adapted state was measured by illuminating a *Tt*CBD microcrystalline sample for 2 min with a 530 nm LED. (**c**) The measured time constant for the formation of intermediate 2 (species *iii* in Fig. 1), as determined by monitoring the absorbance change at 334 nm upon excitation with a laser pulse at 530 nm, of *Tt*CBD in solution as a function of laser fluence. (**d**) The amplitude of the absorbance change at 334 nm upon excitation with a laser pulse at 530 nm of *Tt*CBD in solution as a function of laser fluence.

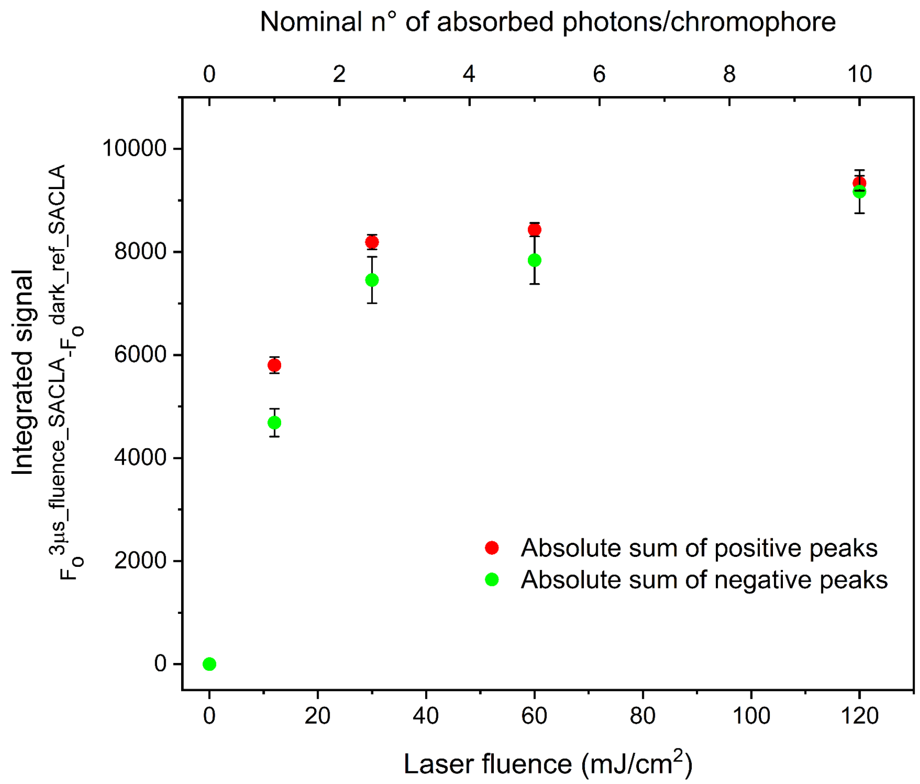


**Extended Data Fig. 4. Crystallographic pump-laser power titration using the light-induced difference signal at 3 µs time delay.**

Absolute sum of positive (> 3$\sigma$, green) and negative (< -3$\sigma$, red) integrated peaks in the Fourier difference maps calculated between light and dark data sets ($F_o^{3µs\_fluence\_SACLA}$ - $F_o^{dark\_ref\_SACLA}$, fluence: 12, 30, 60 and 120 mJ/cm$^2$) and located within 2 Å from the AdoCbl chromophore was computed and plotted as a function of pump-laser fluence.

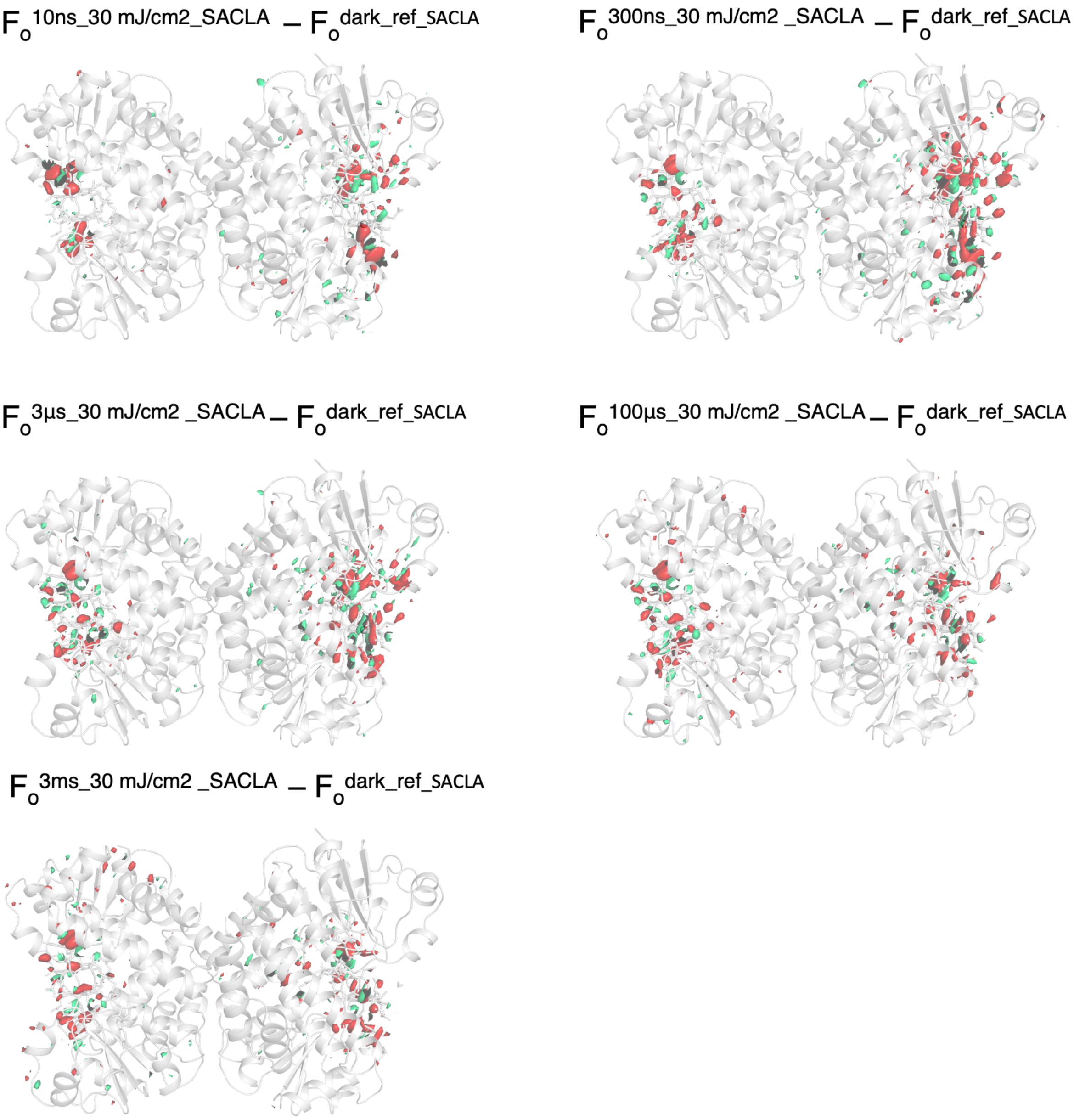


**<u>Extended Data Fig. 5. Overall view of Fourier difference maps from the time series collected at SACLA.</u>**

Model of *dark_ref_SACLA Tt*CBD tetramer is represented as a grey cartoon. Fourier difference map $F_o^{\Delta t\_30mJ/cm2\_SACLA} - F_o^{dark\_ref\_SACLA}$ (**Δ**t: 10 ns, 300 ns, 3 µs, 100 µs, 3 ms) are contoured at +3.5 (green) and -3.5 $\sigma$ (red).

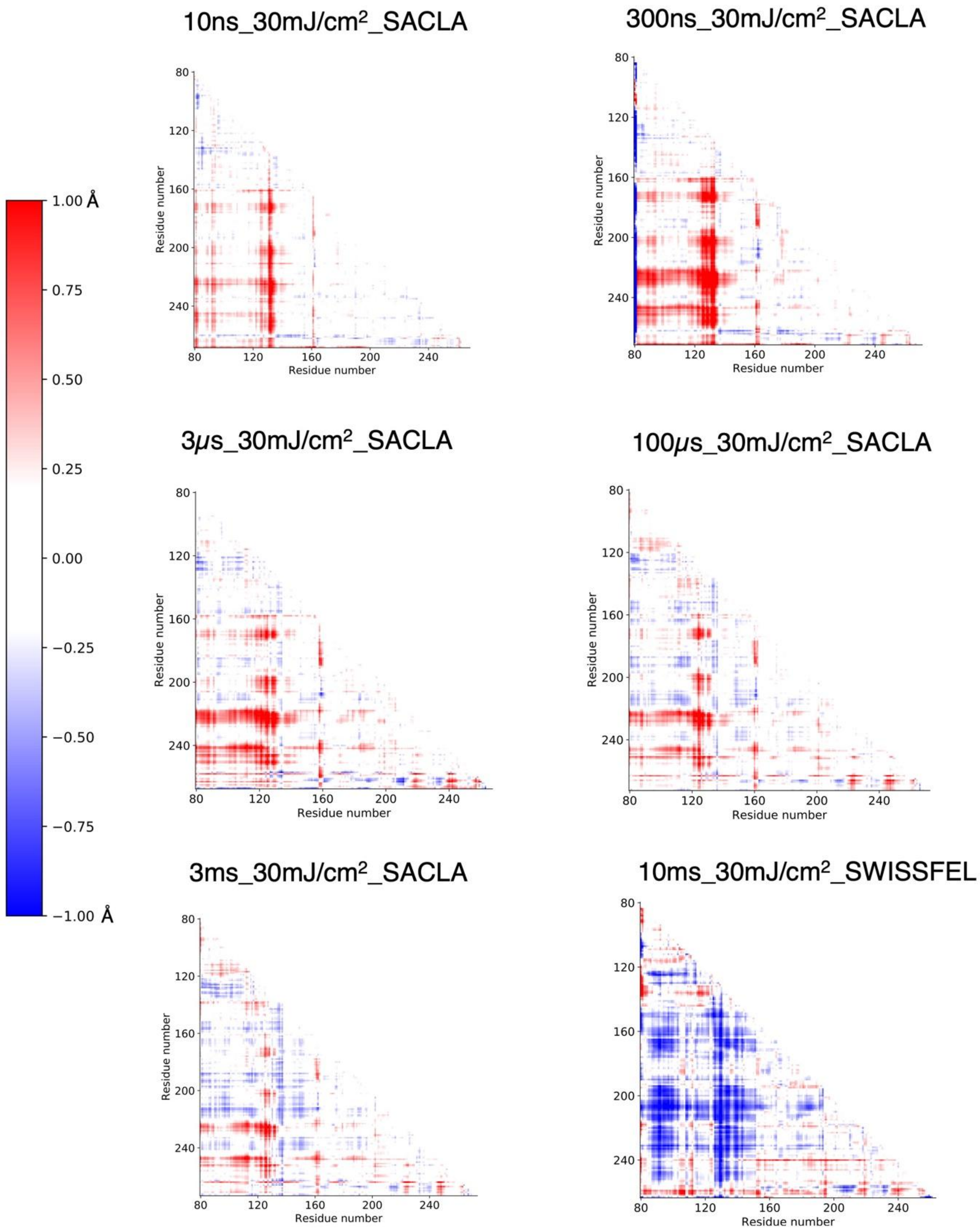


**Extended Data Fig. 6. Intrachain distance difference matrices (DDM) of monomer B at 10 ns, 300 ns, 3 µs, 100 µs, 3 ms, and 10 ms.**

DDMs were calculated using Dark_ref_SACLA as the reference model from 10 ns to 3 ms and Dark_ref_SwissFEL for 10 ms.

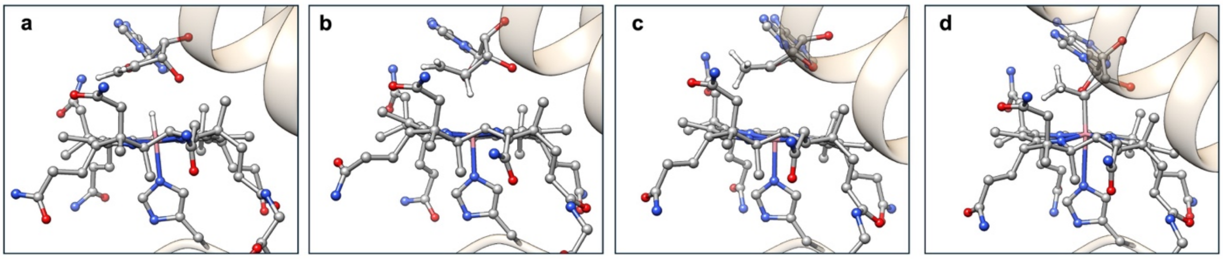


**Extended Data Fig. 7. QM/MM optimized models of *3μs_30mJ/cm$^2$_SACLA*.**

QM/MM models generated from the 3 μs crystal structure where four possible scenarios are optimized including (**a**) the photoproducts, (**b**) a primary diradical species where the Co(II)/C5′• radical pair (RP) are formed, (**c**) a tertiary diradical species where the Co(II)/C4′• RP are formed, and (**d**) a stable Co-C4′ adduct.

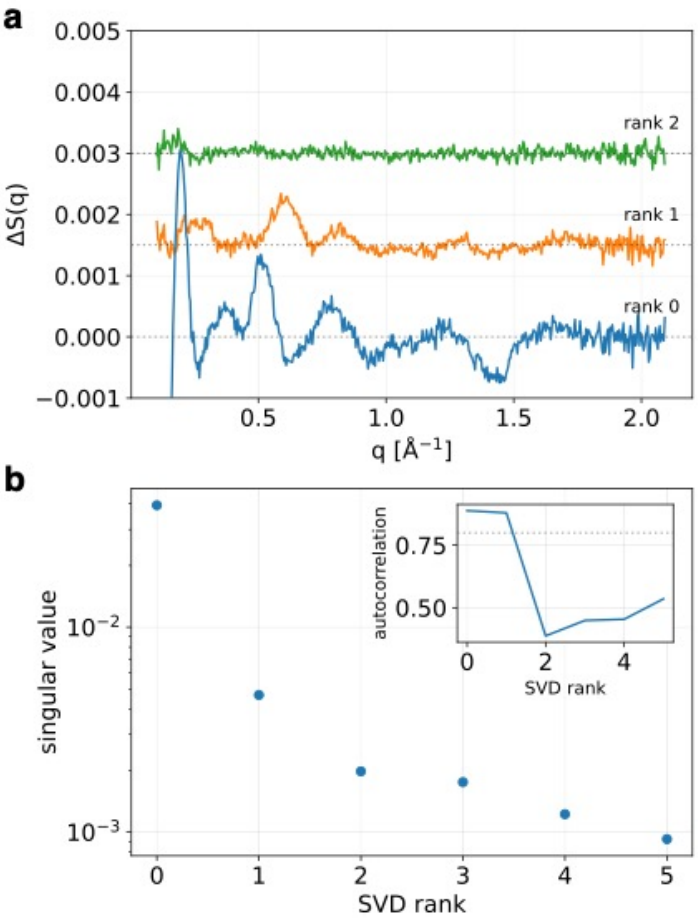


**Extended Data Fig. 8. SVD analysis of TR-XSS data in the 30 µs - 100 ms time range.**

(**a**) First three SVD basis patterns (left singular vectors) (rank 0 - blue, rank 1 - orange, rank 2 - green). (**b**) The singular values are plotted with blue dots as a function of the rank. The inset shows the autocorrelation of the basis patterns as a function of the rank. A grey dotted line indicates an autocorrelation of 0.8. SVD analysis shows that only two basis patterns contain a signal out of noise.

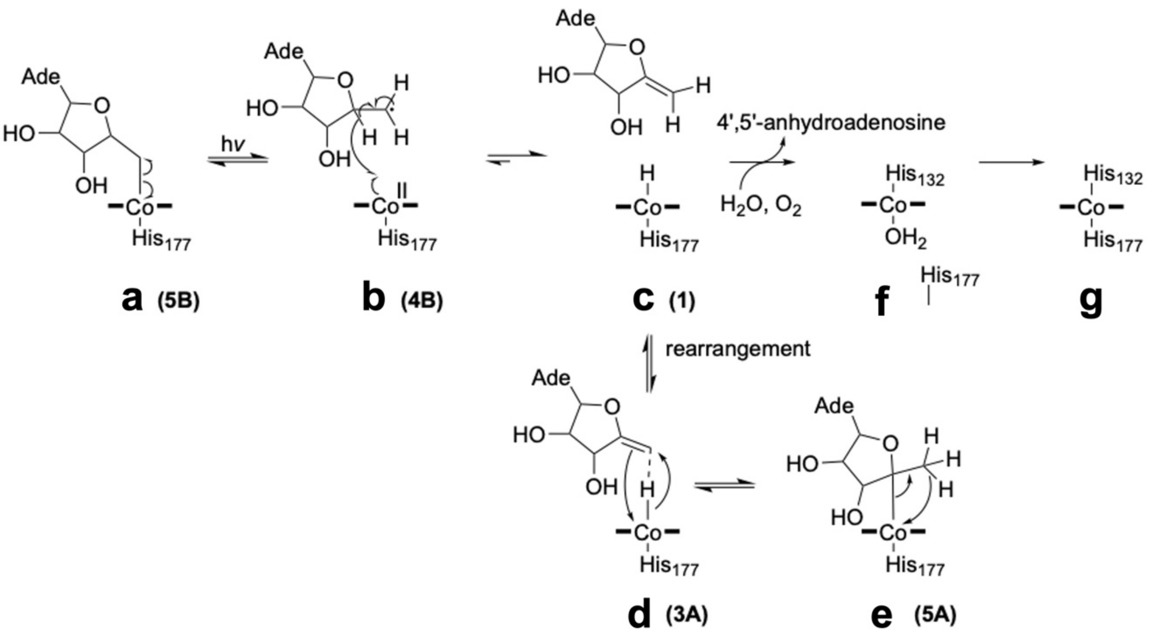


**Extended Data Fig. 9. Proposed CarH photoactivation mechanism based on DFT cluster model calculations.**

Shown is homolytic photolysis from the dark state (**a**), to a diradical (**b**) followed by formation of the photoproduct (**c**), which can lead to the reversible formation of the Co–C4′ adduct (**e**), via an undetected intermediate species (**d**), or the release of the photoproduct (4′,5′-anhydroadenosine), which leads to the formation of a water-ligated Co(III) species (**f**) and the final light state (**g**)**.** Names in parentheses refer to the corresponding structures (where appropriate) from Supplementary Fig. 23 and Supplementary Tables 3 and 10. The adenine (Ade) moiety of the 5′-deoxyadenosyl group was omitted for clarity.
