## Supplementary_information for "Integrated structural dynamics uncover new modes of B_12_ photoreceptor activation"

**This document file includes:**

Supplementary Methods

Supplementary Notes 1 to 6

Supplementary Figures

Supplementary Tables

**Other supplementary information for this manuscript includes:**

Cluster model coordinates

**Table of contents**

**Supplementary Methods** 9

Expression and purification of recombinant *Tt*CBD 9

Time-resolved absorption spectroscopy 10

Temperature-resolved absorption spectroscopy 11

*Tt*CBD macrocrystallization for the production of seeds 11

*Tt*CBD microcrystallization for the time-resolved serial femtosecond crystallography (TR-SFX) experiment at SACLA 12

*Tt*CBD microcrystallization for the TR-SFX experiment at SwissFEL 12

Microcrystallization of *Tt*CBD H132A mutant for the SSX experiment at Diamond Light Source 13

Determination of photoproduct formed when illuminating *Tt*CBD crystals 13

Embedding *Tt*CBD microcrystals in a viscous carrier medium for high-viscosity extrusion injection during the TR-SFX experiment at SACLA 14

Offline HVE jet-speed calibration 15

TR-SFX data collection at SACLA 15

Processing of TR-SFX data collected at SACLA 17

TR-SFX data collection at SwissFEL 17

Processing of TR-SFX data collected at SwissFEL 18

SSX data collection on *Tt*CBD-H132A mutant microcrystals at Diamond Light Source 19

Processing of *Tt*CBD-H132A mutant SSX data collected at Diamond Light Source 20

Calculation of Fourier difference maps 20

Dependency of integrated Fourier difference peaks as a function of indexed diffraction images at SACLA 20

Assessing potential light contamination of interleaved dark images collected at SACLA 21

Assessing potential light contamination of interleaved dark images collected at SwissFEL 21
*Dark_ref_SACLA* structure refinement 22
*Dark_ref_SwissFEL* structure refinement 23
*Dark_DLS* refinement of H132A mutant structure 23
Structure factor amplitudes extrapolation of TR-SFX data sets 23
Refinement of TR-SFX intermediate-state structures 24
Refinement of SSX H132A mutant structures 24
Coordinate error estimation based on the bootstrapping method 26
Calculation of difference distance matrices (DDM) 26
Cryo-temperature dependent macromolecular crystallography and *in crystallo* UV-vis spectroscopy 27
Low temperature cw-EPR on *Tt*CBD after steady-state illumination at 180 K 28
Preparation of sample for time-resolved X-ray solution scattering (TR-XSS) 29
TR-XSS: data collection and reduction 30
Singular value decomposition of TR-XSS data 31
XSS difference signals calculated from PDB files 32
Cluster-model calculations 32
QM/MM setup and simulations 33
**Supplementary Notes 34**
Supplementary Note 1. Time-resolved absorption spectroscopy of *Tt*CBD in solution and in microcrystals 34
Supplementary Note 2. Spectroscopic and crystallographic pump power titration 35
Supplementary Note 3. Comparison of intermediate-state models in all monomers at the different TR-SFX time-delays 38
Supplementary Note 4. SSX on *Tt*CBD H132A mutant microcrystals at Diamond Light Source 43

Supplementary Note 5. Cryotrapping of intermediate states of *Tt*CBD reaction pathway in solution and *in crystallo* 44

Supplementary Note 6. Cluster-model and QM/MM calculations 47

**Supplementary figures** **52**

Supplementary Fig. 1. Crystal structures of full-length CarH before and after illumination. 52

Supplementary Fig. 2. Comparison of time-resolved absorption spectroscopy changes between *Tt*CBD in solution and *Tt*CBD microcrystals embedded in cellulose. 54

Supplementary Fig. 3. LCMS of the photoproduct formed when illuminating *Tt*CBD crystals 55

Supplementary Fig. 4. Determination by NMR of photoproduct formed when illuminating *Tt*CBD microcrystals 56

Supplementary Fig. 5. *Tt*CBD microcrystals 57

Supplementary Fig. 6. Fourier difference maps computed with the various data sets from the power titration performed at SACLA (3 µs time delay). 58

Supplementary Fig. 7. Fourier difference maps of interleaved dark data sets at different time delays and/or laser fluences compared to *dark_only_SACLA*. 60

Supplementary Fig. 8. Signal in the Fourier difference maps from the time-series collected at SACLA observed in the chromophore binding pocket. 62

Supplementary Fig. 9. Fourier difference maps calculated between interleaved darks and dark_only data sets collected at SwissFEL 63

Supplementary Fig. 10. Fourier difference maps of the 3 µs and 10 ms time delays at a pump-laser fluence of 30 mJ/cm$^2$ (SwissFEL). 64

Supplementary Fig. 11. Fourier difference maps of 3 µs and 10 ms time delays at a laser fluence of 30 mJ/cm$^2$ (SwissFEL). 65

Supplementary Fig. 12. *10ns_30mJ/cm$^2$_SACLA* intermediate-state model and 2m$F_{extr}$-D$F_c$ electron density and m$F_{extr}$-D$F_c$ polder maps (adenosyl moiety omitted). 66

Supplementary Fig. 13. *300ns_30mJ/cm²_SACLA* intermediate-state model and $2mF_{extr}$-$DF_c$ electron density and $mF_{extr}$-$DF_c$ polder maps (omitted adenosyl moiety). 67

Supplementary Fig. 14. *3μs_30mJ/cm²_SACLA* intermediate-state model and $2mF_{extr}$-$DF_c$ electron density and $mF_{extr}$-$DF_c$ polder maps (omitted adenosyl moiety). 68

Supplementary Fig. 15. Comparison of Fourier difference maps, intermediate-state models and $2mF_{extr}$-$DF_c$ electron density maps of *3μs_30mJ/cm²_SACLA* and *3μs_30mJ/cm²_SwissFEL* 69

Supplementary Fig. 16. *100μs_30mJ/cm²_SACLA* intermediate-state model and $2mF_{extr}$-$DF_c$ electron density and $mF_{extr}$-$DF_c$ polder maps (omitted adenosyl moiety). 71

Supplementary Fig. 17. *3ms_30mJ/cm²_SACLA* intermediate-state model and $2mF_{extr}$-$DF_c$ electron density and $mF_{extr}$-$DF_c$ polder maps (omitted adenosyl moiety). 72

Supplementary Fig. 18. *10ms_30 mJ/cm²_SwissFEL* intermediate-state model and $2mF_{extr}$-$DF_c$ electron density and $mF_{extr}$-$DF_c$ polder maps (adenosyl moiety omitted). 73

Supplementary Fig. 19. Cryotrapping of intermediate states in the photochemical reaction of *Tt*CBD in solution as monitored by temperature-resolved absorption spectroscopy measurements. 74

Supplementary Fig. 20. *In crystallo* UV-vis spectroscopy on *Tt*CBD CarH at BM07-FIP2 beamline (ESRF). 75

Supplementary Fig. 21. *Cryo_light_180K* model and $2mF_o$-$DF_c$ electron density and $mF_o$-$DF_c$ polder maps (adenosyl moiety omitted). 76

Supplementary Fig. 22. Characterization of intermediate 2 state in the photochemical reaction of aerobic *Tt*CBD in solution as monitored by temperature-resolved EPR spectroscopy measurements 77

Supplementary Fig. 23. Proposed mechanism for the interconversion of the Co–C4′ and Co–C5′ adducts via the photoproduct based on cluster model calculations. 78

Supplementary Fig. 24. Absorption solution spectra of the *Tt*CBD H132A mutant light state at room temperature and of the *Tt*CBD intermediate 2 cryotrapped at 260 K. 79

Supplementary Fig. 25. Comparison of the XSS signals calculated using the cryotrapped 8C76 structure and the SFX 10 ms intermediate. 80

Supplementary Fig. 26. Comparison of the 100 ms TR-XSS patterns from different data sets. 81

Supplementary Fig. 27. Effect of tetramer-dimer and tetramer-monomer transitions on the XSS signal. 82

Supplementary Fig. 28. Longest TR-XSS signal and SVD analysis of TR-XSS data in the 100 ms - 3 s time range. 83

Supplementary Fig. 29. Removal of solvent heating signal from TR-XSS data. 84

Supplementary Fig. 30. Correction of TR-XSS data for the irreversible photoconversion kinetics. 85

Supplementary Fig. 31. Purification of *Tt*CBD assessed by SDS-PAGE 86

Supplementary Fig. 32. Size exclusion chromatogram of *Tt*CBD. 87

Supplementary Fig. 33. UV-vis absorption spectroscopy of *Tt*CBD in solution 88

Supplementary Fig. 34. Jet speed calibration as a function of the HPLC flowrate setting. 89

Supplementary Fig. 35. Dependency of integrated Fourier difference peaks as a function of indexed lattices. 90

Supplementary Fig. 36. *illuminated_DLS* intermediate-state model of *Tt*CBD-H132A and $2mF_o$-$DF_c$ electron density and $mF_o$-$DF_c$ polder maps (adenosyl moiety omitted) 91

Supplementary Fig. 37. Online UV-vis microspectrophotometer mounted on the diffractometer of the BM07-FIP2 beamline of the ESRF 92

Supplementary Fig. 38. Cluster model. 93

Supplementary Fig. 39. QM/MM Partitioning. 94

Supplementary Fig. 40. Penetration of 530 nm light into *Tt*CBD microcrystals. 95

Supplementary Fig. 41. Absorbed photons *vs.* penetration depth in a *Tt*CBD crystal 96

Supplementary Fig. 42. Comparison of the Fourier difference map and the intermediate-state models at 10 ns and 3 μs time delays and at pump-laser fluences of 12 and 30 mJ/cm$^2$ (SACLA). 97

Supplementary Fig. 43. Root mean square fluctuation (RMSF) obtained from the ensemble refinement of the *dark_ref_SACLA* data set. 99

Supplementary Fig. 44. *Dark_ref_SACLA* model and 2m$F_o$-D$F_c$ electron density and m$F_o$-D$F_c$ polder maps (omitted adenosyl moiety). 100

Supplementary Fig. 45. Time-resolved absorption spectroscopy on the ms time scale in *Tt*CBD H132A mutant in solution after excitation at 530 nm. 101

Supplementary Fig. 46. Comparison of QM/MM optimized models with TR-SFX structures 102

Supplementary Fig. 47. CarH Diradical Potential Energy Surface. 103

**Supplementary Tables 104**

Supplementary Table 1. AdoCbl geometric parameters at 10 ns. 104

Supplementary Table 2. AdoCbl geometric parameters at 3 μs. 105

Supplementary Table 3. Ground state potential energies relative to the photoproduct (HCbl + anhAdo) and selected structural parameters for the cluster model. 106

Supplementary Table 4. Data collection and refinement statistics of TR-SFX power titration at SACLA 107

Supplementary Table 5. Data collection and refinement statistics of TR-SFX time series at SACLA 109

Supplementary Table 6. SwissFEL TR-SFX Data collection and refinement statistics 111

Supplementary Table 7. Diamond SSX Data collection and refinement statistics of *Tt*CBD mutant - H132A 113

Supplementary Table 8. Cryo-T dependent MX Data collection and refinement statistics 115

Supplementary Table 9. Comparison of Co–C5′ and Co–C4′ distances in TR-SFX structures at 10 ns and 3 μs at two pump-laser fluences. 116

Supplementary Table 10. Atomic charges and spin densities for selected atoms in the cluster model. 117

Supplementary Table 11. AdoCbl geometric parameters in the dark state. 118

**Supplementary References** **119**

### Supplementary Methods

#### Expression and purification of recombinant *Tt*CBD

The expression and purification protocol was adapted from the previously published one for full-length CarH[1]. Apo-*Tt*CBD (*Thermus thermophilus* CarH cobalamin binding domain, residues 78-285) with a C-terminal 6×Histag was cloned in *E. coli* BL21 (DE3) cells (New England Biolabs) by the introduction of the expression vector pET21b (ampicillin resistance) containing the gene. A single colony from the freshly transformed plate was used to inoculate 100 ml of LB medium supplied with 0.1 ml of ampicillin 100X (Euromedex, 100 µg/ml). The pre-culture was kept in agitation at 150-170 rpm at 37 °C and left overnight. The pre-culture was then used to inoculate 1 l of fresh LB medium, adjusting the initial optical density at 600 nm (OD600) to around 0.05 and adding 1 ml of ampicillin at 100 mg/ml. The culture was kept in agitation at 160 rpm at 37 °C until the OD reached a value of ~ 0.5 (usually reached after 2-2.5 hours). Adding IPTG (Euromedex) to a final concentration of 0.5 mM (500 µl IPTG 1 M in 1 l culture) induced the protein expression. The temperature was reduced to 25 °C and the cultures were allowed to grow for another 20 h with agitation at 160 rpm. Cells were harvested by centrifugating the overnight culture at 5000 g for 30 min at 4 °C. From 2 l of culture, ~ 8 g of wet pellet was obtained and stored at -20 °C. Cells were thawed at room temperature and suspended in 100 ml of buffer A (50 mM Tris-HCl pH 7.5, 100 mM NaCl). One complete tablet of protease inhibitor (Roche, 5056489001), DNase (Sigma, DN25-1G) at a concentration of 25 µg/ml, $MgCl_2$ and $CaCl_2$ at 5 mM were added to the cell suspension. The cells were always kept on an ice bath and were lysed using a sonicator (40% power output, 10 s on, 50 sec off, 30 cycles, 4 min of sonication in total). The lysed cells were centrifuged at 39000 g for 30 min and the cell debris was removed. The supernatant containing the apo-*Tt*CBD was collected and imidazole was added to a final concentration of 10 mM.

Protein purification was achieved using a nickel column, followed by size exclusion chromatography (SEC). In between the two purification steps, the chromophore adenosylcobalamin (AdoCbl) was added to the apo-*Tt*CBD. Purification of the apo-*Tt*CBD solution was performed using a Ni-NTA gravity flow column on the bench at room temperature and under ambient light conditions. The resin was washed with 100 ml of water followed by 100 ml of buffer A. The supernatant was loaded onto the column which was then washed with 100 ml of wash buffer (buffer A + 10 mM imidazole) and the flow-through was discarded. The protein

was then eluted using 100 ml of buffer B (50 mM Tris-HCl pH 7.5, 100 mM NaCl, 250 mM imidazole). The presence of *apo-Tt*CBD was confirmed by SDS-PAGE (Supplementary Fig. 31).
To generate the holo-*Tt*CBD (called *Tt*CBD hereafter), around 30-35 mg of chromophore adenosylcobalamin (Sigma-Aldrich catalog n° C0884) was added to the 100 ml of the eluted solution (i.e. final chromophore concentration of 190 - 220 μM). AdoCbl:apo-*TtCBD* ratio should be greater than one to avoid protein precipitation in the subsequent protein-concentrating step. Since AdoCbl is light sensitive, all manipulations and subsequent purification steps were performed under red light conditions (> 650 nm) and the samples were stored in opaque containers. The solution was left overnight at room temperature and with constant agitation and then centrifuged and filtered through a 0.2 μm filter to remove any precipitate. The filtered solution was concentrated using 50 kDa MW cutoff concentrators down to 15 ml (*Tt*CBD concentration around 35 mg/ml).
For the SEC purification step, 2 ml of the previously concentrated *Tt*CBD sample was injected onto a Superdex 200 Hiload 16/600 prep grade column. The protein eluted in 50 mM Tris-HCl pH 7.5, 100 mM NaCl as a single peak around 70 ml (Supplementary Fig. 32) corresponding to a tetramer in its dark-state as assessed by UV-vis absorption spectroscopy (Supplementary Fig. 33). A minor peak at around 87 ml originates from residual monomeric *Tt*CBD (Supplementary Fig. 32). A remaining peak around 114 ml corresponds to the free AdoCbl used in excess. Only the top half fraction of the elution peak at 70 ml containing *Tt*CBD was collected for crystallization.

**Time-resolved absorption spectroscopy**

Time-resolved absorption spectroscopy experiments after laser photoexcitation were carried out at 298 K using an LP980-KS flash photolysis instrument (Edinburgh Instruments Ltd). Samples in solution contained 50 μM TtCBD (200 μM, or 500 μM when comparing with microcrystals) in 50 mM Tris-HCl pH 7.5 and 150 mM NaCl. Photoactivation was initiated by ns laser excitation at 530 nm, using an optical parametric oscillator of a Q-switched Nd-YAG laser (NT432, EKSPLA) in a 1-cm pathlength cuvette. Laser pulses were between 6-8 ns in duration and varied in energy up to 20 mJ by using an attenuator. Difference absorbance spectra were recorded between 300 and 700 nm at selected time points using an image intensified CCD camera (Andor Technologies). Single-wavelength kinetic absorption transients were recorded at a range of wavelengths between 300 and 700 nm with the detection system (comprising probe light, sample, monochromator and photomultiplier) at right angles to the incident laser beam. Rate constants were obtained from the average of at least five time-dependent absorption measurements by fitting to a single exponential function using Origin Pro 9.1 software. Single-wavelength kinetic absorption transients were also collected on samples of 20% (v/v) TtCBD microcrystals embedded in 4% (w/v) in sodium carboxymethyl cellulose (CMC, Sigma, see preparation details in (*Embedding TtCBD microcrystals in a viscous carrier medium for high-viscosity extrusion injection during the TR-SFX experiment at SACLA*) in a 1-mm pathlength cuvette at a 45° angle to the incident laser beam and the probe beam. Results of the time-resolved absorption spectroscopy measurements are presented and discussed in Supplementary Note 1.

#### Temperature-resolved absorption spectroscopy

All static absorbance spectra were measured using a Cary 50 spectrophotometer (Agilent Technologies). For temperature-resolved absorption spectroscopy measurements, 1 ml samples containing 50 μM *Tt*CBD in 50 mM Tris-HCl pH 7.5, 44 % glycerol (v/v), 20 % sucrose (w/v) were cooled to 77 K at an approximate rate of 10 K per minute in an Optistat DN liquid nitrogen cryostat (Oxford Instruments Inc.) to record spectra. Samples were then warmed to the desired temperature at an approximate rate of 10 K per minute to initiate the reaction, either by illumination (approx. 1000 $\mu mol/m^2/s$) with a 530 nm high power LED (Thorlabs Inc.) for 15 min or by subsequent incubation in the dark for 15 min (following the illumination at 180 K for 15 mins), before cooling again to 77 K to record spectra. Room temperature absorbance spectra were also collected on 20% (v/v) *Tt*CBD microcrystals embedded in 4% (w/v) CMC matrix in a 1-mm pathlength cuvette.

#### ***Tt*CBD macrocrystallization for the production of seeds**

Macrocrystals of *Tt*CBD were produced by the batch method, at room temperature (20 °C) and under red light conditions as described recently[2]. 500 μl of freshly produced protein at 8 mg/ml was mixed with 500 μl of crystallization buffer containing 20% PEG 10000, 0.1 M HEPES pH 7.5. After two days the crystals reached their final size of 10-20 × 300 × 100 $\mu m^3$. Macrocrystals produced under these conditions have been previously defined as form II in[2].

#### *Tt*CBD microcrystallization for the time-resolved serial femtosecond crystallography (TR-SFX) experiment at SACLA

For microcrystallization, form II macrocrystals were used to generate seeds under red-light conditions according to a recently reported protocol[3] as follows: twelve of the 1-ml batches of macrocrystals described in the previous section were centrifuged, pooled, and suspended in 1 ml of the storage buffer (10% PEG 10000, 0.05 M HEPES pH 7.5). This crystal slurry containing macrocrystals was divided into three equal parts, each filled into three 2-ml containers and mixed with the 1.5-mm zirconium beads (Sigma). The containers with the crystals and the beads were shaken for 45 s at 4000 rpm in a BeadBug homogenizer (Benchmark Scientific) followed by another 45 s in an ice bath to cool the sample. This cycle was repeated 12 times. Subsequently, the crystal suspension was transferred to a container with 1.0-mm beads. Washing the 1.5-mm bead container with 100 µl of storage buffer was carried to minimize the loss of crystal fragments. The 12 cycles were then repeated. Bead-exchange and crushing cycles were repeated with the 0.5 mm and 0.1 mm beads. The resulting crystal slurries were pooled and suspended in 10 ml of storage buffer to ease the following filtration step. The crystal slurry was filtered using 20, 10, 2.5, and 0.5 µm stainless steel frit filters. The slurry was then centrifuged for 30 min at 4000 g and the supernatant was removed until 2 ml remained, in which the seeds were suspended.

Microcrystals were grown by the batch method using the same crystallization buffer as for obtaining form II macrocrystals (20% PEG 10000, 0.1 M HEPES pH 7.5.) and adding the submicron seeds to a final concentration of 2.5% (v/v) as follows: in a 1.5 ml Eppendorf tube, 475 µl of crystallization buffer was mixed with 25 µl of suspended seeds and homogenized by pipetting vigorously. Then, 500 µl of freshly produced *Tt*CBD (i.e. without prior storage at cryo-temperatures) at 8 mg/ml were added, again by vigorous pipetting. After two days of incubation at room temperature, plate-like microcrystals reached their final size of ~ 15 × 5 × 2 µm (Supplementary Fig. 5).

#### *Tt*CBD microcrystallization for the TR-SFX experiment at SwissFEL

The same procedure as described in the previous section was used, with the following modifications. Twenty and not 12 of 1-ml batches of macrocrystals were used and the macrocrystals were fragmented using a mixture of zirconium beads (Sigma) with different diameters (1.5, 1.0, 0.5, and 0.1 mm). The fragmentation cycle was repeated 40 times. Seeds were resuspended in 15 ml of storage buffer. Microcrystals were grown exactly as described in the preceding section but submicron seeds were added at a final concentration of 1% instead of 2.5% (v/v). The shape and size of the microcrystals produced for the SwissFEL and SACLA experiments were identical.

#### Microcrystallization of *Tt*CBD H132A mutant for the SSX experiment at Diamond Light Source

Eluted fractions from size-exclusion chromatography corresponding to the top 25% of the dark-adapted tetrameric peak were used in microcrystallization for improved homogeneity. Previously we reported H132A macrocrystals growth[2] from a condition in the LMB screen (Molecular Dimension). The same condition was used for microcrystallization of the H132A mutant. Under dim red light conditions, 350 µl of protein, at a concentration of 50 mg/mL, was mixed with 650 µl of crystallization condition (0.1 M ammonium sulphate, 0.1 M sodium citrate, pH 5.8, 16% w/v PEG 4000, 20% v/v glycerol), and incubated at 20 °C in opaque black 1.5 ml volume Eppendorf tubes (Heathrow Scientific). Small rod-shaped crystals of ~15 × 8 × 5 $\mu m^3$ dimension appeared overnight.

#### Determination of photoproduct formed when illuminating *Tt*CBD crystals

Using the method utilized in[4] we sought to establish whether the same photoproduct is formed *in crystallo* as in solution. For this we used microcrystals produced for the TR-SFX experiments. For photoproduct elucidation we aimed for roughly 450 mg of protein per condition. Firstly, microcrystals were either kept in the dark or placed on a Camlab Choice MX-T6-S Classic Roller Mixer in a clear falcon tube in ambient light at room temperature for 30 min. Protein in solution, microcrystals and PEG-10000 molecules were separated from the mixture of buffer (50 mM Tris-HCl, 100 mM NaCl), crystallography condition (100 mM HEPES) and photoproduct using 10 kDa MWCO vivaspins (Sartorius). The flow through (~20-25 ml) was dried on a Buchi Rotavapor R-300 at 30 mbar 45 °C 250 rpm, resuspended in deuterated methanol and then dried again on a Genevac EZ-2 Mk3 using the low BP programme set at 45 °C. The final powder was resuspended in 550 µL deuterated methanol (Eurisotop). 50µL of the mixture was used for liquid chromatography mass spectrometry (LCMS) and the rest was used for NMR elucidation.

LCMS was undertaken on an Agilent 1100 LC-MSD instrument with an Agilent 150 × 3.0 mm Poroshell 120 SB-C18 (2.7 µM pore size) reversed phase column using $H_2O$ with 0.1% v/v formic acid as solvent A and acetonitrile as solvent B. A linear gradient from 0% to 95% B was used at 8 minutes for 28 min followed by 100% B for 20 minutes then re-equilibration with 0% B for 15 min. Every four samples, acetonitrile blanks were used to determine carry-over. An in-line ESI-TOF single quadrupole mass spectrometer (Agilent Technologies) in positive ion mode was used to determine mass-to-charge ratios (m/z) (Supplementary Fig. 3). For the MS the following settings were used: ESI capillary voltage 3000 V; gas temperature 350 °C; drying gas flow 11 L/min; nebulizer pressure 25 psi; fragmentor voltage 70 V; m/z scan range 120-1500.

NMR spectra were recorded on a Bruker AVIII 500 MHz spectrophotometer with 1H/19F/13C-15N QCI-F cryoprobe equipped with z-gradients. Initially, all NMR spectra were recorded using the same parameters as[4]. Briefly, 1D 1H NMR spectra were collected at 298 K using a 1D 1H NMR method with presaturation water suppression (noesygppr1d, 256 scan, 2s acquisition time, 4s recycle/saturation delay 50 Hz B1 saturation field strength). These spectra (Supplementary Fig. 4 red and green spectra) match those reported previously[4,5]. However, peaks responding to the 2′ and 3′ hydrogens were masked due to solvent, likely water. To resolve this, we recorded a spectrum at 273 K, with double presaturation at 3.7 ppm and 5.04 ppm providing observation of all peaks (Supplementary Fig. 4 blue spectrum). Further information regarding the full identification of the photoproduct is shown in[4].

**Embedding *Tt*CBD microcrystals in a viscous carrier medium for high-viscosity extrusion injection during the TR-SFX experiment at SACLA**

Sodium carboxymethyl cellulose was used as a medium for high-viscosity extrusion (HVE) injection of *Tt*CBD microcrystals during TR-SFX experiments. CMC stock gels were prepared as described previously[6] and adapted as follows: 10% (w/v) CMC was prepared by sprinkling 10 g into 90 ml of filtered milliQ water contained in a 500 ml bottle under constant stirring to yield a rigid gel and incubated overnight at 70 °C. A clear gel formed and was transferred with a spatula to 50 ml falcon tubes and centrifuged for 20 min at 4000 g at room temperature to remove air bubbles. To mitigate possible damage to the crystalline sample the carrier should contain the microcrystallization storage buffer. 25 g of the 10% CMC (w/v) stock was mixed in with 25 g (corresponding to a volume of ~ 25 ml) crystallization solution (20% PEG 10000, 0.1 M HEPES pH 7.5) and incubated at 70 °C overnight. To ease the mixing and the formation of a homogenous gel, the CMC stock gel was extruded through a plastic syringe to generate a continuous thread that enhances the surface contact with the crystallization solution. The resulting hydrogel, called CMC-5% hereafter, thus contained 5% CMC (w/v), 10% PEG 10000, and 0.05 M HEPES pH 7.5. For HVE injection, 10 µl of gravity-settled *Tt*CBD microcrystals were mixed with 190 µl of CMC-5% using two coupled 250 µl Hamilton syringes as described earlier[6].

**Offline HVE jet-speed calibration**

Prior to the TR-SFX experiment at SACLA (see below) the speed of HVE jets was determined for various nominal flow rates set at the pumping HPLC unit. This was done in a dedicated laboratory at the Institut de Biologie Structurale, Grenoble, France (Pepe *et al*, submitted) using methodology developed at the Max Planck Institute for Medical Research, Heidelberg, Germany[7]. Extrusion was carried out by a HVE injector[8] with a sample reservoir of 200 µl and a polyamide-coated glass capillary with an inner diameter of 75 µm. Jet speeds of CMC-5% carrying *Tt*CBD microcrystals were determined at several flow rates at the pumping HPLC unit ranging from 30 to 180 µl/min (Supplementary Fig. 34). The experimental protocol to image the jet and determine its speed is detailed in Pepe *et al*. (submitted). The same injector type with a capillary of the same diameter was used during the TR-SFX experiment at SACLA (see below).

**TR-SFX data collection at SACLA**

A serial femtosecond crystallography (SFX[9]) experiment was carried out at the Spring-8 Angstrom Compact Free-electron Laser (SACLA, Japan[10]) on beamline BL2-EH3 (proposal 2022B8815, 1 – 4 November 2022). *Tt*CBD microcrystals with a size of ~ 15 × 5 × 2 µm (Supplementary Fig. 5) were embedded in CMC-5% and extruded into a helium-filled *Diverse Application Platform for hard X-ray Diffraction in SACLA* (DAPHNIS[11]) chamber by a HVE injector[8] through a polyamide-coated glass capillary with an inner diameter of 75 µm at a flow rate at the pumping HPLC unit of 100 µl/min. At this flow rate, the jet speed had previously been determined to be around 5.2 µm/ms (Supplementary Fig. 34), ensuring a spacing of 172 µm between the centres of jet regions probed by two consecutive XFEL pulses (at 30 Hz XFEL repetition rate). SFX data were collected using XFEL pulses (< 10 fs in length, nominal photon energy 7.996 keV (FWHM 42 eV), photon flux of ~2 × $10^{11}$ photons/pulse, pulse energy of ~300 µJ at the sample position) focused to 1.6 µm (h) × 1.4 µm (v) (FWHM). Data were recorded using a MPCCD detector with eight sensor modules[12]. Sample handling, injector loading and data collection were carried out under red light conditions. The SFX experiments were carried out in a time-resolved mode (TR-SFX[13]) according to an optical pump – X-ray probe scheme. The pump laser (EKSPLA NT230, 530 nm, 5 ns pulse length (FWHM), circularly polarized) was aligned perpendicular to both the X-ray beam and the HVE jet and focused to a Gaussian spot with size of 105 (h) × 98.2 (v) µm² (FWHM). Diffraction data with (*light*) and without (*dark*) pump-laser excitation were collected in an interleaved way with the XFEL operating at 30 Hz and the pump laser at either 0 (*dark_only_SACLA*), 10 (*light-dark1-dark2* sequence), 15 (*light-dark*) or 30 Hz (*light_only*). Online monitoring of the hit-rate (i.e. the average percentage of images containing diffraction) was carried out using the data processing pipeline for serial femtosecond crystallography at SACLA[14].

A crystallographic pump-power titration was carried out in the first part of the beamtime to inform the choice of the pump laser fluence to be used in the subsequent time series. TR-SFX data were collected at a pump-probe delay of 3 µs (for the choice of this time delay see Supplementary Note 2) at 1.4, 3.5, 7, and 14 µJ laser pulse energy. These energies corresponded to fluences at the Gaussian peak of 12, 30, 60, and 120 mJ/cm², to power densities at the Gaussian peak of 2.4, 6, 12, and 24 MW/cm², and, assuming an average crystal size of 5 µm (~$(15 \times 5 \times 2)^{1/3}$ µm, see Supplementary Fig. 5), to nominally 1, 2.4, 4.8, and 10 absorbed photons on average per chromophore, respectively. Results of the crystallographic power titration are presented and discussed in Supplementary Note 2. Data were collected with a *light-dark1-dark2* sequence at 12, 30, and 120 mJ/cm², whereas at 60 mJ/cm² no interleaved dark data were collected (i.e. *light_only*).

A series of TR-SFX experiments was carried out at pump-probe delays of 10 ns, 300 ns, 100 µs, and 3 ms and at a pump laser fluence of 30 mJ/cm² chosen based on the results of the preceding crystallographic power titration (Supplementary Note 2). Data were collected with a *light-dark* sequence, since the *dark1* data set was shown to be free of light contamination in the preceding crystallographic pump-power titration (see below). Together with the data set at 30 mJ/cm² and 3 µs collected as part of the crystallographic pump-power titration, a time series consisting of five pump-probe delays was thus collected. A total of 120 mg of microcrystalline *Tt*CBD was used for the collection of all *dark* and *light* data sets.

#### Processing of TR-SFX data collected at SACLA

Hits identified during online monitoring were immediately indexed and intensities integrated using the CrystFEL software suite[15] version 0.10.1 (https://doi.org/10.5281/zenodo.10598358). Offline data processing was continued using CrystFEL (version 0.10.1). *Peakfinder8*[16] was used for peak finding with the following parameters: threshold=100, SNR=4, min-pix-count=2, and local-bg-radius=3. *Xgandalf*[17] and *Mosflm*[18] algorithms were used for indexing, and peak integration was performed using the *ring-nocen-nograd* method and *int-radius=3,4,7*. Intensities with a peak value below 14000 detector counts (max-adu) were merged using the *partialator* algorithm implemented in CrystFEL using two iterations of the unity model. The estimation of the sample-detector distance was optimized as described by Nass and coworkers[19]. A summary of hit- and success-rates as well as statistics indicators are reported in Supplementary Table 4 for the power titration data sets and in Supplementary Table 5 for the different time series data sets.

#### TR-SFX data collection at SwissFEL

A second TR-SFX experiment was carried out using the SwissMX instrument at the Cristallina experiment station[20] of SwissFEL (Villigen, Switzerland) (proposal 20231018, 7 – 10 June 2024), with pump-probe delays of 3 µs (ensuring temporal overlap with the SACLA time series) and 10 ms. *Tt*CBD microcrystals (~ 15 × 5 × 2 µm, Supplementary Fig. 5) were mounted on micro-structured polymer fixed targets (MISP chips[20], 30 × 30 $mm^2$) with 26244 (162 × 162) cavities (100 µm each in diameter) and a pitch of 120 µm. An opaque version of the chips was used in order to avoid pump-light contamination of neighbouring cavities in TR-SFX experiments. The cavities of a chip were loaded by pipetting 300 µl of a slurry containing 15% (v/v) settled *Tt*CBD microcrystals onto the surface under a constant stream of humid air. The excess solution was removed through the apertures (7 - 13 µm in diameter) by applying a vacuum. After covering both of their surfaces with layers of Mylar film (6 µm thickness), the chips were sealed and mounted on the SwissMX instrument. TR-SFX data were collected according to an optical pump – X-ray probe scheme, using X-ray pulses at a repetition rate of 100 Hz (25 - 45 fs in length, mean photon energy 11 keV, photon flux of $1.59 \times 10^{11}$ photons/pulse, pulse energy of ~280 µJ at the sample position) focused to 3.7 µm (h) × 4.2 µm (v) (FWHM) and recorded using a Jungfrau 8Mpixel detector[21]. Sample handling, chip loading and data collection were carried out under red light conditions. The pump laser (EKSPLA NT230, 532 nm, 3.5 ns pulse length (FWHM)) was coupled colinear to the X-ray beam and perpendicular to the chip plane. A 1 m optical fibre (100 μm core ID, Thorlabs) was used to couple the pump laser to the sample position and to shape the laser profile. The light at the fibre outcoupling was collimated and refocused through a two-lens system to a nearly top-hat spot. The $1/e^2$ width of the laser spot at the sample position was determined by a beam profiler (55 μm) and assumed to be the diameter of a top-hat circular beam. A pump-laser fluence of 30 mJ/cm$^2$ (0.75 μJ, 9 MW/cm$^2$ per pulse, corresponding to nominally 2.5 absorbed photons on average per chromophore assuming an average crystal size of 5 μm) was used to match the pump-laser excitation conditions used in the SACLA experiment (see section 'SFX data collection at SACLA'). Diffraction data with (*light*) and without (*dark*) pump-laser excitation were collected in an interleaved (*light-dark-light-dark*) sequence at pump-probe delays of 3 μs and 10 ms. For the 3 μs delay, a *continuous* data-collection strategy was used such that the sample stages did not stop moving but adjusted their velocity to the XFEL pulse in real time such that the aperture location was at the X-ray/laser interaction point when photons arrived. By comparison, a *stop-and-go* data-collection approach was used for the 10 ms delay. In this routine, the stages move an aperture into the X-ray/laser interaction point and then stop. Laser illumination occurs at a defined time prior to X-ray irradiation, after which the stages move to the next aperture and the process repeats itself. SFX data without pump laser (*dark_only_SwissFEL*) were collected as a reference data set to assess potential light contamination of the interleaved dark data acquired during the pump-probe data collection (see section *Assessing potential light contamination of interleaved dark images collected at SwissFEL*). Online monitoring of diffraction data, such as determination of hit-rate and estimation of the fraction of multiple hits, was carried out using the data processing pipeline for serial femtosecond crystallography at SwissFEL. A total of around 60 mg of microcrystalline *Tt*CBD was used for the collection of the three *dark* and the two *light* data sets.

**Processing of TR-SFX data collected at SwissFEL**

Online data processing was performed using the SwissFEL pipeline, which relies on CrystFEL version 0.10.2. The same CrystFEL version was also used for the offline data processing. The peak finding parameters were set to threshold=12, SNR=4.5 min-pix-count=1, and local-bg-radius=4 for the *peakfinder8* algorithm. Indexing relied on *Xgandalf*[17] and *Mosflm*[18], and the *ring-nocen-*

*nograd* method was used for integration, with integration radii of 3, 4 and 7 pixels for the peak, buffer and background regions, respectively. Intensities were merged using *partialator* (unity mode, two iterations), with a max-adu parameter of 2000. Detector distance was checked and optimized, and statistics of all data sets collected at SwissFEL are presented in Supplementary Table 6.

**SSX data collection on *Tt*CBD-H132A mutant microcrystals at Diamond Light Source**

Serial synchrotron crystallography (SSX) after laser excitation was performed on the *Tt*CBD-H132A mutant at beamline I24 of Diamond Light Source (Didcot, UK)[22]. Microcrystals of the *Tt*CBD-H132A mutant (~15 × 8 × 5 μm$^3$) were mounted on a glow-discharged silicon chip (Southampton Nanofabrication Centre, University of Southampton) with aperture size of ~10-12 μm. Chips were prepared under dim red-light conditions inside a humidity-controlled chamber. 150 μl of a slurry containing ~15% (v/v) settled *Tt*CBD-H132A microcrystals were deposited on the chip, and the excess crystallization solution was removed by applying a vacuum. The chip was then sealed with Mylar films (6 μm thick) on each side. Two data sets were collected at room-temperature, i) without optical-pump (*dark_DLS*) and ii) according to an optical pump – X-ray probe scheme. Using an X-ray beam of 12.4 keV focused to 8 μm (h) × 8 μm (v) (FWHM), each crystal was exposed for 10 ms with a photon-flux of 1.6 × 10$^{12}$ photons/second and diffraction data were recorded on a Pilatus3 6M detector (Dectris). The average diffraction weighted dose per crystal was calculated with Raddose-3D v5[23] to be 0.14 MGy. For the SSX data collection involving illumination, a portable pulsed laser system from Light Conversion (PORTO) was used to photoactivate the *Tt*CBD-H132A microcrystals. Mirrors and an achromatic focusing lens were used to focus the laser beam at the sample position, in such a geometry that the laser beam was ~15° off-axis from the X-ray beam with a laser beam size of 50 μm (FWHM) in both directions. The laser was operated at a wavelength of 515 nm and a repetition rate of 50 kHz, providing individual pulses of 300 fs. The laser intensity at the sample position was attenuated using a neutral density filter, yielding a laser power of ~0.8 mW; i.e. 0.016 μJ per 300-fs pulse. With this setup, pump laser excitation was performed during 10 ms (i.e. 500 pulses), corresponding to a pump laser energy of 8 μJ, a fluence of 320 mJ/cm$^2$ and to nominally ~12 absorbed photons per chromophore assuming an average crystal size of 8 μm. Diffraction images were collected 10 ms immediately after laser exposure. This dataset was collected with the pump-laser on for all apertures of the chip, i.e no dark-interleaved was collected. This dataset is referred to as *illuminated_DLS.*

**Processing of *Tt*CBD-H132A mutant SSX data collected at Diamond Light Source**

SSX data were reduced, scaled and merged using *xia2.ssx* from the DIALS software package (version 3.22.1[24] ). Default parameters were used apart from reducing the minimum spot size from 3 to 2 pixels for the detection of Bragg peaks. Dark-adapted model of *Tt*CBD-H132A mutant (PDB entry code: 8C35) was used to generate reference intensities for scaling. Statistics of the SSX dataset collected after illumination at I24 are presented in Supplementary Table 7.

**Calculation of Fourier difference maps**

To visualize photoinduced structural changes, q-weighted Fourier difference electron density maps ($F_o^{light} – F_o^{dark}$)[25] were computed with *Xtrapol8*[26]. q-weighted Fourier difference electron density maps were also computed between different nominal dark data sets to assess if light contamination occurred (see below).

**Dependency of integrated Fourier difference peaks as a function of indexed diffraction images at SACLA**

To help estimating the required number of indexed diffraction images (or lattices if there are more than one indexable patterns in a diffraction image) following light activation for a given experimental condition (pump laser fluence and time delay), the appearance and evolution of peaks in $F_o^{light} – F_o^{dark}$ maps were monitored as a function of the number of indexed lattices[27]. The sums of absolute values of positive and negative integrated peaks (up to $\pm 3\ \sigma$) in the chromophore region (within a distance of 2 Å from any chromophore atom) was computed for increasing numbers of indexed lattices. Based on the fitted parameters, after indexing ~ 70k light lattices, the increases of both the positive and negative signals in the $F_o^{light} – F_o^{dark}$ were small (only ~ 7% for 10,000 additional indexed light lattices). This number of indexed light lattices was therefore used as a guide throughout the experiment. As an example, this procedure has been repeated after the experiment for the 10-ns time delay (Supplementary Fig. 35) and is representative of those obtained during the beamtime. To generate the different data sets with fewer light lattices, random selection from the overall number of light lattices (86,000) was performed. To estimate the signal variability in the Fourier difference maps, 25 different data sets were composed for each number of light lattices sampled (from 10,000 to 80,000 in steps of 10,000 lattices).

**Assessing potential light contamination of interleaved dark images collected at SACLA**

A *dark_only_SACLA* data set without any pump-laser excitation (115k indexed lattices) was collected as a control to identify potential light contamination of interleaved dark data sets (*dark*, *dark1* and *dark2*). At each pump laser fluence used in the power titration at the 3 µs time delay, the Fourier difference electron density maps $F_o^{dark1} - F_o^{dark\_only\_SACLA}$ and $F_o^{dark2} - F_o^{dark\_only\_SACLA}$ were computed. In all these maps (contoured at ± 3.5 σ), peaks of similar height are randomly distributed all over the protein indicating no pump-laser contamination (Supplementary Fig. 7). *Dark* data sets from all time delays in the time series (Δt: 10 ns, 300 ns, 3 µs, 100 µs, and 3 ms) were also compared to the *dark_only_SACLA* data set. $F_o^{dark} - F_o^{dark\_only\_SACLA}$ maps resulted in a similar random distribution of positive and negative peaks, except for the 3 ms time delay where non-negligible peaks (up to 5 σ) in the $F_o^{dark} - F_o^{dark\_only\_SACLA}$ at the chromophore were observed (Supplementary Fig. 7). We concluded that the crystals corresponding to interleaved dark images of the 3-ms data set were light contaminated for unknown reasons. Consequently, these dark images were discarded. A large *reference dark* data set (*dark_ref_SACLA*, 847,084 indexed lattices, 2.05 Å resolution, Supplementary Table 5) was assembled from all remaining dark data sets (*dark_only_SACLA, dark, dark1* and *dark2*). Expectedly, a Fourier difference electron density map computed from the *dark_ref_SACLA* and *dark_only_SACLA* data sets ($F_o^{dark\_ref\_SACLA} - F_o^{dark\_only\_SACLA}$) only showed noise peaks distributed over the protein (Supplementary Fig. 7).

**Assessing potential light contamination of interleaved dark images collected at SwissFEL**

The strategy employed for the assembly of the dark dataset at SACLA has also been used for the dark dataset at SwissFEL. First, a *dark_only_SwissFEL* dataset was collected (~69k indexed lattices, Supplementary Table 6) as a reference to assess any potential light contamination of dark interleaved data (*dark*). Interleaved *dark* data sets from the 3 µs and 10 ms series were compared to the *dark_only_SwissFEL* dataset by means of a Fourier difference map ($F_o^{dark}$ – $F_o^{dark\_only\_SwissFEL}$). In these maps contoured at ± 3.5 σ, no sign of pump-laser contamination is observed as peaks are randomly distributed over the whole asymmetric unit tetramer and no peaks are specifically located on the AdoCbl (Supplementary Fig. 9). Therefore, a *dark_ref_SwissFEL* data set (151k indexed lattices) was generated using all available SwissFEL dark data sets (*dark_only_SwissFEL* and *dark* from the 3 µs and 10 ms series).

#### ***Dark_ref_SACLA* structure refinement**

The previously reported *Tt*CBD tetramer structure solved from form II crystals (PDB entry code: 8C73[2]) was used as a search model for molecular replacement with *PHASER*[28]. Iterative cycles of reciprocal space refinement performed using *REFMAC5*[29] from the CCP4 software suite[30] were interspersed with local real-space refinements and model building using *COOT*[31]. Ten cycles of refinement were performed for each *REFMAC5* run and consisted of restrained refinement of coordinates and isotropic atomic displacement parameters, as well as occupancy refinement of alternate conformers. Automatic B-sharpening of resulting $2mF_o$-$DF_c$ and $mF_o$-$DF_c$ electron density maps was applied in REFMAC5. This refined model was used for structure factor amplitudes extrapolation of TR-SFX data collected at SACLA.

Residual positive density was observed in the $mF_o$-$DF_c$ electron density map along large parts of the polypeptide chain of monomer D, which prompted us to perform ensemble refinement with *PHENIX*[32,33]. Values of *pTLS*, *tbath* and *tx* were sampled. For *pTLS*, which represents the fraction of the molecules included in the TLS fitting procedure, values of 0.6, 0.8, 0.9 and 1.0 were tested. *tbath* is a parameter coupled to the thermostat temperature of the simulation and controls as well the X-ray weight (2.5, 5 and 10 K have been tested). Finally, the relaxation time (*tx)* used during the simulation has been tested with values of 0.35, 0.7 and 1.4 ps. The lowest $R_{free}$ value of these refined ensembles was chosen as a criterion to determine the most appropriate set of values of these three parameters.

***Dark_ref_SwissFEL* structure refinement**

The previously refined *dark_ref_SACLA* model was used as a search model for molecular replacement with *PHASER*[28]. The subsequent refinement procedures were identical to those described in the section above. Similar $mF_o$-$DF_c$ peaks were observed in monomer D as reported previously for the *dark_ref_SACLA* data set; no additional ensemble refinement was performed for this data set.

***Dark_DLS* refinement of H132A mutant structure**

Molecular replacement was performed with *PHASER*[28] using as a search model the structure of the dark-adapted *Tt*CBD-H132A mutant (PDB entry code: 8C35). The refinement procedure was similar to the one described in the *Dark_ref_SACLA structure refinement* section above. This model is referred to as *dark_DLS* model.

**Structure factor amplitudes extrapolation of TR-SFX data sets**

We resorted to scalar structure factor extrapolation[34] to determine the structures of intermediate states at all time delays from experiments at both SACLA and SwissFEL. *Xtrapol8*[26] (version 1.0.0) has been used for the computation of extrapolated structure factor amplitudes as follows:

$$\text{(Eq. 1)}\ F_{extr} = \frac{q}{\langle q \rangle} \times \alpha \times (F_o^{light} - F_o^{dark}) + F_o^{dark}$$

*q* is the *q*-weighting factor, $\alpha$ is the extrapolation factor (> 1) which needs to be determined. In this work, we assume the occupancy of the light state to be equal to $1/\alpha$. Here, a newly developed method (Jacques-Philippe Colletier, personal communication) based on Singular Value Decomposition (SVD) has been used. In short, the $2mF_o$-$DF_c$ map of the appropriate *dark_ref* data set (*dark_ref_SACLA* or *dark_ref_SwissFEL*), as well as extrapolated $2mF_{extr}$-$DF_c$ maps computed for discrete and increasing α values are used to generate the matrix to be decomposed (each matrix column corresponds to a unique set of flattened electron density grid points). Performing the decomposition on such a matrix always yields the electron density map of the *dark* state as the first vector. The α value at which the computed extrapolation is the most meaningful is obtained when the amplitude of the dark state (i.e. *first left singular vector*) is minimal. This method yielded α values of 3.3 for data sets *10ns_12mJ/cm²_SACLA, 10ns_30mJ/cm²_SACLA* and *3µs_12mJ/cm²_SACLA* (corresponding to an occupancy of 30 %) and 3.6 for data sets *300ns_12mJ/cm²_SACLA, 3µs_30mJ/cm²_SACLA, 100µs_30mJ/cm²_SACLA* and *3ms_30mJ/cm²_SACLA* (28% occupancy). For the different time delays collected at SwissFEL, α values of 2.5 (40% occupancy) were obtained. Of note, a single α value is used in extrapolation (Eq. 1), meaning the same occupancy of the light state is assumed for all four monomers, which might not be the case. Negative extrapolated structure factors have been rescued via the *truncate* method implemented in *Xtrapol8*.

**Refinement of TR-SFX intermediate-state structures**

First, all models were refined with *REFMAC* as described for the *dark_ref_SACLA* and *dark_ref_SwissFEL* models (see above) against q-weighted extrapolated structure factor amplitudes. Further refinement was then performed with *phenix.refine*[32]. In addition to the global refinement parameters, coordinates were refined in both real- and reciprocal space, atomic displacement parameters (ADPs) were individually refined, and refinement of occupancies was also performed. To properly refine models with a distance between the cobalt atom and the C4′ atom of the adenosyl group (Co–C4′) shorter than the sum of the van der Waals radii (3.2 Å), a custom bond was generated to be used in *phenix.refine*. As the ideal bond distance was unknown, we tested different values (here 2.0, 2.3 and 2.9 Å) as well as different standard deviations (sigma values) to allow deviation from the ideal restrained distance (0.1, 1 and 10 Å). These different trials showed that with a sigma of at least 1 Å, the ideal bond distance was of little importance as refinements yielded similar Co–C4′ distances (within 0.1 Å). Such settings allow us to turn off the van der Waals repulsion while not imposing any strong restraints on the resulting bond length. We therefore performed all our refinements with an ideal bond length of 2.9 Å and a sigma value of 1 Å for the Co–C4′ distance. Refinement statistics of all intermediate-state models are summarized in Supplementary Tables 4-6.

**Refinement of SSX H132A mutant structures**

Following the same methodology applied with the TR-SFX data sets of WT *Tt*CBD, we first resorted to the computation of a Fourier difference map based on the *dark_DLS* and the *illuminated_DLS* data sets. The overall $R_{iso}$ was unexpectedly high (19.3%), yielding uninterpretable Fourier difference maps that prevented structure factor extrapolation. In comparison, the overall $R_{iso}$ values for all TR-SFX data sets collected at SACLA range from 8.6% to 11.3%. An alternative strategy was therefore used to model and refine the intermediate-state structure of the *illuminated_DLS* dataset. As a first step, a $mF_o$-$DF_c$ polder map was computed using the *illuminated_DLS* dataset and the *dark_DLS* model with the adenosyl moiety omitted. This map unambiguously showed that for a majority of crystalline molecules the Co–C5′ bond is broken and the adenosyl group adopts a different conformation to the one observed in the dark model (Supplementary Fig. 36). The $mF_o$-$DF_c$ polder map was used to fit the conformation of the adenosyl moiety in the intermediate state (referred to as *10ms_DLS* adenosyl conformation).

To estimate the occupancy of the intermediate-state model, a composite model was made with a single conformer for the protein chain corresponding to the dark-adapted state and two alternate conformers for the adenosyl group, i.e. the one of the adenosyl moiety in the intermediate state (see above; *10ms_DLS* adenosyl conformation) and a second corresponding to the dark-adapted state (extracted from the *dark_DLS* model). This model was then subjected to reciprocal-space refinement against *illuminated_DLS* structure factors with *phenix.refine*[32] during which for the protein chain only the isotropic Atomic Displacement Parameters (ADPs) were refined, while for the two adenosyl conformers atomic positions, ADPs and occupancies were refined. This refinement yielded occupancies for the *10ms_DLS* adenosyl conformation of 0.81, 0.89, 0.97 and 0.81 for monomers A, B, C and D, respectively.

To obtain a complete intermediate-state model (i.e. comprising both protein and adenosyl moieties), another composite model for refinement against *illuminated_DLS* structure factors was generated as follows. Conformer A corresponded to the *dark_DLS* model with occupancies set to 0.20, 0.10, 0.05 and 0.20 for all atoms in monomers A, B, C and D, respectively. Conformer B contained the protein chains and the cobalamin groups of the *dark_DLS* model and the adenosyl group in the *10ms_DLS* adenosyl conformation. Occupancy of all conformer B atoms was set to 0.80, 0.90, 0.95 and 0.80 for monomers A, B, C and D, respectively. Water molecules were added to conformer B with an occupancy of 1. Subsequently, reciprocal-space refinement was performed with *phenix.refine*[32]. First, only the atomic positions of conformer B were refined, while the atomic positions of conformer A remained fixed. Then, refinement of isotropic ADPs was performed for all atoms in conformers A and B. Occupancies of atoms in conformers A and B were kept at their set values (see above) and not refined. The custom-defined Co–C4′ bond was also used for conformer B (see section above "Refinement of TR-SFX intermediate-state structures"). Refinement statistics are summarized in Supplementary Table 7.

**Coordinate error estimation based on the bootstrapping method**

The *bootstrapping* resampling approach can be used in serial crystallography to estimate the precision of atomic coordinates when individual observations are varied[27]. Individual observations correspond here to the indexed diffraction patterns, which serves as a pool to constitute resampled data sets. In the bootstrapping method, a given indexed pattern can be drawn multiple times (random drawing with replacement) and the resampled data sets contain the same number of indexed patterns as the original one. All TR-SFX data sets (*dark_ref_SACLA*, *dark_ref_SwissFEL* and all *light*) have been resampled 100 times to compute for each time delay 100 resampled sets of extrapolated structure factor amplitudes with the appropriate α value (3.3 and 3.6 for the different time delays collected at SACLA and 2.5 for the ones collected at SwissFEL). Then, 100 refinements were performed with *phenix.refine* for each intermediate state using the corresponding refined model as starting model. All atom coordinates were reshuffled by 0.3 Å using the sites.shake option in *phenix.refine*. All coordinate errors given in the main text and the Supplementary Note 3 have been estimated by the bootstrapping method. For the distances, errors were computed using the RMSD of the 100 distances obtained from the 100 refined models.

**Calculation of difference distance matrices (DDM)**

Intramolecular difference distance matrices and their representations were generated using the *ddm.py* script available in *Xtrapol8* (using only $C_\alpha$ atoms). These are shown in Extended Data Fig. 6 and in the main text Fig. 2d.

#### Cryo-temperature dependent macromolecular crystallography and *in crystallo* UV-vis spectroscopy

Cryo-temperature dependent macromolecular crystallography (MX)[35-37] in combination with *in crystallo* UV-vis absorption microspectrophotometry[38] on the BM07-FIP2 beamline of the European Synchrotron Radiation Facility (ESRF)[39] was used to characterize structural and spectroscopic changes in crystalline *Tt*CBD after light illumination, respectively. Macrocrystals of *Tt*CBD (form I)[2] were produced with the sitting drop method at room temperature (20 °C) and under red light conditions as described earlier[2]. Crystals reached their final size (50 × 100 × 250-300 $\mu m^3$) overnight. They were transferred to a drop containing mother liquor (form I) and 25% (w/v) PEG 200 as a cryo-protectant and flash-cooled in liquid nitrogen. For spectroscopic and X-ray data collection, crystals were mounted in the gaseous nitrogen stream of a cryostream (Oxford Cryosystems 1000) operating at 100 K on the BM07-FIP2 beamline equipped with an online UV-vis microspectrophotometer (Supplementary Fig. 37).

Online UV-vis absorption spectra were collected under red light conditions on dark-state and illuminated crystals with a microspectrophotometer (Ocean Optics, QEPRO) connected to the lower 4× demagnifying objective with a 600 µm (ID) optical fibre and a deuterium - tungsten halogen light source (Ocean Optics DH2000 BAL) connected to the upper 4× demagnifying objective with a 400 µm (ID) optical fibre (Supplementary Fig. 37). Illumination of the crystals was performed at different cryo-temperatures with an optical fibre-coupled 530-nm LED (Thorlabs) through a 1000 µm fibre connected to the upper objective. The power measured at the fibre exit was 24.7 mW. To ensure homogeneous illumination, crystals were rotated around the diffractometer spindle axis while being illuminated. At 100 K, constant illumination for 40 min was necessary to fully photoconvert *Tt*CBD (Supplementary Fig. 20a). At 180 K, 6 min were sufficient (Supplementary Fig. 20b). With the aim to generate and structurally characterize photointermediates at temperatures above 100 K, we increased the temperature of a crystal from 100 K to either 150, 180, 200 or 210 K at 360 K/h, followed by illumination as described above for as long as the spectrum still changed. The temperature was then lowered back to 100 K at 360 K/h. Below, X-ray data collection is described only for a crystal after temperature excursion to 180 K. Without temperature excursion (i.e. illumination at 100 K) and after excursion to and illumination at 150 K, structural changes in the chromophore region were observed (not shown), but not to the extent of the changes obtained with illumination at 180 K. Increasing the temperature

to either 200 or 210 K led to crystalline ice formation as the solvent glass transition was passed[40] and, consequently, to degradation of diffraction quality.
X-ray crystallographic data were recorded on a Pilatus 6M detector (DECTRIS) at 100 K on a *Tt*CBD crystal previously activated by illumination at 180 K (6 min, monitored by UV-vis spectroscopy (Supplementary Fig. 20b)). The X-ray beam (photon energy 12.657 keV, photon flux $4.7 \times 10^{11}$ ph/s) had a size of 250 µm × 250 µm at the sample position. 1500 images were collected with a 0.2° oscillation range and an exposure time of 150 ms each. An average diffraction weighted dose of 1.1 MGy per data set was calculated using Raddose-3D v5[23]. The diffraction intensities were integrated, scaled, and merged using XDS (version 20230630). The previously reported tetrameric model of *Tt*CBD (PDB entry code: 8C31[2]) was used as a search model for molecular replacement. Based on spectroscopy, we assumed all crystalline molecules were fully light-activated. Therefore, we included only one conformation in the model and refined it at 1.9 Å resolution by iterative cycles of reciprocal space refinement using both *REFMAC5*[29] from the CCP4 software suite[30] and phenix.refine[32]. Refinement of the Co–C4′ distance was performed as previously described for the refinement of the time-resolved intermediate-state structures (see section *Refinement of intermediate-state structures*). The bond parametrization was set to a length of 2.6 Å and sigma value of 1 Å. The refined model is referred to as *cryo_light_180K*. Electron density maps ($2mF_o$-$DF_c$ and $mF_o$-$DF_c$_polder) are shown in Supplementary Fig. 21. Local real space refinement and model building were carried out in *COOT*[31]. Data collection and refinement statistics are reported in Supplementary Table 8.

**Low temperature cw-EPR on *Tt*CBD after steady-state illumination at 180 K**

For electron paramagnetic resonance (EPR) spectroscopy, two samples were prepared, one under aerobic and the other under anaerobic conditions. For aerobic conditions, 200 µl of dark adapted *Tt*CBD at ~ 250 µM was added into 4 mm outer diameter/3 mm inner diameter Suprasil quartz EPR tubes (Wilmad LabGlass), flash-cooled in liquid nitrogen, illuminated at 180 K and EPR spectra collected at 20 K (Supplementary Fig. 22a). Additional EPR spectra were also collected after increasing the temperature to above 180 K where indicated (Supplementary Fig. 22b).
The sample under anaerobic conditions was prepared identically, except that it was first moved inside an anaerobic glove box to equilibrate in an $O_2$ deficient environment for 48 hrs. Subsequently, 200 µl of this sample was transferred to 4 mm outer diameter/3 mm inner diameter

Suprasil quartz EPR tubes inside the glove box and tubes were sealed, illuminated for 15 secs at room temperature, left in the dark for 1 hour to ensure full conversion from Co(I) to Co(II), flash-cooled in liquid $N_2$ outside the glove box and EPR spectra collected at 20 K (Fig. 3F).

The photoactivation was carried out using a Thorlabs Mounted LED (M530L3) emitting at 530 nm with the output beam collimated with a Thorlabs collimation adaptor (SM2F32-A). The nominal output power (370 mW) was maintained by driving with a constant current of 1 A from a Thorlabs LED Driver. All EPR samples were measured on a Bruker EMXplus EPR spectrometer equipped with a Bruker ER 4112SHQ/ X-band resonator. Sample cooling was achieved using a Bruker Stinger[41] cryogen free system mated to an Oxford Instruments ESR900 cryostat and temperature control was maintained using an Oxford Instruments MercuryITC as reported previously[42,43]. The optimum conditions used for recording the spectra are as follow: microwave power 20 dB (2.19 mW), modulation amplitude 5 G, time constant 82 ms, conversion time 16.7 ms, sweep time 60 s, receiver gain 30 dB and an average microwave frequency of 9.385 GHz. The processing and analysis of the continuous wave EPR spectra were performed using the EasySpin toolbox (5.2.36) for the Matlab program package[44].

**Preparation of sample for time-resolved X-ray solution scattering (TR-XSS)**

*Tt*CBD was purified following a protocol similar to that outlined above. SEC was performed using 50 mM Tris-HCl pH 7.5, 100 mM NaCl as an eluent. Fractions containing *Tt*CBD were pooled and concentrated to 16 mg/ml (~700 μM). To produce full length *Tt*CarH a previously published protocol was used[1]. Full length *Tt*CarH was concentrated to 5.5 mg/ml (165 μM). To generate a DNA bound *Tt*CarH sample, a double stranded 26 bp long[45] DNA segment was synthesized (Integrated DNA Technologies) and incubated in the ratio of 1:1.5 (purified protein: DNA concentration) at room temperature for 2 hrs. The sample was then passed through a gel filtration column to remove excess DNA. Fractions containing DNA bound protein were pooled and concentrated to 8 mg/ml (240 μM). Samples were flash-cooled and kept at -80° C until use in TR-XSS experiments. Just before TR-XSS experiments, samples were thawed and the concentration adjusted at the values reported in the SI section dedicated to TR-XSS experiments.

**TR-XSS: data collection and reduction**

TR-XSS data were collected at the ID09 beamline of the ESRF[46,47]. The protein sample (*Tt*CBD in 50 mM Tris-HCl pH 7.5, 100 mM NaCl, 20 °C) was photoexcited with a 5 ns pulsed laser (EKSPLA NT342B) at 532 nm. The laser was focused with cylindrical lenses to an elliptical spot approximately 2.0 × 0.2 $mm^2$ (FWHM) corresponding to a fluence of ~50 $mJ/cm^2$ and ~2.5 absorbed photons/chromophore. X-ray (14.7 keV) pulse trains 10 or 80 µs long were selected using the ID09 chopper system[47] and probed the sample at 90 degrees with respect to the laser incoming direction. The 30 µs - 3 s time delay range was explored. Scattering patterns were recorded on a Rayonix MX170-HS detector placed 358 mm away from the sample and azimuthally integrated to obtain 1-dimensional scattering signals S(q) in the 0.05-2.4 $Å^{-1}$ q-range. TR-XSS difference signals ΔS(q) were obtained as the difference between the signals measured after photoexcitation minus the signal measured before photoexcitation. Several pump-probe sequences were collected on each sample and the ΔS(q) signals reported in the main text (Fig. 5) are typically the average over ~ 100 different repetitions. In addition, an independent measurement of a dye in solution was used as a reference curve for the solvent heating signal[48], which was later subtracted from the data to retain only the structural signal originating from the protein (Supplementary Fig. 29). All the above data reduction processing steps were performed using the *txs* python package developed at ID09 (https://gitlab.esrf.fr/levantin/txs).

Two different TR-XSS data sets collected on *Tt*CBD are reported in the main text (Fig. 5). The first data set (time delay from 30 µs to 100 ms) was obtained by flowing the sample (*Tt*CBD at ~ 16 mg/ml) through a 1.5 mm ID quartz capillary in a closed loop with a peristaltic pump (Gilson, Minipuls). The flow rate was adjusted to ensure that, at each time delay: (1) the sample cleared from the X-ray probed region in between consecutive laser pulses; (2) the sample moved slowly enough to ensure that the X-ray pulse probed a region well within the laser photoexcited region within the longest investigated time delay.

The second data set reported in the main text (time delays from 100 ms to 3 s) was obtained by flowing the sample (*Tt*CBD at ~ 4 mg/ml) through a 1 mm quartz capillary with a syringe pump (Chemyx, Fusion 4000X). Once a fresh sample volume reached the capillary, the sample flow was stopped and the scattering pattern from a single laser (pump) / X-ray (probe) pulse pair was collected. The volume illuminated by the laser was then flushed away between each measurement by the syringe pump before the next scattering pattern was collected. This second data set was collected on a sample at a protein concentration four times lower than the first data set to ensure that the protein-chromophore concentration was approximately half of the oxygen concentration in solution (thus ensuring fully aerobic conditions allowing completion of the photoreaction up to tetramer to monomer dissociation[2,5]. A comparison of the 100 ms TR-XSS signal from the two reported data sets (Supplementary Fig. 26) demonstrates that the different data collection protocols described above resulted in TR-XSS signals having the same shape. A protocol analogous to that used for the second data set (time delays from 100 ms to 3 s) was also used to collect TR-XSS data on *Tt*CarH and a *Tt*CarH-DNA complex in the time interval 100 ms - 3 s (Fig. 5g in the main text and Supplementary Fig. 28).

#### Singular value decomposition of TR-XSS data

Since photoactivation of *Tt*CBD is an irreversible process, the photoactive fraction of the sample (i.e. the part of the sample that can be still photolysed) decreases during the data collection, leading to a progressive reduction in the TR-XSS signal amplitude along the collection. To correct for this effect, all signals corresponding to the same time delay between X-ray and laser pulses were analysed by singular value decomposition (SVD)[49]. SVD analysis was applied on each data set relative to a given time delay in a data collection run, which is structured as a 2D data array containing the difference scattering signals ΔS(q) computed for all the images. The SVD results in a decomposed signal $A = USV^T$: U is a $\mathbf{n} \times \mathbf{m}$ matrix containing the left singular vectors (lSV, basis patterns), **n** being the number of momentum transfer q-values measured and **m** the number of repetitions; S is a $\mathbf{m} \times \mathbf{m}$ matrix containing the singular values; V is a $\mathbf{m} \times \mathbf{l}$ matrix, **l** being the number of repetitions, that contains the amplitude (right singular) vectors (rSV). rSV vectors describe the temporal evolution of the photoactivatable fraction in the sample during the acquisition of consecutive images. The basis patterns presented a temporal decay which could be modelled with a single exponential thus allowing to easily correct the data for the reduction of the sample photoactive fraction (Supplementary Fig. 30).

The SVD was also used to analyse the kinetic behaviour of the TR-XSS data set (dependence on the time delay after photoexcitation). In the case of the 30 µs - 100 ms data set two basis patterns were enough to accurately reproduce the time evolution of the data in the entire investigated time range (see Fig. 5 in the main text and Extended Data Fig. 8). For the 100 ms - 3 s data set, only one basis pattern was required to reproduce the time evolution of the data (see Fig. 5 in the main text and Supplementary Fig. 28).

**XSS difference signals calculated from PDB files**

The S(q) scattering signal corresponding to each *Tt*CBD crystallographic structure was calculated using the CRYSOL software[50] from the ATSAS package[51] on the PDB files obtained by X-ray crystallography. Difference signals ΔS(q) were then calculated using the signal from the *Tt*CBD dark structure (PDB entry code: 8C73 and the dark-state structure determined by SFX (*dark_ref_SACLA*) for Fig. 5c and Fig. 5d, respectively) as a reference for the resting state.

The atomic coordinates of the previously reported corrin-ring-shifted *Tt*CBD structure (PDB entry code: 8C76[2]) and the dark-state *Tt*CBD structure (PDB entry code: 8C73[2]) were used to compute the XSS difference signal reported in Supplementary Fig. 25. A second XSS difference signal was computed using the atomic coordinates of the 10 ms model from SwissFEL (*10ms_30mJ/cm²_SwissFEL*) and the corresponding dark reference model (*dark_ref_SwissFEL*) and reported in Supplementary Fig. 25. The signals simulating *Tt*CBD dimers and monomers (Supplementary Fig. 27) were calculated using the coordinates of the AB dimer or the A monomer (PDB entry code: 8C73[2]), respectively, were retained.

A hybrid model was generated from the tetrameric dark state *Tt*CBD model (PDB entry code: 8C73) and from the model with a shifted corrin ring observed after illuminating a macrocrystal for 5 s at room temperature (PDB entry code: 8C76[2]). The hybrid model was generated by using the PDB 8C73 as a reference for the protein atoms. PDB 8C76 was then superimposed to PDB 8C73 by minimizing the distance between their respective $C_\alpha$ atoms using VMD[52]. Finally, the AdoCbl coordinates from PDB 8C73 were replaced by those from PBD 8C76. The resulting structure is referred to as 8C73* in Fig. 5c of the main text.

**Cluster-model calculations**

We used Density Functional Theory (DFT) cluster models to examine the mechanism for the formation of a Co–C4′ adduct and explore the inherent differences in the chemistry of the Co–C5′ and Co–C4′ adducts. These calculations were performed in Gaussian16 rev. D[53] using the BP86 functional[54] which has been shown to perform well for $B_{12}$ systems[55,56], with the Def2-TZVP basis set on cobalt and the 6-31G(d,p) basis sets on the remaining atoms, an implicit solvation model

(CPCM) with dielectric constant of 5.0 and empirical dispersion (GD3BJ[57,58]). The cluster model was constructed from the *dark_DLS* model of the H132A mutant (monomer A) and consisted of a truncated Cbl, adenosyl, and binding-site residues W131 and H177 truncated as $-CH_3$ at the Cβ and V138, E141 and H142 truncated as CHO and $NH_2$ C- and N- termini, respectively (Supplementary Fig. 38) for a total of 211 atoms, with a net charge of 0. Six atoms were kept fixed during the calculations: the Cβ of W131 and H177, the Cα of V138, E141 and H142 and the Co. Comparison with the WT crystal structure (*dark_ref_SACLA*) revealed that the fixed atoms could be overlayed with the root mean square deviation (RMSD) of 0.1 Å. The potential energy profile for the interconversion between Co–C4′ adduct, photoproduct and Co–C5′ adduct (Supplementary Fig. 23) was calculated using relaxed potential energy scans, and all transition states were fully optimized and frequency calculations confirmed the presence of a single imaginary frequency.

#### QM/MM setup and simulations

The dark and intermediate state crystal structures (*dark_ref_SACLA*, *10ns_30mJ/cm²_SACLA*, and *3µs_30mJ/cm²_SACLA*) were used to create computational models to corroborate the assignments at the time-steps prior to the formation of the light state. For each model, a monomeric subunit of the tetramer was used. The model systems were minimized using the ff14sb AMBER force field[59] and the hydrogens were added with Amber16[60]. Subsequently, the models were studied using the quantum mechanics/molecular mechanics (QM/MM) approach. The AMBER parameters for coenzyme-$B_{12}$ were obtained from Marques, *et al.*[61]. The models were divided into two-layers namely the high- and low-layer which were treated with QM and MM methods, respectively. The high-layer, also called the QM region, was comprised of a portion of the cobalamin, the adenosyl group and the imidazole portion of H177 (Supplementary Fig. 39). A hydrogen link atom was placed at the sp3 carbon that was bound to the amide nitrogen of the nucleotide loop and the majority of the nucleotide loop including the dimethylbenzimidazole were treated in the MM region. The corrin ring, cobalt ion, and the corrin ring side chains were included in the QM region (see the grey, red, blue, and pink atoms in Supplementary Fig. 39). The upper 5′-deoxyadenosyl ligand and the imidazole portion of the lower ligand, H177, were also placed in the high layer. The H177 backbone atoms were in the low layer also called the MM region. The QM region was treated with density functional theory (DFT) using the BP86 functional along with the TZVP basis set for H and the TZVPP basis set for Co, C, N, and O[62-66]. In total the QM region contained 170 atoms.

In addition to the portion of the nucleotide loop along with the dimethylbenzimidazole of cobalamin all the amino acids were considered in the MM region. Any residue which had at least one atom within 10 Å of the cobalt was kept unfrozen, meaning it was allowed to freely move throughout the course of the optimizations. The QM/MM models were optimized using Gaussian09[67].

#### Supplementary Notes

##### Supplementary Note 1. Time-resolved absorption spectroscopy of *Tt*CBD in solution and in microcrystals

Time-resolved absorption measurements of *Tt*CBD in solution (Extended Data Fig. 2) show similar spectral intermediates and kinetics of formation / decay to those reported for full-length *Tt*CarH[68]. Difference absorbance spectra recorded at 10 ns and 3 µs after ns laser excitation at 530 nm (Extended Data Fig. 2a) show that an initial spectral change (<3 ns time constant[68,69]) leads to the formation of a new spectral intermediate state within 3 µs. The new spectral species at 3 µs is characterized by a decrease in the absorbance bands at 386 and 572 nm, and simultaneous increase in absorbance features at 334 and 502 nm (Extended Data Fig. 2a). Kinetic absorbance transients at 500 nm show that this species is formed with a time constant of 470 ± 60 ns (Extended Data Fig. 2b), which is in agreement with the kinetics reported for the full-length protein[68]. To investigate the accumulation of any further intermediate, prior to the formation of the final light-adapted state, a difference absorbance spectrum was recorded 50 ms after laser excitation (Extended Data Fig. 2c). The difference spectrum indicates the formation of a new intermediate state that involves a number of absorbance changes, including increases at 353 nm and between 470 and 540 nm and further decreases at both 380 and 568 nm (Extended Data Fig. 2c). To measure the kinetics of formation of this new state the rate of absorbance increase at 360 nm was measured upon laser excitation at 530 nm, yielding an associated time constant of 7.3 ± 0.3 ms (Extended Data Fig. 2d). This species is converted into the final light-adapted state, as shown by a further increase and red-shift of the peak at 353 to 359 nm (indicated by the 5 s spectrum in Extended Data Fig. 2e). This spectral change is indicative of the bis-histidine ligated light-adapted state[2] and is formed with an associated time constant of approximately 0.59 ± 0.03 s (Extended Data Fig. 2f).

The time-resolved absorption spectroscopy measurements lead us to propose a sequential reaction pathway for the photoactivation of *Tt*CBD, together with the corresponding time constants for each of the respective steps (Extended Data Fig. 2g). Intermediate states 1-3 in Extended Data Fig. 2g correspond to the species *ii* - *iv* in Fig. 1 of main text.

To investigate whether the same reaction pathway occurs in the crystalline state we repeated the time-resolved absorption spectroscopy measurements on 20 % (v/v) *Tt*CBD microcrystal slurries in CMC-4% (4% CMC, 10% PEG 10000 and HEPES 0.05 M pH 7.5) (Supplementary Fig. 2). Despite significantly lower signal-to-noise levels due to the large amount of scattering of the probe light from the crystals, these measurements (Supplementary Fig. 2) yielded similar rates of formation for intermediates 2 and 3 as *Tt*CBD in solution (Extended Data Fig. 2). The rate of formation of intermediate 2 was measured by following the absorbance change at 334 nm (instead of 500 nm as in Extended Data Fig. 2), due to the larger absorbance increase at this wavelength, with a respective time constant of 540 ± 90 ns in solution (panel A) compared to 520 ± 120 ns *in crystallo* (panel B). The rate of formation of intermediate 3 was determined by measuring the absorbance change at 460 nm (instead of 360 nm as in Extended Data Fig. 2). This probe wavelength was used for *in crystallo* measurements as it minimizes the photoconversion in the crystals, whilst maximizing the signal-to-noise-ratio, and yielded time constants of 6.5 ± 0.8 ms in solution (panel C) and 6.4 ± 0.9 ms *in crystallo* (panel D). As described previously[2] the slower completion of the light-adapted state, associated with the dissociation of the *Tt*CBD tetramer, causes disintegration of the crystals. Therefore, spectral changes accompanying the formation of the bis-histidine ligated light-adapted state were not accessible in crystals. However, the photoproduct was identified using LCMS (Supplementary Fig. 3) and NMR (Supplementary Fig. 4) from microcrystals exposed to light, confirming the microcrystals are photoactive and capable of producing 4′,5′-anhydroadenosine.

**Supplementary Note 2. Spectroscopic and crystallographic pump power titration**

In preparation for the ns TR-SFX experiments described in this work, spectroscopic and structural pump laser power titrations have been carried out on *Tt*CBD microcrystals to select the appropriate pump-laser fluence to be used. Indeed, if a high fluence is chosen, in particular for fs excitation, multiphoton effects might alter the peaks and their heights in Fourier difference maps and change the kinetics of structural changes with respect to the functionally meaningful single-photon excitation[70]. When using ns excitation, as in the present study, multiphoton effects are unlikely to occur when excitation is carried out with nominally more than one absorbed photon per chromophore per pulse, yet sequential photon absorption by light-induced intermediate states can still result in the population of non-functional states[71]. The results of spectroscopic and structural pump laser power titrations on *Tt*CBD microcrystals are presented and discussed in the following. Furthermore, care has been taken that the size of the used *Tt*CBD microcrystals (~ 15 × 5 × 2 µm (Supplementary Fig. 5) does not exceed in any dimension the 1/e penetration depth at 530 nm (17 µm; Supplementary Fig. 40) to ensure efficient and homogeneous photoexcitation at low laser fluences.

Due to the significantly lower signal-to-noise levels caused by the large amount of scattering of the probe light from the crystals it was not possible to measure transient spectra by time-resolved absorption spectroscopy on microcrystals as a function of pump-laser fluence. Consequently, static UV-vis spectroscopy was used to perform a spectroscopic power titration on crystals. Stationary spectral changes were measured on microcrystals of tetrameric *Tt*CBD embedded in CMC-4% following 6-8 ns laser excitation with a single pulse at 530 nm at different pump-laser fluences (Extended Data Fig. 3). In particular, the change in absorbance at 359 nm, which reports directly on the accumulation of the bis-histidine ligated light-adapted state (Poddar 2023), was monitored after a single-pulse excitation at laser energies between 6 and 60 mJ/cm² (corresponding to 0.4 and 4.8 nominally absorbed photons per chromophore, respectively; Extended Data Fig. 3). The amount of light-adapted state formed increases linearly up about to 20 mJ/cm² (corresponding to 1.6 nominally absorbed photons per chromophore) and reaches a plateau of about 60% formation of the light-adapted state from a single laser pulse at approximately 30 mJ/cm² (corresponding to 2.4 nominally absorbed photons per chromophore). To assess the effect of laser fluence on photodynamics of *Tt*CBD in solution, kinetic absorbance transients at 334 nm were measured on the µs timescales at laser fluences ranging from 6 to 60 mJ/cm² (corresponding to 0.4 and 4.8 nominally absorbed photons per chromophore; Extended Data Fig. 3). Single exponential fits yielded characteristic time constants of approximately 600 ns with little dependence on the pump-laser fluence (Extended Data Fig. 3c). However, the magnitude of the absorbance increase at 334 nm (Extended Data Fig. 3d) was linearly-dependent on the pump-laser fluence up to about 20 mJ/cm² (corresponding to 1.6 nominally absorbed photons per chromophore) and levelled off at around 30 mJ/cm² (corresponding to 2.4 nominally absorbed photons per chromophore), as for CMC-embedded *Tt*CBD microcrystals. From the spectroscopic power titrations on the ns to µs time scale, we conclude that the photochemical yield does not increase any further at fluences corresponding to nominally more than 2.4 absorbed photons per chromophore and that the photokinetics are largely independent of pump-laser fluence in the range examined.

The aforementioned spectroscopic power titration was then followed by a crystallographic power titration at SACLA to inform the decision on the adequate pump laser fluence to be used in the subsequent TR-SFX experiment at SACLA at various pump-probe delays. The study presented here is the first TR-SFX experiment on a $B_{12}$-dependent photoreceptor protein. Consequently, no prior knowledge was available on peaks and their heights to be expected in Fourier difference maps at different time delays. Based on the reaction scheme proposed above from the time-resolved absorption spectroscopy measurements (Extended Data Fig. 2g), we reasoned substantial structural changes corresponding to a fully populated intermediate might be observed at 3 µs, a delay corresponding to > 5× the characteristic time constant (~500 ns) of conversion from intermediate 1 (species *ii* in Fig. 1 of the main text) to intermediate 2 (species *iii* in Fig. 1 of the main text) but still ≪ 6-8 ms, i.e. the time constant of the successive spectral intermediate (Extended Data Fig. 2g). In the crystallographic power titration, hence carried out at a pump-probe delay of 3 µs, TR-SFX data were collected in a *light-dark1-dark2* sequence, at pump-laser fluences at the Gaussian peak of 12, 30, 60, and 120 mJ/cm², corresponding to peak power densities at the Gaussian peak of 2.4, 6, 12, and 24 MW/cm², and, assuming an average crystal size of 5 µm, to nominally 1, 2.4, 4.8, and 10 absorbed photons on average per chromophore (see details in the section *SFX data collection at SACLA* and Supplementary Table 4). For calculation of Fourier difference maps, *dark_ref_SACLA* and 66000 indexed light lattices were used per pump laser fluence (see justification in section *Dependency of integrated Fourier difference peaks as a function of indexed diffraction images at SACLA*). The resulting $F_o^{3\mu s\_fluence\_SACLA}$ - $F_o^{dark\_ref\_SACLA}$ maps (Supplementary Fig. 6) show light-induced features concentrated around the chromophore in each monomer that increase in strength with increasing pump-laser fluences. Positive (above $3\sigma$) and negative peaks (below $-3\sigma$) located within 2 Å from the AdoCbl chromophore were integrated around and plotted as a function of pump-laser fluence (Extended Data Fig. 4). Due to the limited number of data points (five, including (0, 0)) and the inherent error, the linear excitation regime cannot be determined precisely. It is reasonable to assume that excitation conditions at 30

$mJ/cm^2$ are not too far outside the linear excitation regime. To assure a certain safety margin, we decided to carry out the following TR-SFX time series at SACLA and at the SwissFEL at this fluence, corresponding to nominally 2.4 absorbed photons on average per chromophore (nominally 2.8 and 2 absorbed photons per chromophore in the front and the rear end of a 5 µm *Tt*CBD crystal, respectively; Supplementary Fig. 41).

In order to assess if the two key intermediate-state structures reported in the main text (10 ns and 3 µs, determined following pump-laser excitation with 30 $mJ/cm^2$) differ if excitation is carried out under single-photon excitation conditions, we collected an additional TR-SFX data set according to a *light-dark* sequence with a pump-probe delay of 10 ns at 12 $mJ/cm^2$ (corresponding to nominally 1 absorbed photon on average per chromophore). Intermediate-state structures refined against extrapolated structure factor amplitudes are almost identical at 12 and 30 $mJ/cm^2$, at both 10 ns and 3 µs (Supplementary Fig. 42). In particular, the Co–C5′ and Co–C4′ distances are identical within the coordinate error calculated by bootstrapping analysis (Supplementary Table 9). We conclude that photolysis of the Co–C5′ bond at 10 ns and formation of a putative Co–C4′ adduct at 3 µs are not the result of a multi-photon process.

#### Supplementary Note 3. Comparison of intermediate-state models in all monomers at the different TR-SFX time-delays

In the asymmetric unit of the crystals used in this study, a whole tetramer of *Tt*CBD is present. For clarity, monomer B has been the focus of structural interpretation in the main text. Here, we intend to describe the different intermediates modelled at all time delays in the four different monomers. The reference model is the structure of the *dark* state, which features a covalent bond between the cobalt atom and the C5′ (Co–C5′) of the adenosyl moiety. Careful inspection of the $mF_o$-$DF_c$ electron density maps of both *dark_ref_SACLA* and *dark_ref_SwissFEL* revealed residual positive peaks along some regions of the polypeptide chain of monomer D which triggered us to perform ensemble refinement with *PHENIX*[32,33]. From the 100 models present in the resulting ensemble from the *dark_ref_SACLA* data set, the root mean square fluctuation (RMSF) has been computed for Cα atoms of each monomer Supplementary Fig. 43). It clearly indicates increased RMSF values for the N-terminal part of monomer D (residues 80-155) with respect to the other monomers, corresponding to the four-helix bundle. Interestingly, monomer C also has large RMSF values, but for its C-terminal region (residues 200-275) which corresponds to the region of the Rossmann fold domain interfacing the four-helix bundle domain of monomer D. This clearly indicates an increased mobility of these structural domains of the tetramer in the crystalline environment. Also, the adenosyl moiety of AdoCbl, which is the leaving group upon photoactivation, is largely exposed to solvent in monomers A, B and D. In monomer C, a symmetry related molecule of the crystal greatly reduces the solvent accessibility of the adenosyl moiety, potentially affecting its capacity to leave the chromophore binding pocket. Supplementary Fig. 44 shows the electron density maps ($2mF_o$-$DF_c$ and $mF_o$-$DF_c$) for the four monomers at the active site, as well as a polder map ($mF_o$-$DF_c$_polder) with the adenosyl moiety omitted for *dark_ref_SACLA*. The maps indicate that the adenosyl moiety is well defined in all four monomers, with electron density present for the Co–C5′ covalent bond.

All intermediate-state structures described in this Supplementary Note refer to data collected with a pump fluence of 30 mJ/cm$^2$ (corresponding to nominally 2.4 absorbed photons per chromophore on average) for the different time-delays (10 ns, 300 ns, 3 μs, 100 μs, 3 ms at SACLA, 3 μs and 10 ms at SwissFEL). They have been refined against extrapolated structure factor amplitudes and modelled using extrapolated electron density maps ($2mF_{extr}$-$DF_c$ and $mF_{extr}$-$DF_c$).

At a time delay of 10 ns (referred to as *10ns_30mJ/cm²_SACLA*) following photoexcitation similar structural changes are observed in all four monomers. Strong negative peaks localized on the Co–C5′ bond of the dark model in the Fourier difference map $F_o^{10ns\_30mJ/cm2\_SACLA} - F_o^{dark\_ref\_SACLA}$ (-12.8 σ in monomer A, Extended Data Fig. 5 and Supplementary Fig. 8) are observed and strongly suggest that the Co–C5′ bond is broken. This observation is reinforced by the lack of electron density in the $2mF_{extr}$-$DF_c$ map between the cobalt atom of the cobalamin and the adenosyl group, as well as by the positive signal in a $mF_{extr}$-$DF_c$_polder map for the omitted adenosyl group which clearly indicates its separation from the cobalamin (Supplementary Fig. 12). The Co–C5′ bond lengths in the modelled dark state (*dark_ref_SACLA*) are 2.0 Å for all four monomers. In the 10 ns intermediate-state model, C5′ now points away from the cobalamin and is localized at 4.8 ± 0.1, 4.5 ± 0.1, 4.2 ± 0.1 and 4.1 ± 0.1 Å from the cobalt atom of monomers A, B, C and D, respectively. Such distances are not compatible with a covalent Co–C5′ bond. Other negative peaks lay on the

O3′ atom of the ribose ring and on the side chain of W131 (-7.0 σ for both). Two positive peaks can be seen next to the ribose ring (11.7 σ) and above residue W131 (6.7 σ). The displacement of the C5′ atom away from the corrin ring results in a reorientation of the W131 side chain, as well as a slight movement of the α-helix bearing this residue. The bond breakage causes a reorientation of the ribose ring via a rotation along the C1′–N9 bond linking the adenine as well as a conformational change from a 3′-endo in the dark model to a 2′-endo in the 10 ns intermediate-state model.

At the 300 ns time-delay (*300ns_30mJ/cm$^2$_SACLA*), the conformation of the adenosyl group did not evolve in monomer B with respect to that at 10 ns, with Co–C5′ distances of 4.4 ± 0.1 (compared to 4.5 ± 0.1 Å). For monomer A, this distance increases up to 5.3 ± 0.1 Å (4.8 ± 0.1Å at 10 ns) while it decreases in monomers C and D to 3.4 ± 0.1 and 3.7 ± 0.1 Å, respectively (compared to 4.2 ± 0.1 and 4.1 ± 0.1 at 10ns, respectively). The reorientation of the ribose ring brings the C4′ atom closer to the cobalt atom, with distances of 4.0 ± 0.1, 3.3 ± 0.1 and 2.9 ± 0.1 Å for monomers A, B and D, respectively. In monomer C, the adenosyl group is much closer to the corrin ring, with a distance between Co and C4′ atoms as short as 2.3 ± 0.1 Å. Intermediate-state models and 2mF$_{extr}$-DF$_c$ and mF$_{extr}$-DF$_c$_polder (adenosyl group omitted) electron density maps at 300 ns are shown in (Supplementary Fig. 13).

For the 3 µs data point measured at SACLA (*3µs_30mJ/cm$^2$_SACLA*), structural models refined against extrapolated structure factor amplitudes of all monomers (Supplementary Fig. 14) are similar to the model of monomer C at 300 ns (Supplementary Fig. 13). The Co–C4′ distance is now as short as 2.4 ± 0.1 Å for monomer A, 2.2 ± 0.1 Å for monomer B and 2.1 ± 0.1 Å for monomers C and D. The ribose ring of the adenosyl group still adopts the 2′-endo conformation observed at 10 ns. Intermediate-state models and 2mF$_{extr}$-DF$_c$ and mF$_{extr}$-DF$_c$_polder (adenosyl group omitted) electron density maps at 3 µs are shown in (Supplementary Fig. 14)

A 3 µs data point was measured at SwissFEL (*3µs_30mJ/cm$^2$_SwissFEL*) to serve as an overlap with the SACLA time series. The adenosyl group conformations of the SwissFEL and SACLA intermediate-state models refined against extrapolated structure factor amplitudes are very similar (Supplementary Fig. 15). First, the Co–C4′ distances are *ca.* 2.6 ± 0.1, 2.4 ± 0.1, 2.3 ± 0.1 and 2.4 ± 0.1 Å for monomers A, B, C and D, respectively in the SwissFEL intermediate-state model. The comparison was also performed by superimposing each monomer of the 3-µs SwissFEL intermediate-state model onto the respective monomer of the 3-µs SACLA intermediate-state model (using Cα atoms only). Then, the toot mean square deviation (RMSD) was computed using only the ribose atoms (i.e. C1' to C5' and O2' to O4') of the adenosyl group, yielding values of 0.40, 0.38, 0.18 and 0.40 Å for monomers A, B, C and D, respectively. Not only do these small values attest for very similar conformations of the ribose moiety of the adenosyl group, but they also indicate that their position within the chromophore binding pocket are nearly identical, as the atoms of the adenosyl group were not used in the superimposition calculation (Supplementary Fig. 15).

For the 100 µs time delay (*100µs_30mJ/cm$^2$_SACLA*), no evolution is observed in the refined intermediate-state model (Supplementary Fig. 16) compared to those at 3 µs (Supplementary Figs. 8 and 14). The Co–C4′ distances are 2.3 ± 0.1 Å and 2.4 ± 0.1 Å for monomers A and B, respectively, and 2.2 ± 0.1 Å for monomers C and D. Intermediate-state models and 2m$F_{extr}$-D$F_c$ and m$F_{extr}$-D$F_c$_polder (adenosyl group omitted) electron density maps at 100 µs are shown in (Supplementary Fig. 16).

For the next time-delay at 3 ms (*3ms_30mJ/cm$^2$_SACLA*), the modelled intermediate-state structures still indicate the presence of the adenosyl group close to the corrin ring with Co–C4′ distances of 2.5 ± 0.1 Å for monomers A, B and C, and 2.6 ± 0.1 Å for monomer D. If full occupancy of the adenosyl group is imposed, B-factors of that group increase by up to 43% with respect to the 100 µs model and negative peaks are observed in the m$F_{extr}$-D$F_c$ map on the adenosine moiety for monomers A and B (Supplementary Fig. 17). Consequently, either the occupancy of the adenosyl group is reduced at 3 ms compared to 100 µs or the disorder increased. Of note, the adenosyl group is still well ordered and present in monomer C. As stated above, the accessible solvent area of the chromophore-binding pocket is reduced in monomer C compared to the other monomers, because of the nearby presence of a symmetry related molecule of the crystal.

At a time-delay of 10 ms (*10ms_30mJ/cm²_SwissFEL*), large structural reorganization in the chromophore binding pockets of monomers A, B and D are observed. In these three monomers, large peaks of negative density are observed in the Fourier difference map $F_o^{10ms\_30mJ/cm2\_SwissFEL} - F_o^{dark\_ref\_SwissFEL}$ over the entire adenosyl group, suggesting that this group left the chromophore pocket in these monomers (Supplementary Fig. 10). This has been confirmed by the lack of any positive signal in the $mF_{extr}$-$DF_c$_polder map computed using the extrapolated structure factor amplitudes with the adenosyl group omitted (Supplementary Fig. 18). Furthermore, in monomers B and D, negative peaks over a large part of the corrin ring are observed in the Fourier difference map $F_o^{10ms\_30mJ/cm2\_SwissFEL} - F_o^{dark\_ref\_SwissFEL}$, with its maximum located on the cobalt atom (-12 $\sigma$ in monomer B, Supplementary Fig. 10). For these monomers, a positive peak is observed within a few Angstroms from the position of the cobalt atom in the dark-state structure (*dark_ref_SwissFEL*). Intermediate-state models refined against extrapolated structure factor amplitudes confirmed the movement of the corrin ring of about 4.3 Å in monomer B and 3.0 Å in monomer D (Supplementary Fig. 18) compared to the *dark_ref_SwissFEL* model. Importantly, reorientation of the H132 side chain is observed in monomers B and D, now pointing towards the Co atom, with H132NE2 - Co distances of 2.6 ± 0.2 and 4.7 ± 0.2 Å, respectively (11.0 ± 0.1 and 10.7 ± 0.1 Å in *dark_ref_SACLA* model). In monomer A, even though a negative peak is observed on the cobalt atom (-9 $\sigma$), no associated positive peak was present in the Fourier difference map, and only a minor shift of 0.5 Å was observed in the intermediate-state model when compared to the reference *dark_ref_SwissFEL* model. In monomer C, the adenosyl group is still present and has been modelled in a conformation similar to that observed at 3 ms, with a Co–C4′ distance of 2.7 ± 0.2 Å (from 2.5 Å at 3 ms).

A comparison of the 10 ms intermediate-state model and the cryo-trapped structure obtained after 5 s illumination at room temperature (PDB entry code: 8C76[2]) reveals very similar corrin-ring, H132 and H177 conformations in monomer B (Fig. 4c in main text).

#### **Supplementary Note 4. SSX on *Tt*CBD H132A mutant microcrystals at Diamond Light Source**

The H132A mutant of *Tt*CBD lacks the histidine residue that replaces the leaving adenosyl group and partakes in bis-His ligation of the corrin-ring Co atom in the light-adapted state of WT *Tt*CBD[45]. Upon illumination, corrin-ring displacement within the chromophore-binding pocket of the *Tt*CBD H132A mutant occurs as in the WT *Tt*CBD, yet without subsequent tetramer dissociation[2].

SSX experiments were conducted on H132A *Tt*CBD microcrystals with light-activation during 10 ms (*illuminated_DLS*) and without light activation (*dark_DLS*) at the I24 beamline at Diamond Light Source (see SSX data collection on *Tt*CBD H132A mutant microcrystals at Diamond Light Source for details). Because of the non-isomorphism observed between the structure factors of *dark_DLS* and *illuminated_DLS* data sets, the Fourier difference map was uninterpretable and therefore structure factor extrapolation was meaningless. However, as a large fraction of the crystalline molecules have been light-activated, computation of a $mF_o$-$DF_c$ Polder map with the adenosyl moiety omitted clearly indicates that the Co–C5′ bond has been broken and an intermediate species has been captured (Supplementary Fig. 36). This new adenosyl moiety conformation was first modelled using this $mF_o$-$DF_c$ polder map, and further refined against *illuminated_DLS* structure factors. The final model comprises two distinct conformations for the entire protein, representing i) crystalline molecules which have been light-activated (estimated occupancy of 0.80, 0.90, 0.95 and 0.80 for the monomers A, B, C and D, respectively) and ii) the crystalline molecules which remained in the dark state (this conformation was held fixed during the refinement). The intermediate state conformation of the final model refined at a resolution of 2.2 Å clearly indicates the structural rearrangement of the ribose ring of the adenosyl moiety: the Co–C5′ bond is broken, and the C4′ atom repositions closer to the Co atom of the cobalamin ring, with Co–C4′ distances of 2.5 Å for monomers A and C, and 2.8 Å for monomers B and D. For comparison, the Co–C4′ distances observed in the intermediate-state model of WT *Tt*CBD at 3 µs (*3µs_30mJ/cm$^2$_SACLA*) are 2.1 Å (monomers C and D), 2.2 Å (monomer B) and 2.4 Å (monomer A). The ribose rings of the adenosyl groups in models *illuminated_DLS* and *3µs_30mJ/cm$^2$_SACLA* are in a 2′-endo conformation. For each of the four monomers, atoms of the ribose rings (i.e. C1' to C5' and O2' to O4') have been used to first perform the superimposition and then compute the RMSD, yielding values of 0.27, 0.20, 0.12 and 0.18 Å for each of the ribose rings. Experimental evidence by SSX that a Co–C4′ adduct is present in the *Tt*CBD H132A mutant after photoexcitation during 10 ms (Supplementary Fig. 36) facilitated identification of a Co–C4′ adduct in TR-SFX of WT *Tt*CBD at 3 μs after photoexcitation (Fig. 3b).

That the Co–C4′ adduct is still present on the ~10 ms time scale in the *Tt*CBD H132A mutant crystal structure (*illuminated_DLS)* while on the same timescale it already decayed in WT *Tt*CBD (Fig. 4a) is in line with time-resolved UV-vis absorption spectroscopy that shows the spectroscopic intermediate associated with the Co–C4′ adduct decays with a time constant of 50 ms in the *Tt*CBD H132A mutant (Supplementary Fig. 45), i.e. ~5 times more slowly than in WT *Tt*CBD (Extended Data Fig. 2d and Supplementary Fig. 2c-d).

**Supplementary Note 5. Cryotrapping of intermediate states of *Tt*CBD reaction pathway in solution and *in crystallo***

In order to determine whether the spectroscopic intermediate states resolved at room temperature could be cryotrapped, we monitored the spectral features following light-excitation of *Tt*CBD in solution at various cryogenic temperatures (Supplementary Fig. 19). Firstly, samples were illuminated with green-light (530 nm) for 15 minutes at a range of successive temperatures (77-190 K; Supplementary Fig. 19a-b), allowing any photochemical steps to be identified. Difference spectra, using a non-illuminated sample as the blank (Supplementary Fig. 19b), show that the initial photochemistry results in the formation of an identical intermediate to that observed in the time-resolved measurements in ~500 ns (Fig. 3E, intermediate 2 in Extended Data Fig. 2g). This species is characterized by a decrease in the absorbance bands at 384 nm and 575 nm, and simultaneous increase in absorbance features at 334 nm, 473 nm and 507 nm. As this step can occur below 200 K, a temperature that is generally regarded to be the 'glass transition' temperature of proteins[72,73], it is unlikely that any large scale motions occur during this photochemical step and implies that the adenosyl group remains in the binding pocket at this stage. Any subsequent non-photochemical steps were identified by firstly illuminating samples with green light (190 K for 15 min) to trigger the initial photochemistry. These samples were then warmed to progressively higher temperatures in the dark for 15 mins before re-cooling to measure absorbance spectra at 77 K (Supplementary Fig. 19a). Difference spectra show a number of absorbance changes at 260 K, including an increase at 356 nm and a decrease at both 378 nm and 568 nm (Supplementary Fig. 19a). These spectral features are identical to those observed in the species that is formed with a time constant of ~7 ms in the time-resolved measurements, confirming that they represent the same intermediate (intermediate 3 in Extended Data Fig. 2g).

The final light-adapted state is only formed by warming to 298 K, and is characterized by a further increase and red-shift of the band at 355 nm to 358 nm, a decrease at 331 nm and a new absorbance feature at 412 nm (Supplementary Fig. 19c). As these latter non-photochemical steps can only occur above 200 K (Supplementary Figs. 19a and 19d), which is regarded to be the 'glass transition' of proteins, they are therefore likely to be linked to more significant structural changes in the protein, such as the shifting of the corrin ring and related movements that accompany photoactivation. Overall, we can identify 4 spectrally distinct species from the cryotrapping absorption spectroscopy measurements (Supplementary Fig. 19d) leading us to propose the sequential reaction pathway shown in Supplementary Fig. 19e for the photoactivation of *Tt*CBD. Intermediate states 2 and 3 in the scheme correspond to the species iii and iv in Fig. 1 of main text and in Extended Data Fig. 2g.

It is currently unclear how the redox state of the Co atom in AdoCbl changes during the photocycle of CarH although EPR spectroscopy has been used to identify the Co(II) state of AdoCbl in anaerobically photolyzed CarH[2]. Hence, we have now coupled our cryotrapping approach with EPR measurements at 20 K to characterize the redox state of the Co upon formation of intermediate 2 (species iii in Fig. 1 and in Extended Data Fig. 2g). As previously reported[2], dark state and final aerobic light state of *Tt*CBD do not give rise to any EPR signals, demonstrating the Co(III) state of the AdoCbl in these species. Similar to previous measurements[2], photolysis under anaerobic conditions produced a paramagnetic Co(II) signal with a low-spin Co(II) species observed in the EPR spectra (Fig. 3F). To investigate the redox state of intermediate 2 (species iii in Fig. 1) we initially illuminated samples at 180 K for varying lengths of time to trap this intermediate in the pathway (Fig. 3F and Supplementary Fig. 22a). The intermediate is EPR silent, confirming that no Co(II) species are present at this stage of the reaction. Moreover, no EPR signatures are observed when EPR spectra are recorded at higher temperatures (up to 220 K), suggesting that a coupled Co(II)-radical species is unlikely (Supplementary Fig. 22b). Therefore, based on the observed UV/vis absorbance changes, which show the absence of a typical Co(I) cobalamin absorbance peak at 388 nm (Supplementary Fig. 19d), we suggest that the AdoCbl in intermediate 2 (species iii in Fig. 1) is in the Co(III) state. Similarly, the UV-vis absorbance features (Supplementary Fig. 19d) of the subsequent intermediate 3 (species iv in Fig. 1), in particular the absorbance band at 355 nm, is identical to those observed for an H132A mutant (Supplementary Fig. 24), which has been previously shown to form a Co(III) light state species that is ligated with water/OH as the 6th ligand[2]. Hence, this suggests that species iv in Fig. 1 is also likely to represent a water/OH-ligated Co(III)-H132 species, although it should be noted that Co(I)/Co(II) species may be formed when $O_2$ concentration is limited[2].

The cryogenic spectroscopic measurements performed in solution have guided the cryo-temperature dependent macromolecular crystallography experiments with the aim to cryotrap intermediate state(s) (see *Cryo-temperature dependent macromolecular crystallography and in crystallo UV-vis spectroscopy* section). *In crystallo* UV-vis absorption microspectrophotometry on the FIP-2 (ESRF) beamline was performed to monitor spectroscopic changes after illumination of *Tt*CBD macrocrystals at different temperatures ranging from 100 K up to 210 K. The intermediate state trapped after illumination at 180 K and characterized by cryo MX (referred to as *cryo_light_180K*) presents structural similarities in the chromophore binding pocket with the intermediate-state models obtained at room temperature by TR-SFX with a pump-probe delay of 3 µs (*3µs_12mJ/cm²_SACLA, 3µs_30mJ/cm²_SACLA and 3µs_30mJ/cm²_SwissFEL*). The structural comparison described below has been performed with the *3µs_30mJ/cm²_SACLA* intermediate-state model (for an overlay see Fig. 3c in main text). In all four monomers of the *cryo_light_180K* model, the Co–C5′ bond linking the adenosyl moiety to the corrin ring has been cleaved, as shown by lack of continuous $2mF_o$-$DF_c$ electron density between the two atoms (Supplementary Fig. 21) and Co–C5′ distances of 3.5, 3.2, 3.3 and 3.5 Å for monomer A, B, C and D, respectively and in good agreement with the Co–C5′ distances of 3.6 ± 0.1, 3.2 ± 0.1, 3.0 ± 0.1 and 3.0 ± 0.1 Å observed in the *3µs_30mJ/cm²_SACLA* intermediate-state structure. The ribose conformations of the adenosyl group are almost identical in both models, with an RMSD of 0.27, 0.18, 0.13 and 0.21 Å for monomers A, B, C and D, respectively (after superimposition of the ribose atoms). Furthermore, the C4′ atom of the ribose approaches the Co atom at distances of 2.7, 2.5, 2.6 and 2.6 Å for *cryo_light_180K* structure and of 2.4 ± 0.1, 2.2 ± 0.1, 2.1 ± 0.1 and 2.1 ± 0.1 Å for *3µs_30mJ/cm²_SACLA* structure for monomers A, B, C and D, respectively. In the *3µs_30mJ/cm²_SACLA* intermediate-state model, all Co–C4′ distances are in excellent agreement with the C4′ adduct species proposed by QM/MM (2.1 Å). The Co–C4′ distances in the *cryo_light_180K* structure are slightly longer. For monomers B, C and D, these distances are still indicative of a C4′ adduct being formed at 180 K, rather than any other chemical species determined by QM/MM that would have a Co–C4′ distance of at least 3.3 Å (Supplementary Table 2). The discrepancy in the Co–C4′ distances between the *cryo_light_180K* and *3μs_30mJ/cm²_SACLA* structures most probably arises from the different positioning of the corrin ring, and consequently of the Co atom. The position of the corrin ring in the *cryo_light_180K* intermediate-state is identical to that of the dark state (*dark_ref_SACLA*), while the corrin ring is shifted in the *3μs_30mJ/cm²_SACLA* one (Fig. 3c, ranging from 0.4 Å for monomer C to 0.7 Å for monomer B). This discrepancy most probably originates from the different temperatures at which the reaction has been triggered (180 K vs room-temperature), but the structural similarities between these models strongly suggest that the same intermediate-state species is present at 180 K and at 3 μs.

**Supplementary Note 6. Cluster-model and QM/MM calculations**

Comparing the conversion of the hydridocobalamin species (HCbl) + anhydroadenosine (anhAdo) photoproduct (structure **1** in Supplementary Fig. 23b) to the Co–C4′ and Co–C5′ adducts in our cluster model help explain why the Co–C4′ adduct is formed and what role it might play in the overall mechanism. The potential energy profile and structures are shown in Supplementary Fig. 23 and structural parameters in Supplementary Table 3. The barrier for hydrogen atom transfer from HCbl to C5′ in anhAdo (structure **1** in Supplementary Fig. 23) is much lower than the barrier for hydrogen atom transfer to C4′ (2.08 *cf.* 14.2 $kcal.mol^{-1}$). After transfer to C5′, an intermediate Co···H···C5′ structure (**3A**) is formed, which seems to correspond formally to a C5′ cation and therefore Co(I) based on the atomic charges (Supplementary Table 10) and the zero spin density on all atoms. Transfer to C4′ on the other hand leads to a Co(II) + C5′· diradical (**3B**), with the transferring hydrogen oriented towards the Co, and a rotation around the C4′ then repositions the adenosyl radical (**4B**) to allow bond formation between the Co(II) and C5′· to form the Co–C5′ adduct (**5B**). By contrast, we have not observed a stable Co(II) + C4′·diradical structure, but rather a [Co(II) + C4′·]$^‡$ transition state (**4A**) which corresponds to breaking the Co–H bond and rotation of the C4′ towards the Co, which then leads to the formation of the Co–C4′ adduct (**5A**). Note that the [Co(II) + C4′·]$^‡$ transition state (**4A**) is lower in energy than the corresponding diradical along the pathway towards the Co–C5′ adduct, Co(II) + C5′· (**4B**), owing to the greater stability of the tertiary C4′· radical compared to the primary C5′· radical.

For both the Co–C4′ and Co–C5′ adducts two structures with different Co–C and Co–$N_{H177}$ bond distances (Supplementary Table 3) were identified after a relaxed Co–N distance scan. In the profile we included the structures that are in closest agreement with the crystal structures (*dark_DLS* for the Co–C5′ adduct and *illuminated_DLS* for the Co–C4′ adduct) in terms of these two bonds. For the Co–C5′ adduct the selected structure has the shorter Co–$N_{H177}$ bond compared to the alternative Co–C5′ adduct structure (2.09 Å *vs* 2.51 Å), a slightly shorter although very similar Co–C5′ bond (2.01 Å *vs* 2.05 Å), and is energetically favoured by -3.26 kcal $mol^{-1}$. For the Co–C4′ adduct the selected structure has a slightly higher energy of 1.07 kcal $mol^{-1}$ than the alternative Co–C4′ adduct, although this small energy difference would not be sufficient to confidently discriminate between the two structures. The selected Co–C4′ adduct structure has a longer Co–$N_{H177}$ bond (2.44 Å *vs* 2.14 Å), and a longer Co–C4′ bond (2.17 Å *vs* 2.09 Å) than the alternative structure.

Overall, the largest potential energy barrier for the formation of the Co–C5′ adduct is 14.2 kcal $mol^{-1}$, while for formation of Co–C4′ it is 4.67 kcal $mol^{-1}$ (forming the $[Co(II) + C4'\cdot]^{\ddagger}$ species). Therefore, once the photoproduct (**1**) has been formed after photoexcitation, the formation of Co–C4′ adduct can occur before the release of the photoproduct (occurring on the ms time scale, cf. Fig. 4a). From the Co–C4′ adduct, the photoproduct can be reformed after homolytic cleavage of the Co–C4′ bond, with a potential energy barrier of 16.6 kcal $mol^{-1}$ (**5A → 4A**), which is significantly lower than the computed 34.7 kcal $mol^{-1}$ barrier for Co–C5′ bond cleavage (**5B → 4B**). Note that our computed potential energy barrier for Co–C5′ bond cleavage is somewhat larger than that computed previously using QM/MM calculations[74], which suggests that our model is not fully capturing the catalytic effect of the protein environment. Nevertheless, any such effect is likely to be similar for both adducts, and these calculations reveal inherent differences between the adducts and their chemistry within the protein binding pocket. Crucially, our results suggest that once the Co–C4′ adduct has been formed, a return to the photoproduct is thermally accessible (especially if our computed barriers are overestimated). This suggests that the Co–C4′ adduct might serve to “protect” the system from returning to the dark Co–C5′ state before the photoproduct can leave the binding site. The proposed CarH photoactivation mechanism based on the DFT cluster model calculations is shown in Extended Data Fig. 9.

To explore the chemical nature of the crystallographically determined intermediate-state structures, we performed QM/MM calculations. Selected geometric parameters for AdoCbl of the dark state form of the crystal structure (*dark_ref_SACLA*) and of the corresponding optimized QM/MM model are compared in Supplementary Table 11. For a superposition of both structures see Supplementary Fig. 46a. The Co–C5′ distance remains consistent between all four monomers with the QM/MM calculated distance. The other important distinction to highlight is that each monomer from the crystal structure indicates a Co–$N_{H177}$ bond distance that is shorter than the optimized QM/MM distance of 2.33 Å. It has been demonstrated based on several crystal structures of various $B_{12}$ enzymes that the elongation of the Co–R bond either shortens (normal trans influence) or lengthens (inverse trans influence) the Co–Nax bond[75-79]. The Co–$N_{H177}$ bond is typically elongated in cobalamins due to the inverse trans effect[80]. The conformation of the ribose portion of the adenosyl moiety was determined and is collected in Supplementary Table 11. For each monomer of the crystal structure and for the QM/MM model, the ribose is found in the 3′-endo conformation.

In order to interpret the experimental 10 ns structure (*10ns_30mJ/cm²_SACLA*) in terms of the electronic structure two QM/MM models were considered. Namely one which assumed a diradical species Co(II)/C5′• (referred to as the 10ns-diradical-model hereafter) and another which assumed the formation of the photoproducts (HCbl and anhAdo) which is referred to as the 10ns-photoproduct-model hereafter. In order to achieve the latter model, the hydrogen from the C4′ of the ribose ring of the adenosyl ligand was transferred to the cobalt in the initial setup of the model. The 10ns-diradical-model required the use of an unrestricted wavefunction which allowed for the spin density for the atoms in the QM region to be determined. The optimized 10-ns-diradical-model involved a Co(II)/C5′• radical pair with spin densities of 0.93/-1.0, respectively. For the 10ns-photoproduct-model, a restricted wavefunction was used. A comparison of the geometric parameters from the crystal structure (*10ns_30mJ/cm²_SACLA*) and the QM/MM results for both the 10ns-diradical-model and the 10ns-photoproduct-model are listed in Supplementary Table 1. The largest difference in distances for the two QM/MM models are in Co–C4′ (difference of 1.4 Å). Based on the comparison of the Co–C4′ bond distances of the 10ns-photoproduct-model and the 10-ns-diradical model with the one in the 10 ns structure (*10ns_30mJ/cm²_SACLA*), it appears that the AdoCbl in the experimental 10 ns structure is best described as a Co(II)/C5′• radical pair. The optimized geometry of AdoCbl is shown in Supplementary Fig. 46b superimposed on the 10 ns crystal structure (*10ns_30mJ/cm²_SACLA*). It is apparent that the geometry of the optimized diradical is qualitatively in-line with the 10 ns crystal structure (*10ns_30mJ/cm²_SACLA*) especially in terms of the orientation of the ribose and adenine portions of the adenosyl ligand.

The possibility that the 10 ns structure (*10ns_30mJ/cm²_SACLA)* is best described as a diradical was further explored using the geometry of the experimental 3 μs structure (*3μs_30mJ/cm²_SACLA*) as the initial guess for the QM/MM model. Optimizing a diradical species from a different geometry than the 10 ns crystal structure would be an independent verification of the electronic structure of the 10 ns crystal structure. To set up this test using the 3 μs structure (*3μs_30mJ/cm²_SACLA*), an unrestricted wavefunction was used, the C5′ was doubly protonated and the Co–C5′ distance was elongated prior to the optimization. The spin density on the cobalt was 0.91 and for the C5′ the spin density was -0.96 which confirms that a diradical species was optimized.

The AdoCbl diradical species was previously investigated using QM/MM methods from the CarH crystal structure (PDB entry code: 5C8E[45]) where a potential energy surface (PES) based on the Co–C5′ and Co–C4′ distances was created[81]. This PES connected the dark state, the diradical, and the hydridocobalamin and 4′,5′-anhydroadenosine photoproducts (Supplementary Fig. 47). A major conclusion from this study was that the plateau-like region on the PES corresponded to the diradical species. This region spanned from Co–C5′ distances of ~3.2-4.2 Å and Co–C4′ distances of 3.7-4.1 Å. It was found that any of the structures in this region could be optimized to a stable diradical. The photoproduct was formed with 10ns_30mJ/cm²_SACLA distances longer than 5.0 Å and Co–$C_{4'}$ distances greater than 4.4 Å. We compared the Co–C5′ and Co–C4′ distances of the 10-ns-diradical model and the 10ns-photoproduct-model to this PES (Supplementary Fig. 47). The 10-ns-diradical model has Co–C5′ and Co–C4′ distances of 3.89 Å and 3.08 Å, respectively, while the 10-ns-photoproduct model has Co–C5′ and Co–C4′ distances of 4.10 Å and 4.48 Å, respectively. Based on the previous QM/MM study[81], these distances are in line with those in the diradical region.

The experimental 3 μs structure (*3μs_30mJ/cm²_SACLA*) was optimized considering four different assumptions (Extended Data Fig. 7). The first model (Extended Data Fig. 7a) optimized the photoproducts, 4′,5′-anhydroadenosine and hydridocobalamin. The initial set up of this model necessitated that the C4′ hydrogen be transferred to the cobalt ion. After optimization, the Co–C5′

distance was 4.18 Å while the Co–C4′ distance was 4.24 Å. The second model (Extended Data Fig. 7b) we tested was the presence of a singlet diradical species which included the C5′ primary radical and the Co(II) ion. The spin density was -0.96 for the C5′ and 0.91 for the Co(II) and the optimized Co–C5′ and Co–C4′ distances were 4.01 and 3.25 Å, respectively. The third model (Extended Data Fig. 7c) we tested included a C4′ tertiary radical with a Co(II) ion. For this optimized singlet diradical the Co–C5′ and Co–C4′ distances were 3.92 and 4.10 Å, respectively. Here the spin density was 0.99 on the cobalt, -0.75 on the C4′ tertiary radical and -0.12 on the oxygen of the ribose ring (O4′ in Extended Data Fig. 1). The fourth model (Extended Data Fig. 7d) that we tested was that an adduct was formed which involved a bond between the cobalt ion and the C4′. The Co–C4′ adduct species involved a sp3 C5′. The optimized distances for the Co–C5′ and Co–C4′ were 2.92 Å and 2.12 Å for the adduct species. It should also be pointed out that the Co–C4′ optimized adduct exhibited an elongated Co–$N_{H177}$ distance of 2.92 Å. We were unable to locate a stable minimum where the Co–C4′ adduct had a shorter Co–$N_{H177}$ distance. The key geometric parameters for the cobalamin for each of the four QM/MM optimized models are listed in Supplementary Table 2 which also includes the dihedral angles associated with the ribose conformation. The 3 µs crystal structure indicates a ribose conformation of 2′-endo. The photoproducts QM/MM model (HCbl + anhAdo), the primary diradical QM/MM model (Co(II)/C5′• RP) and the tertiary diradical QM/MM model (Co(II)/C4′• RP), and finally the adduct QM/MM model all indicate a ribose conformation of 2′-endo. Based on the analysis of the Co–C5′ and Co–C4′ distances, it appears that the 3 µs crystal structure should be classified as a stable adduct with a Co–C4′ bond. The 3 µs crystal structure (*3µs_30mJ/cm²_SACLA*) and the optimized Co–C4′ adduct QM/MM model are superimposed in Supplementary Fig. 46c.

### Supplementary figures

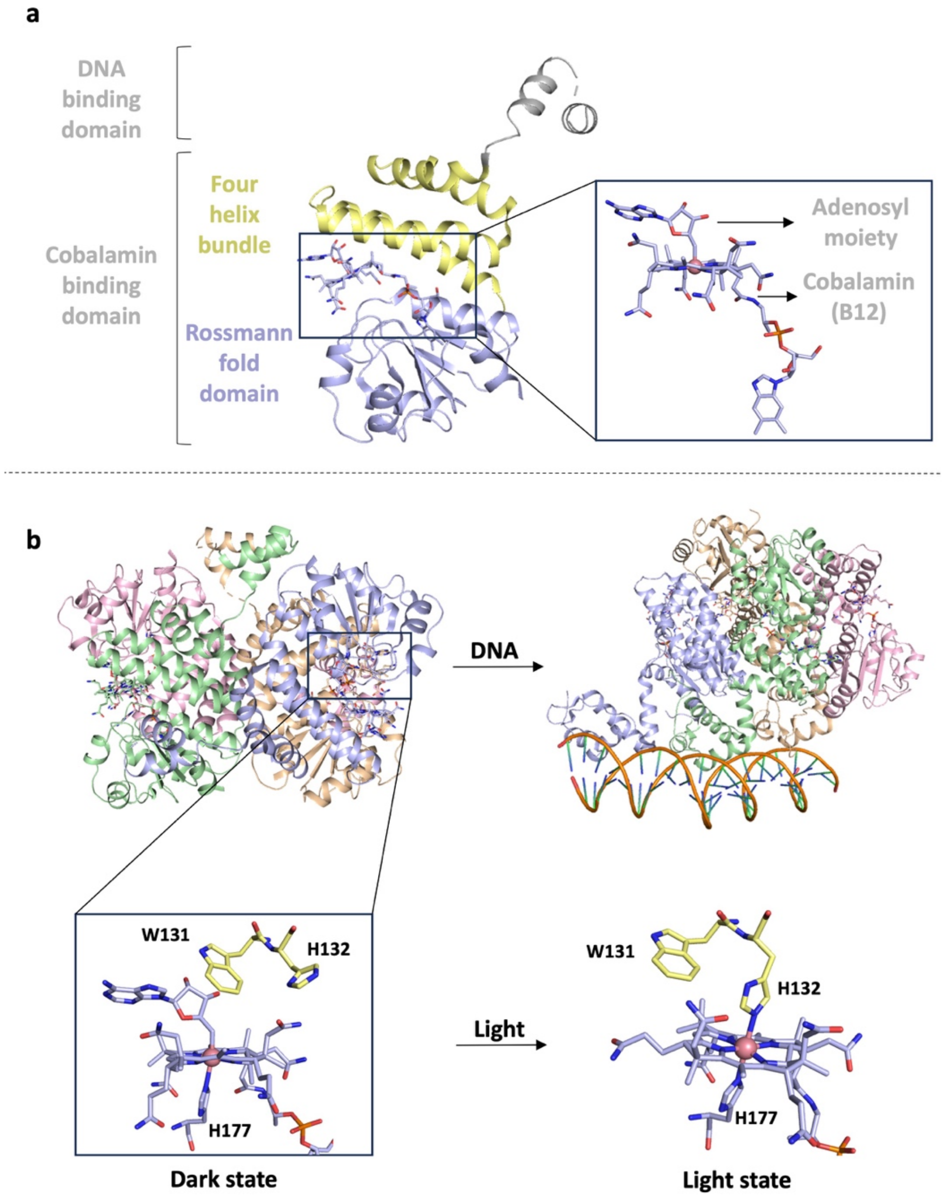


**<u>Supplementary Fig. 1. Crystal structures of full-length CarH before and after illumination.</u>**

(**a**) The monomer of full length *Tt*CarH is composed of two domains: a DNA binding domain (DBD) and the cobalamin binding domain (CBD). The CBD consists of two motifs, a four-helix bundle and a Rossmann fold domain. (**b**) Structural changes after photoactivation of CarH. On the upper row, models of full length *Tt*CarH in its dark state (tetrameric form) are shown in cartoon in absence of (left, PDB code: 5C8D) and complexed (right, PDB code: 5C8E) to DNA. The inset (lower left) represents the AdoCbl in the dark state as well as key residues (H132, W131 and H177) in sticks. In the dark state, the cobalt atom is covalently bonded to the adenosyl moiety (Co–C5′) and is also coordinated to H177. Upon photoactivation, the Co–C5′ bond is broken. Thus, the adenosyl group leaves the chromophore binding pocket and structural reorganizations yield to tetramer dissociation and formation of a bis-histidine adduct referred to as light state (lower right, PDB code: 5C8F (*5*).

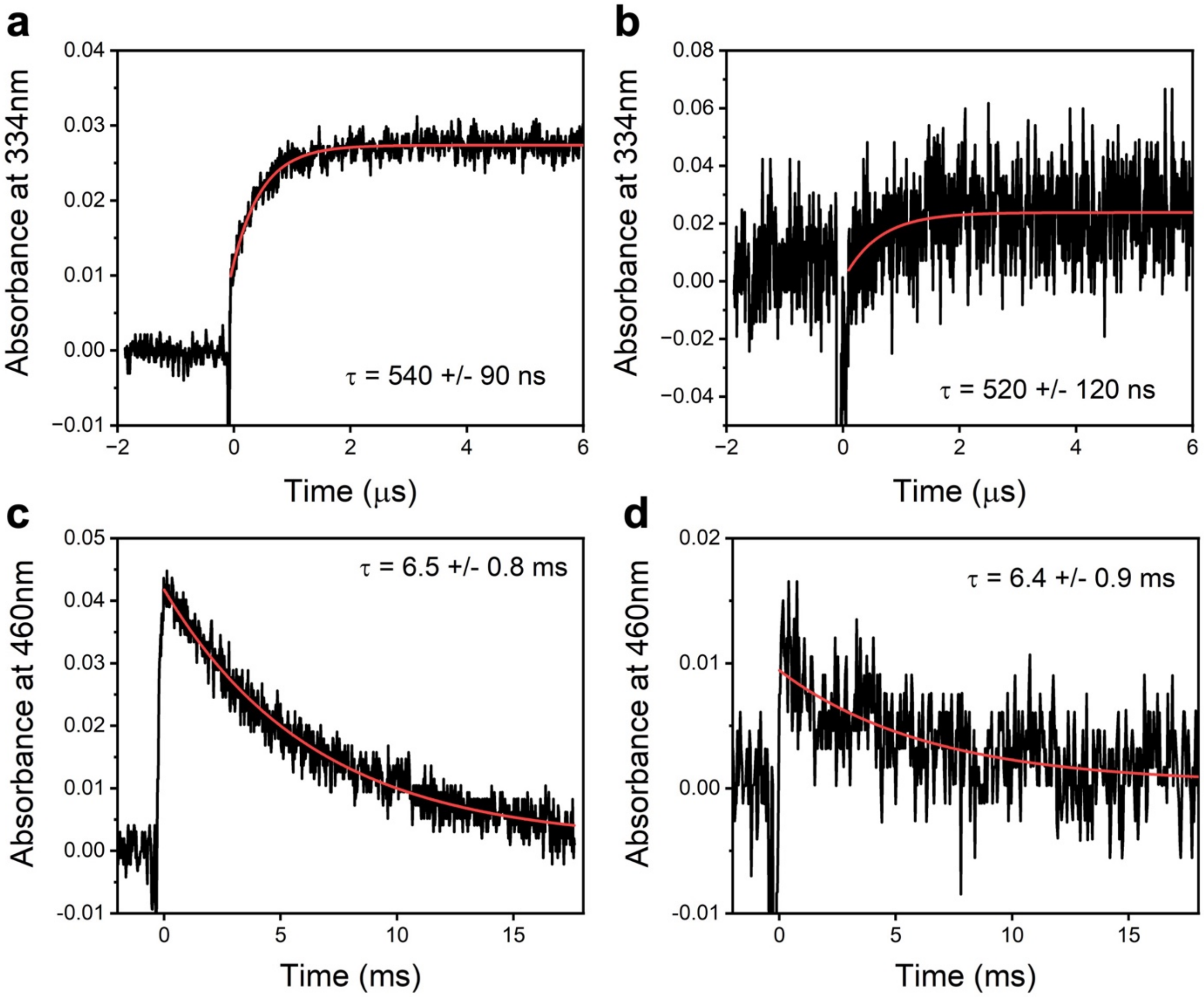


**Supplementary Fig. 2. Comparison of time-resolved absorption spectroscopy changes between *Tt*CBD in solution and *Tt*CBD microcrystals embedded in cellulose.**

Kinetic transients at 334 nm on the μs timescale of samples containing 500 μM *Tt*CBD in solution (**a**) or 20 % (v/v) microcrystal slurry in CMC-4% (**b**). Kinetic transients at 460 nm on the ms timescale of samples containing 200 μM *Tt*CBD in solution (**c**) or 20 % (v/v) microcrystal slurry in CMC-4% (**d**). Data were fitted to a single exponential equation to obtain time constants (red lines). All transients were measured at room temperature and data shown are the average of at least five traces.

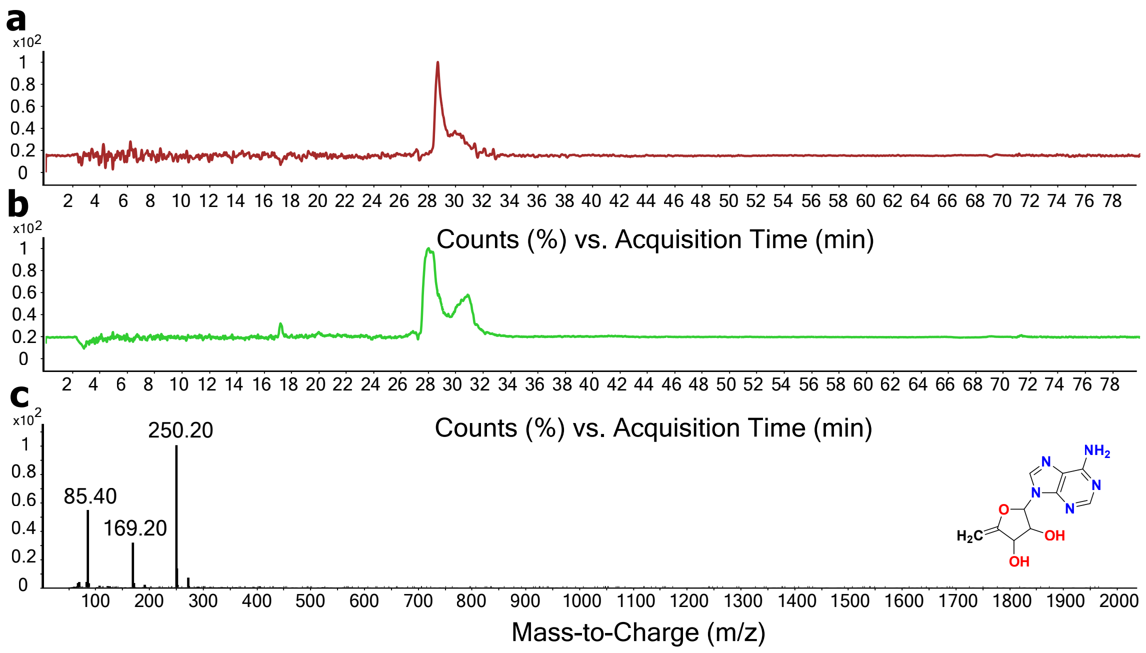


**Supplementary Fig. 3. LCMS of the photoproduct formed when illuminating *Tt*CBD crystals**

Microcrystals illuminated with ambient light were analysed by LCMS. Panels (**a**) and (**b**) show the HPLC traces measured at 254 nm of crystals never exposed to light (red) and crystals illuminated with ambient light. Due to a slight drifting of the baseline, likely due to the presence of PEG, a control acetonitrile spectrum was subtracted from the final spectrum. The dark spectrum shown in red is devoid of any peaks of interest. The photoproduct is observed in the green spectrum as a small peak at around 17 minutes. The masses observed at this retention time are shown in panel (**c**). A mass-to-charge of 250 corresponds to the photoproduct observed earlier[4,5].

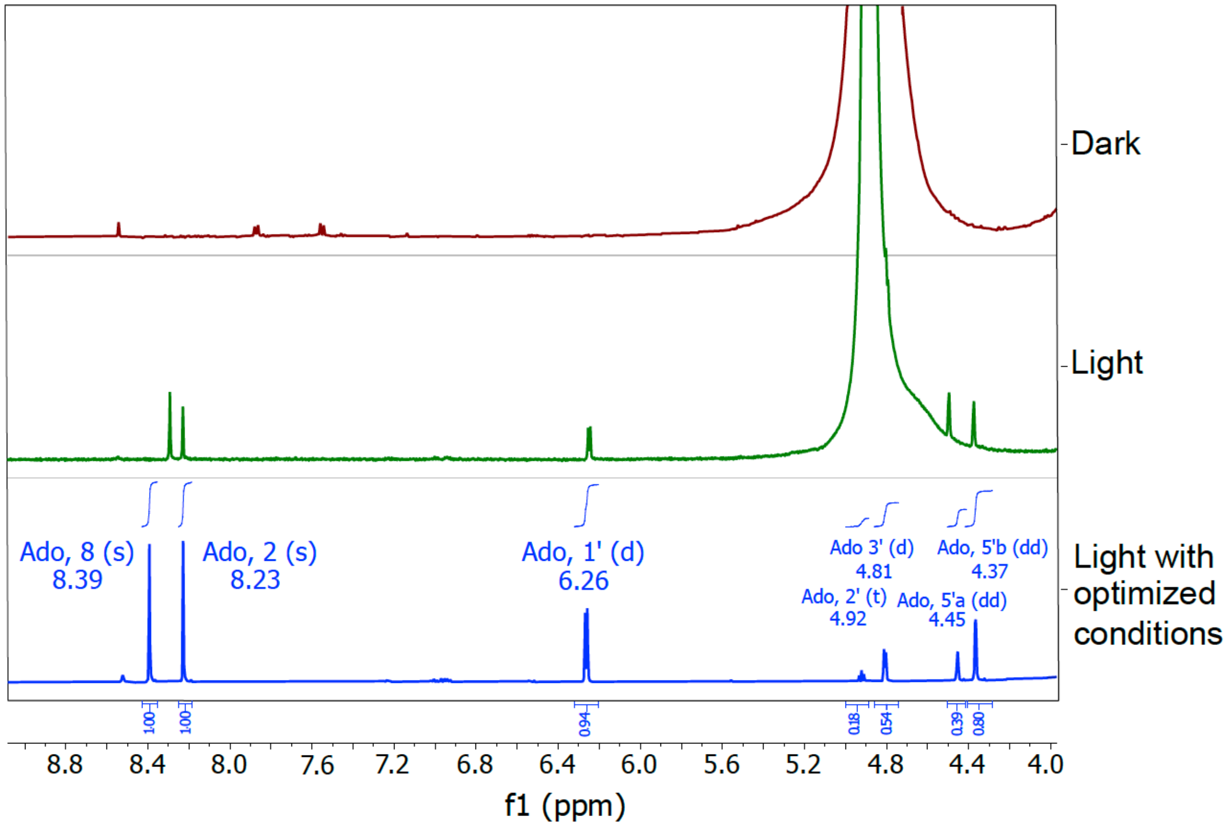


**<u>Supplementary Fig. 4. Determination by NMR of photoproduct formed when illuminating *Tt*CBD microcrystals</u>**

Microcrystals illuminated with ambient light were analysed by NMR to detect the presence of the photoproduct (*5*)[5]. The dark spectrum shown in red is devoid of any peaks of interest. The photoproduct observed in the green spectrum shows several key peaks for identification. This compound is comparable to signals already reported earlier[4,5]. However, two characteristic peaks of 4′,5′-anhydroadenosine were missing due to large solvent peaks observed around 4.8 ppm. Optimization of the parameters led to the generation of the blue spectrum with all characteristic peaks of the photoproduct. Associated values are shown in blue boxes. The red and green spectra are multiplied by a factor of two for comparison with the blue spectrum.

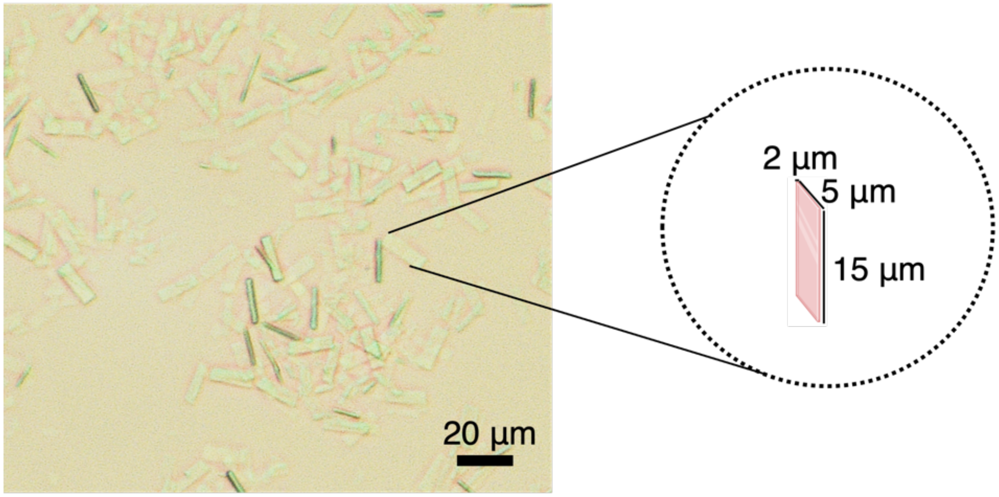


**Supplementary Fig. 5. *Tt*CBD microcrystals**

*Tt*CBD microcrystals in the storage buffer (10% PEG 10000, 0.05 M HEPES pH 7.5) were imaged with a Hirox HRX-01 optical microscope. Microcrystals were homogeneous in size with a plate-like shape (shown in the dashed circle), with respective dimensions of ~15 × 5 × 2 μm.

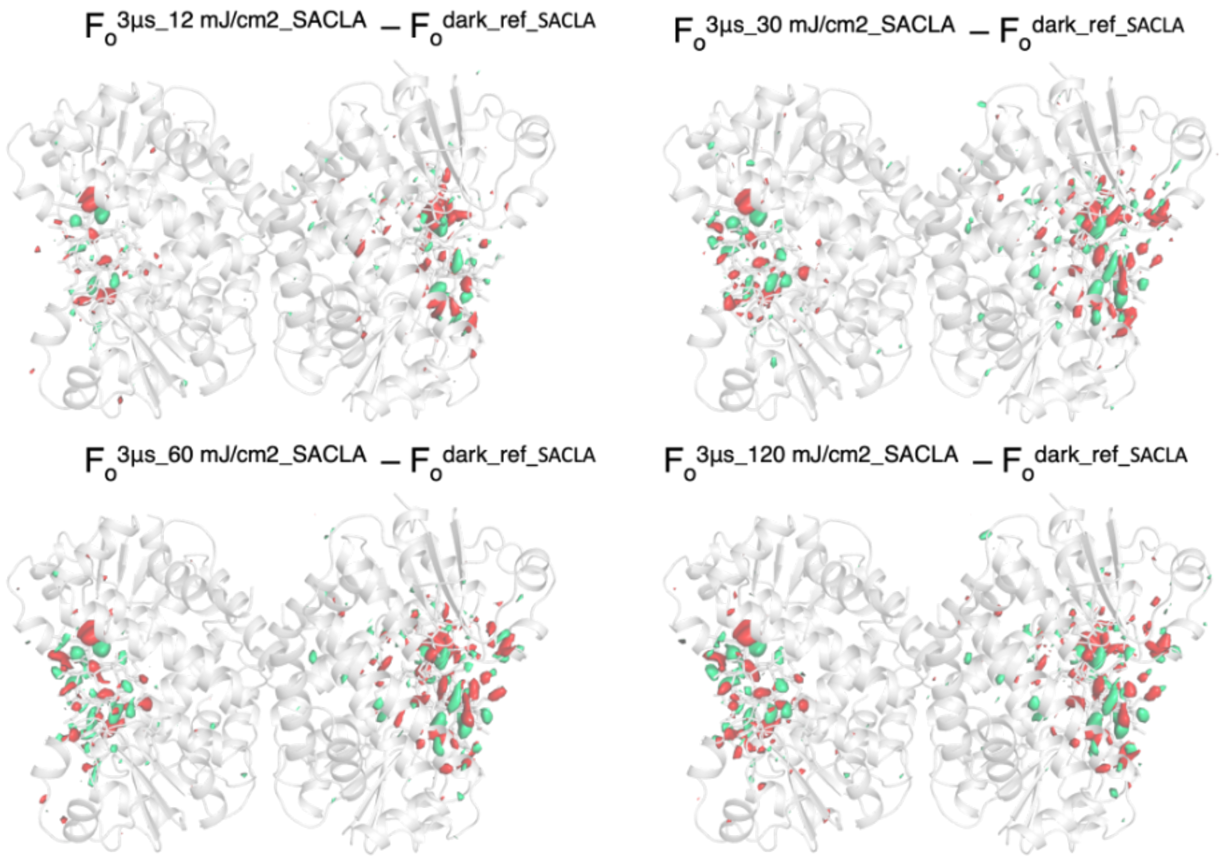


**Supplementary Fig. 6. Fourier difference maps computed with the various data sets from the power titration performed at SACLA (3 µs time delay).**

Model of *dark_ref_SACLA Tt*CBD tetramer is represented as a grey cartoon. Fourier difference maps $F_o^{3µs\_fluence\_SACLA} - F_o^{dark\_ref\_SACLA}$ (fluence: 12, 30, 60 and 120 $mJ/cm^2$) are contoured at +3.5 σ (green) and -3.5 σ (red).

$F_o^{dark} - F_o^{dark\_only\_SACLA}$
10 ns, 12 mJ/cm$^2$

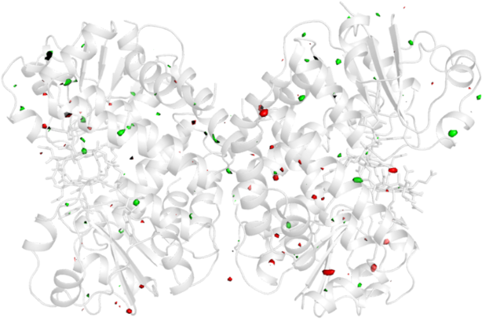

$F_o^{dark} - F_o^{dark\_only\_SACLA}$
10 ns, 30 mJ/cm$^2$

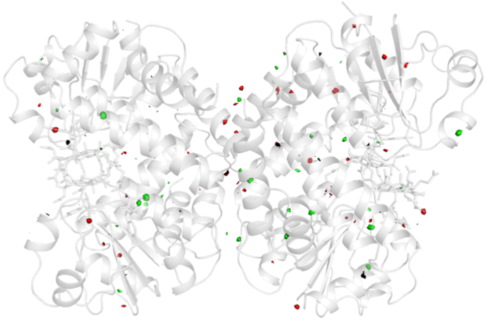

$F_o^{dark} - F_o^{dark\_only\_SACLA}$
300 ns, 30 mJ/cm$^2$

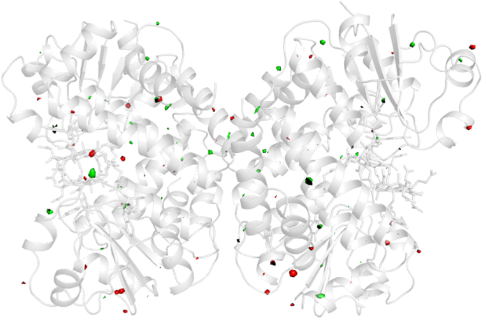

$F_o^{dark\_1} - F_o^{dark\_only\_SACLA}$
3 μs, 12 mJ/cm$^2$

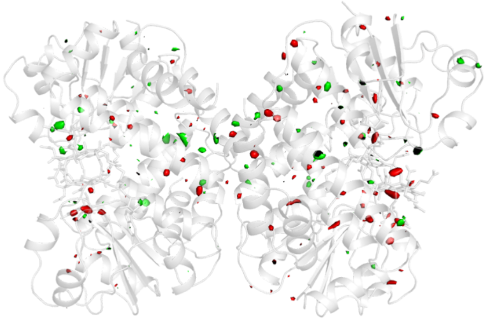

$F_o^{dark\_2} - F_o^{dark\_only\_SACLA}$
3 μs, 12 mJ/cm$^2$

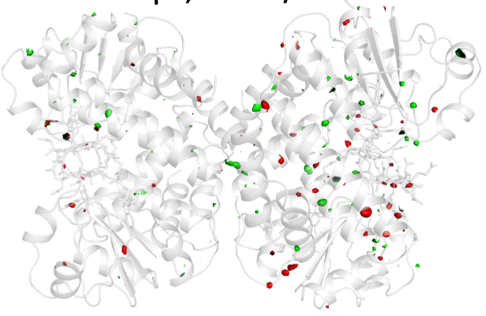

$F_o^{dark\_1} - F_o^{dark\_only\_SACLA}$
3 μs, 30 mJ/cm$^2$

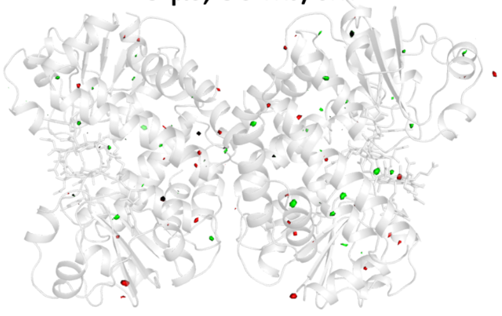

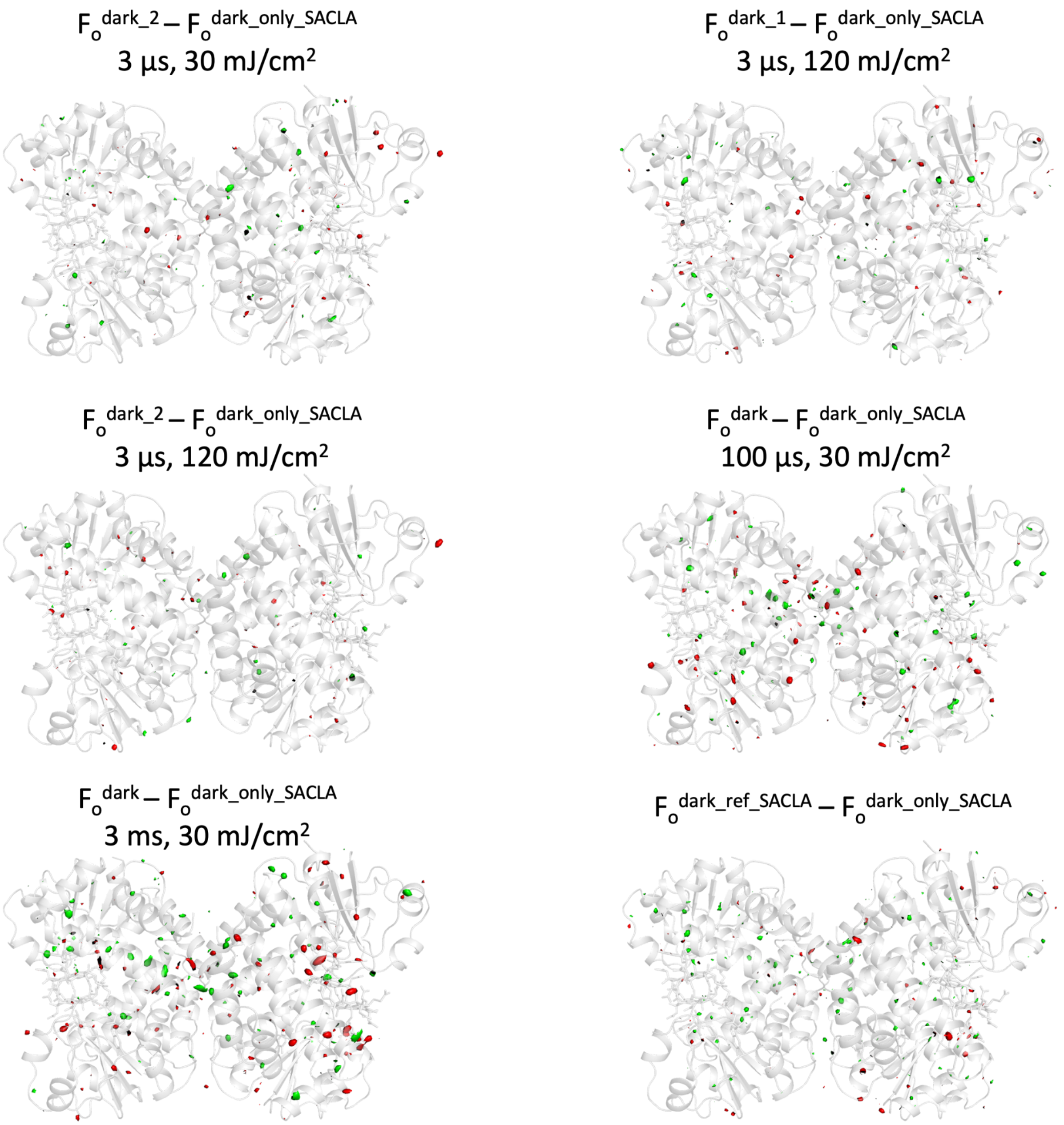


**Supplementary Fig. 7. Fourier difference maps of interleaved dark data sets at different time delays and/or laser fluences compared to *dark_only_SACLA*.**

Model of *dark_ref_SACLA TtCBD* tetramer is represented as a grey cartoon. Fourier difference maps $F_o^{dark} – F_o^{dark\_only\_SACLA}$ are contoured at +3.5 $\sigma$ (green) and -3.5 $\sigma$ (red). *Dark* data sets were collected in a *light-dark* sequence, *dark1* and *dark2* in a *light-dark1-dark2* sequence.

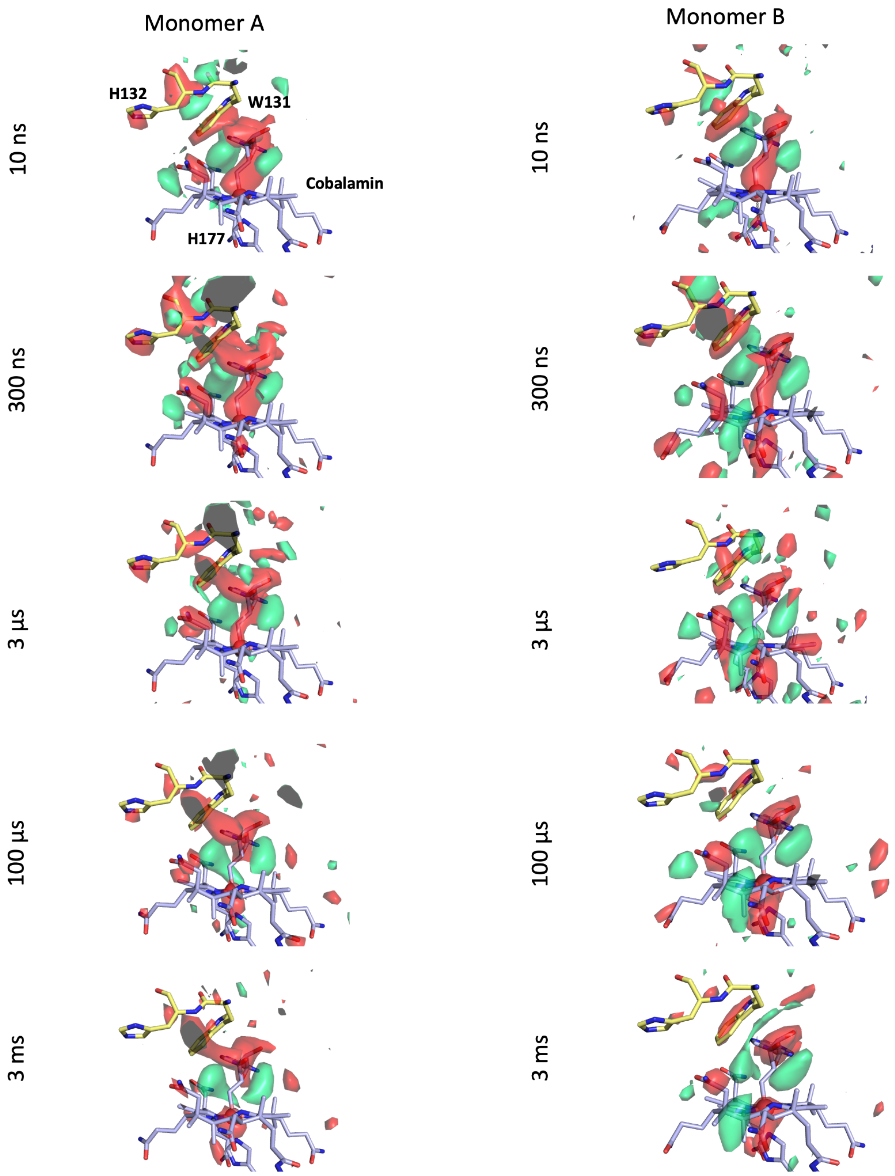

Monomer A
Monomer B
10 ns
300 ns
3 µs
100 µs
3 ms
H132
W131
Cobalamin
H177

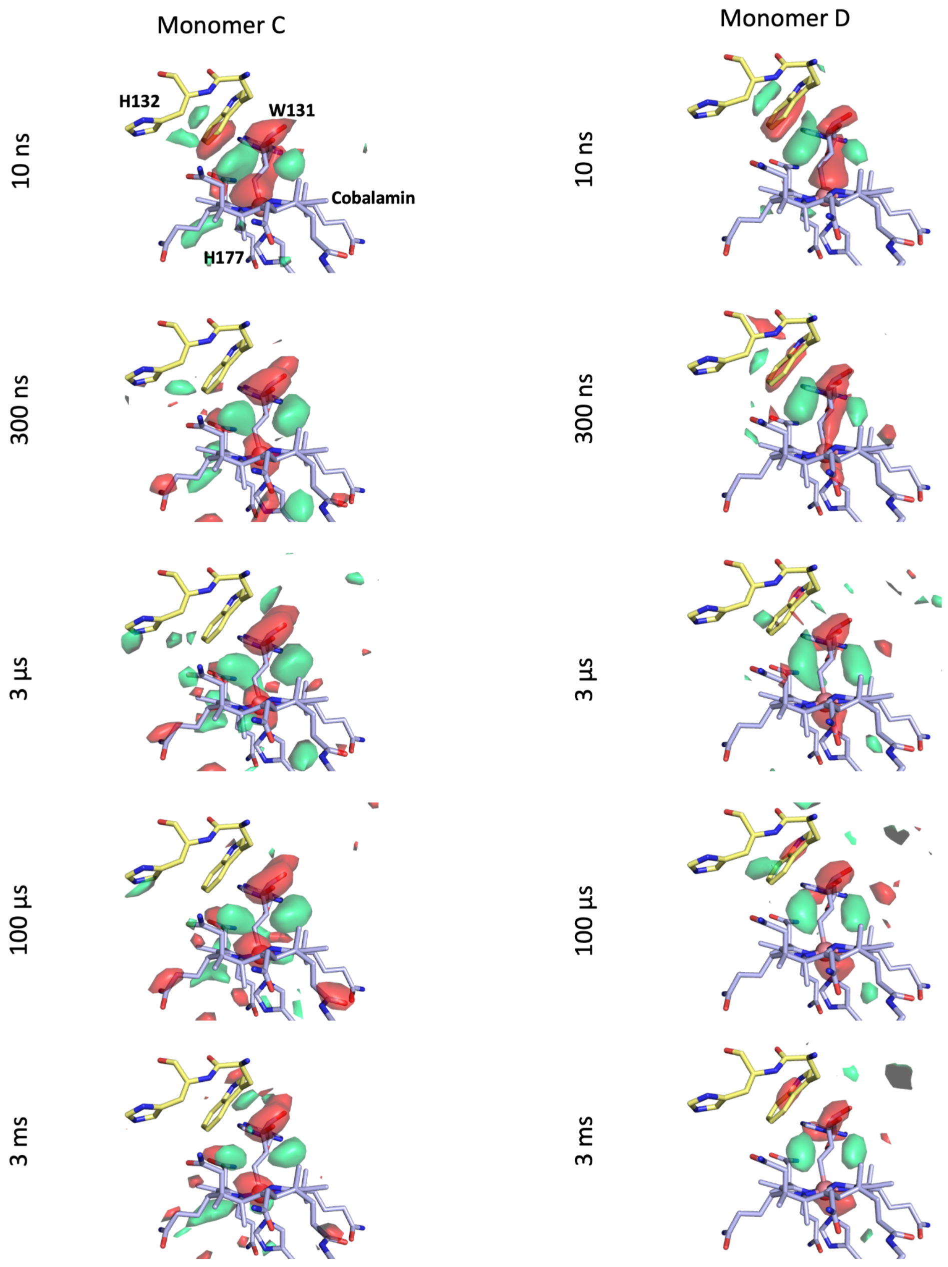


**<u>Supplementary Fig. 8. Signal in the Fourier difference maps from the time-series collected at SACLA observed in the chromophore binding pocket.</u>**

Close-up view of the chromophore region of *Tt*CBD. Fourier difference maps $F_o^{\mathbf{\Delta}t\_30\ mJ/cm2\_SACLA} - F_o^{dark\_ref\_SACLA}$ (**Δ**t: 10 ns, 300 ns, 3 µs, 100 µs, 3 ms) are contoured at +3.5 $\sigma$ (green) and -3.5 $\sigma$ (red). Residues H132 and W131 from the four-helix bundle domain are shown as yellow sticks, while H177 (from the Rossmann fold) and the cobalamin are represented as blue sticks.

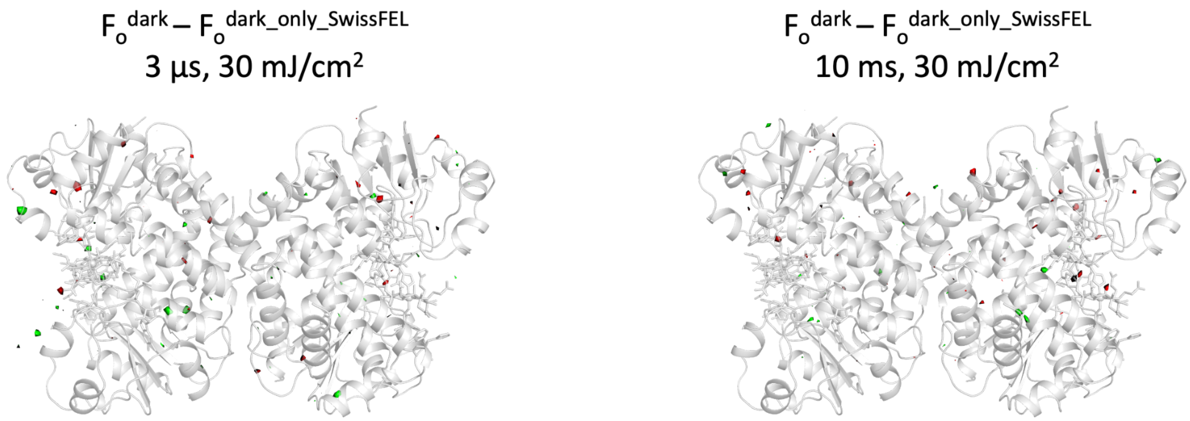


**<u>Supplementary Fig. 9. Fourier difference maps calculated between interleaved darks and dark_only data sets collected at SwissFEL</u>**

Fourier difference maps $F_o^{dark(\Delta t)} - F_o^{dark\_only\_SwissFEL}$ (**Δ**t: 3 µs, 10 ms) are contoured at +3.5 (green) and -3.5 σ (red). The dark data sets were collected in an interleaved way, while collecting light data at 3 µs and 10 ms time delays and with a laser fluence of 30 mJ/cm$^2$. The *Tt*CBD tetramer of the *dark_ref_SwissFEL* model data is overlaid (grey cartoon).

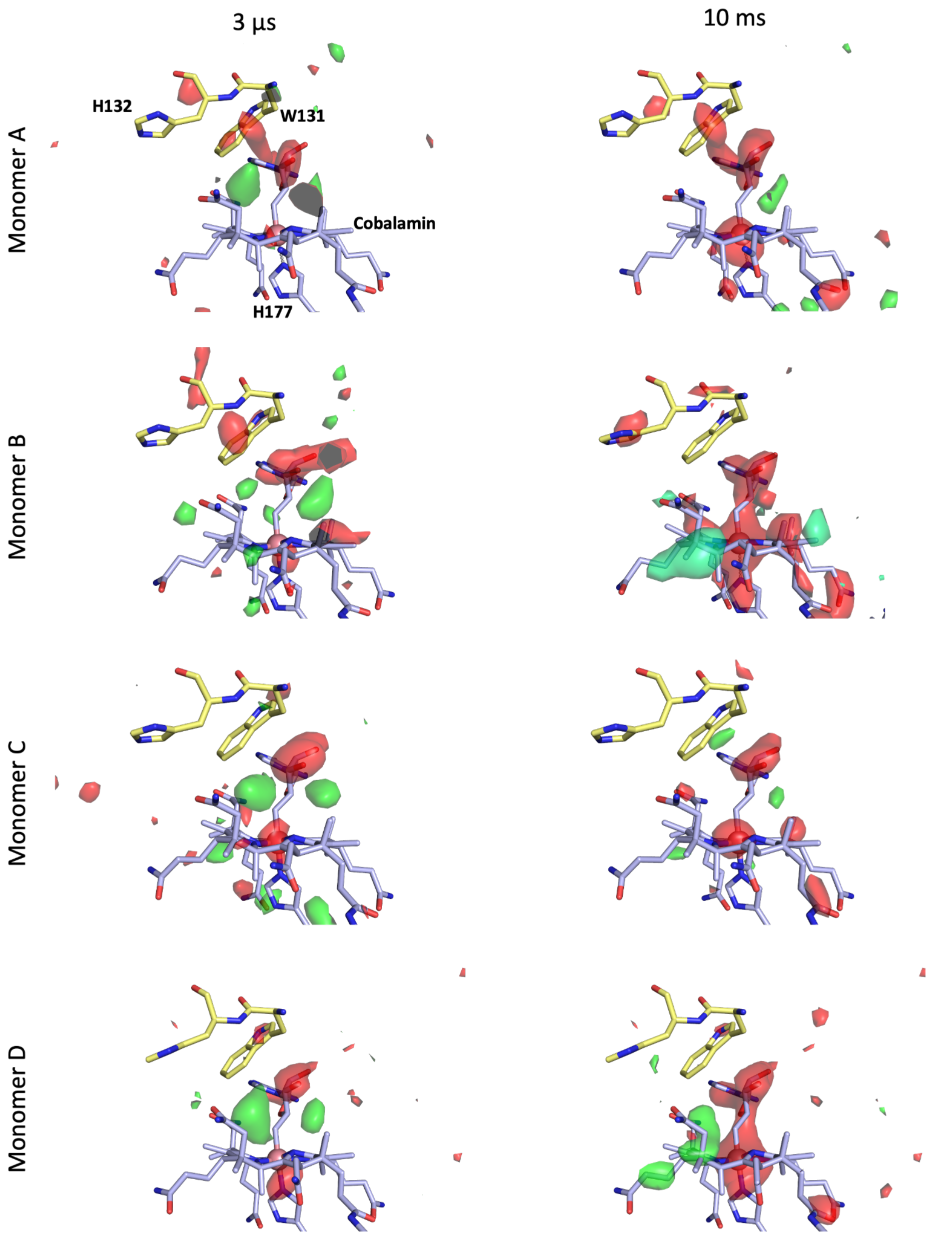


**Supplementary Fig. 10. Fourier difference maps of the 3 µs and 10 ms time delays at a pump-laser fluence of 30 mJ/cm² (SwissFEL).**

Close-up view in the chromophore region for the different monomers. *Dark_ref_SwissFEL* model is represented in yellow (protein) and blue (AdoCbl) sticks. Fourier difference maps $F_o^{\Delta t\_30mJ/cm2\_SwissFEL} - F_o^{dark\_ref\_SwissFEL}$ (**Δ**t: 3 µs, left and 10 ms, right) are contoured at +3.5 (green) and -3.5 σ (red).

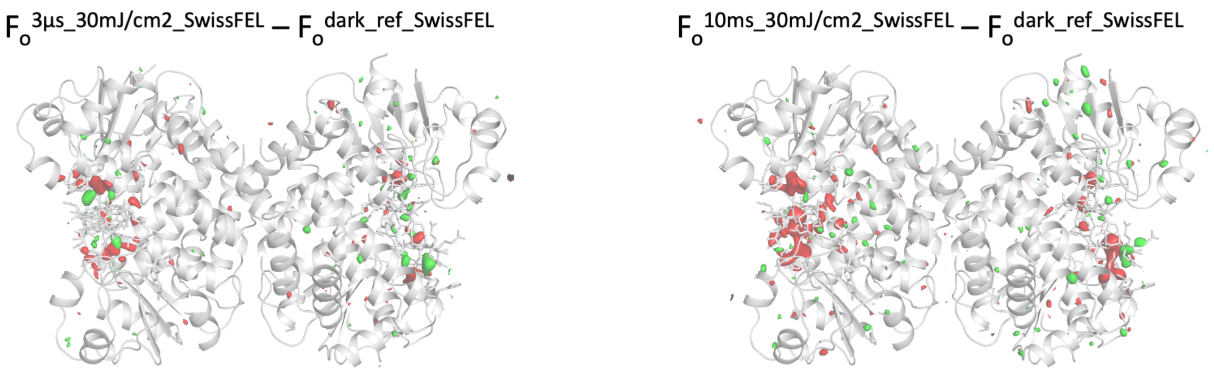


**Supplementary Fig. 11. Fourier difference maps of 3 µs and 10 ms time delays at a laser fluence of 30 mJ/cm$^2$ (SwissFEL).**

Model of *dark_ref_SACLA Tt*CBD tetramer is represented as a grey cartoon. Fourier difference maps $F_o^{\Delta t\_30mJ/cm2} - F_o^{dark\_ref\_SwissFEL}$ (**Δ**t: 3 µs, 10 ms) are contoured at +3.5 (green) and -3.5 σ (red).

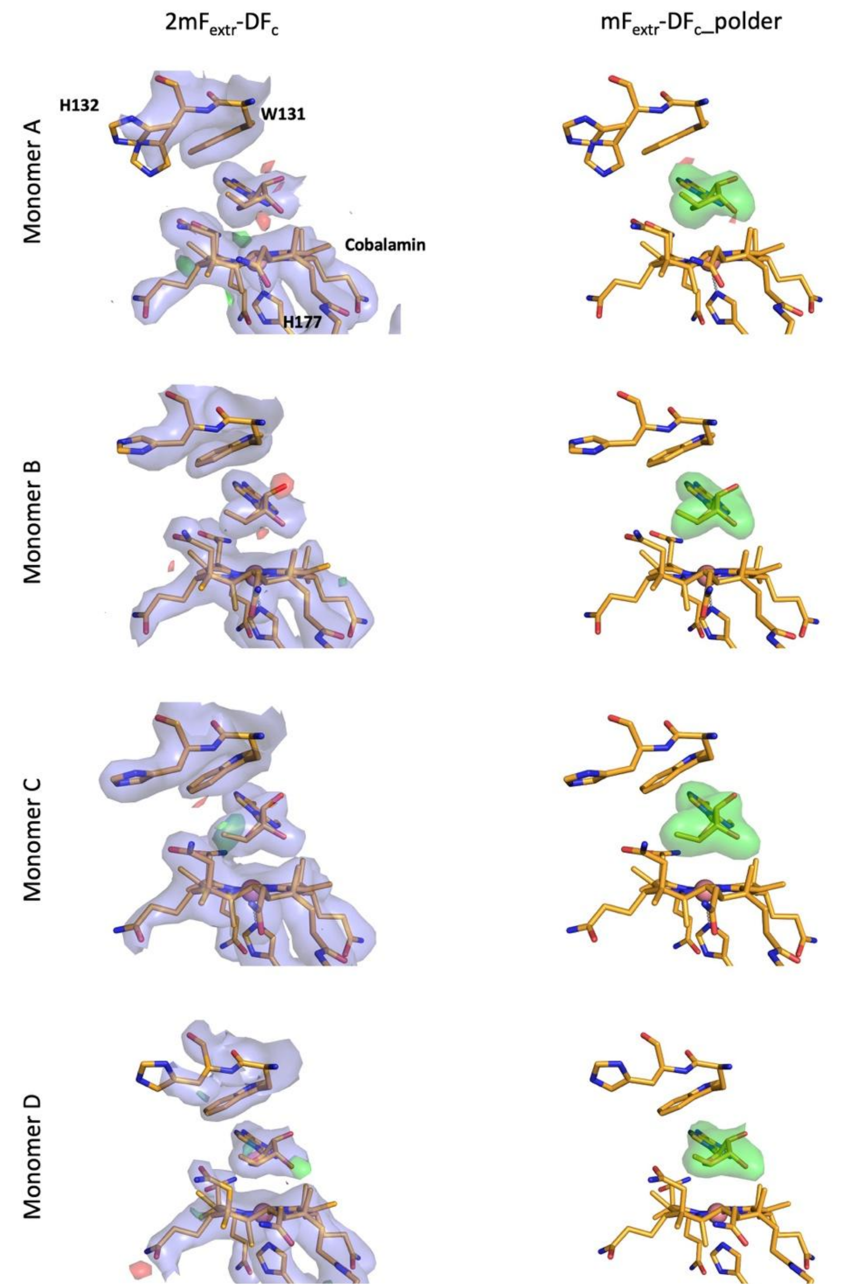


**<u>Supplementary Fig. 12. *10ns_30mJ/cm²_SACLA* intermediate-state model and 2mF$_{extr}$-DF$_c$ electron density and mF$_{extr}$-DF$_c$ polder maps (adenosyl moiety omitted).</u>**

Close-up view of the extrapolated electron density maps in the chromophore region and mF$_{extr}$-DF$_c$_polder map omitting the adenosyl group for each monomer. The 10 ns intermediate-state model is represented in orange sticks. Extrapolated structure factor amplitudes were calculated with an $\alpha$ value of 3.6 (28% occupancy, occupancy assumed to be $1/\alpha$). 2mF$_{extr}$-DF$_c$ is contoured at +1 σ, and mF$_{extr}$-DF$_c$ at +3.5 (green) and -3.5 σ (red). mF$_{extr}$-DF$_c$_polder map is shown at +3.5 (green) and -3.5 σ (red).

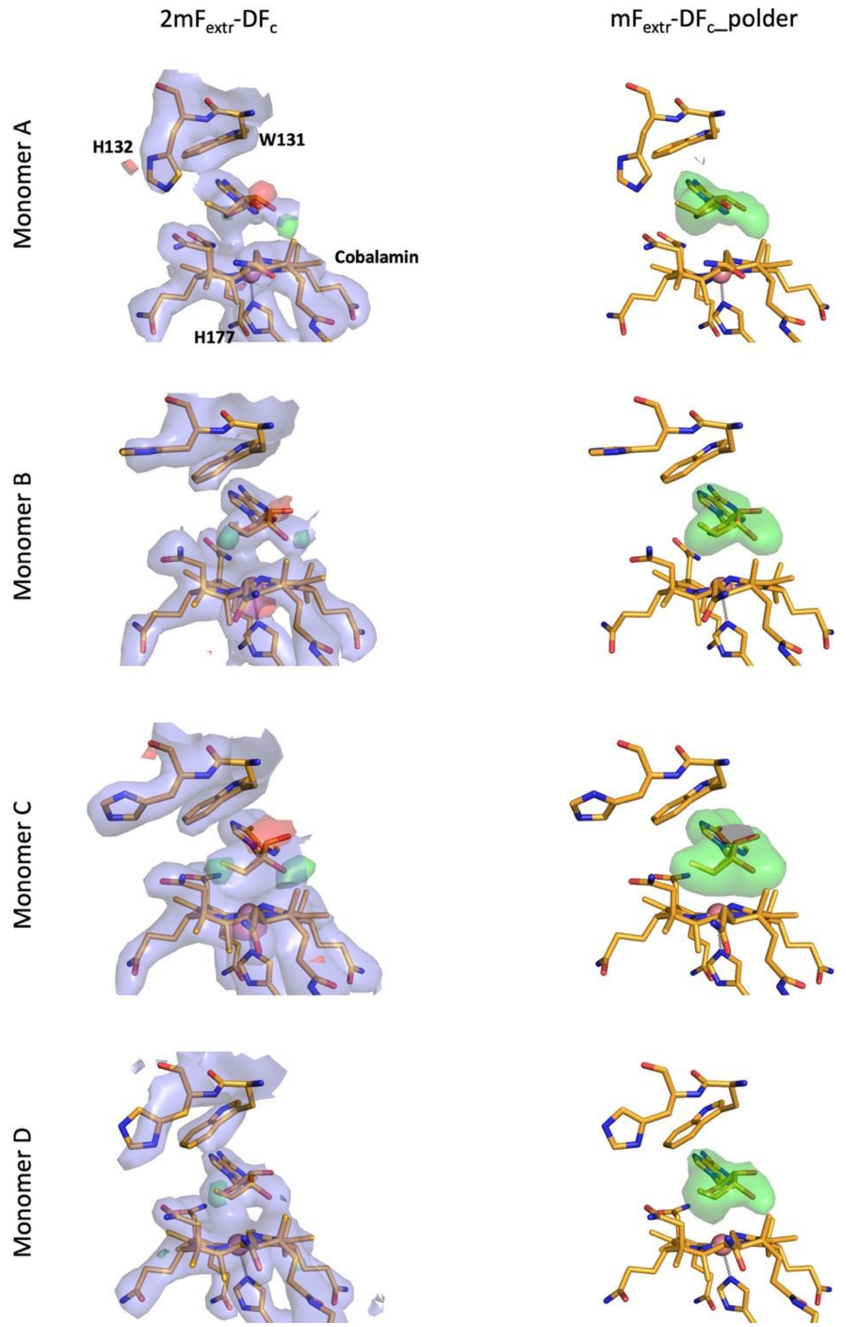


**Supplementary Fig. 13. *300ns_30mJ/cm² _SACLA* intermediate-state model and 2mF$_{extr}$-DF$_c$ electron density and mF$_{extr}$-DF$_c$ polder maps (omitted adenosyl moiety).**

Close-up view of the extrapolated electron density maps in the chromophore region and mF$_{extr}$-DF$_c$_polder map omitting the adenosyl group for each monomer. The 300 ns intermediate-state model is represented in orange sticks. Extrapolated structure factor amplitudes were calculated with an $\alpha$ value of 3.3 (30% occupancy). 2mF$_{extr}$-DF$_c$ map is contoured at +1 $\sigma$, and mF$_{extr}$-DF$_c$ at +3.5 (green) and -3.5 $\sigma$ (red). mF$_{extr}$-DF$_c$_polder map is shown at +3.5 (green) and -3.5 $\sigma$ (red).

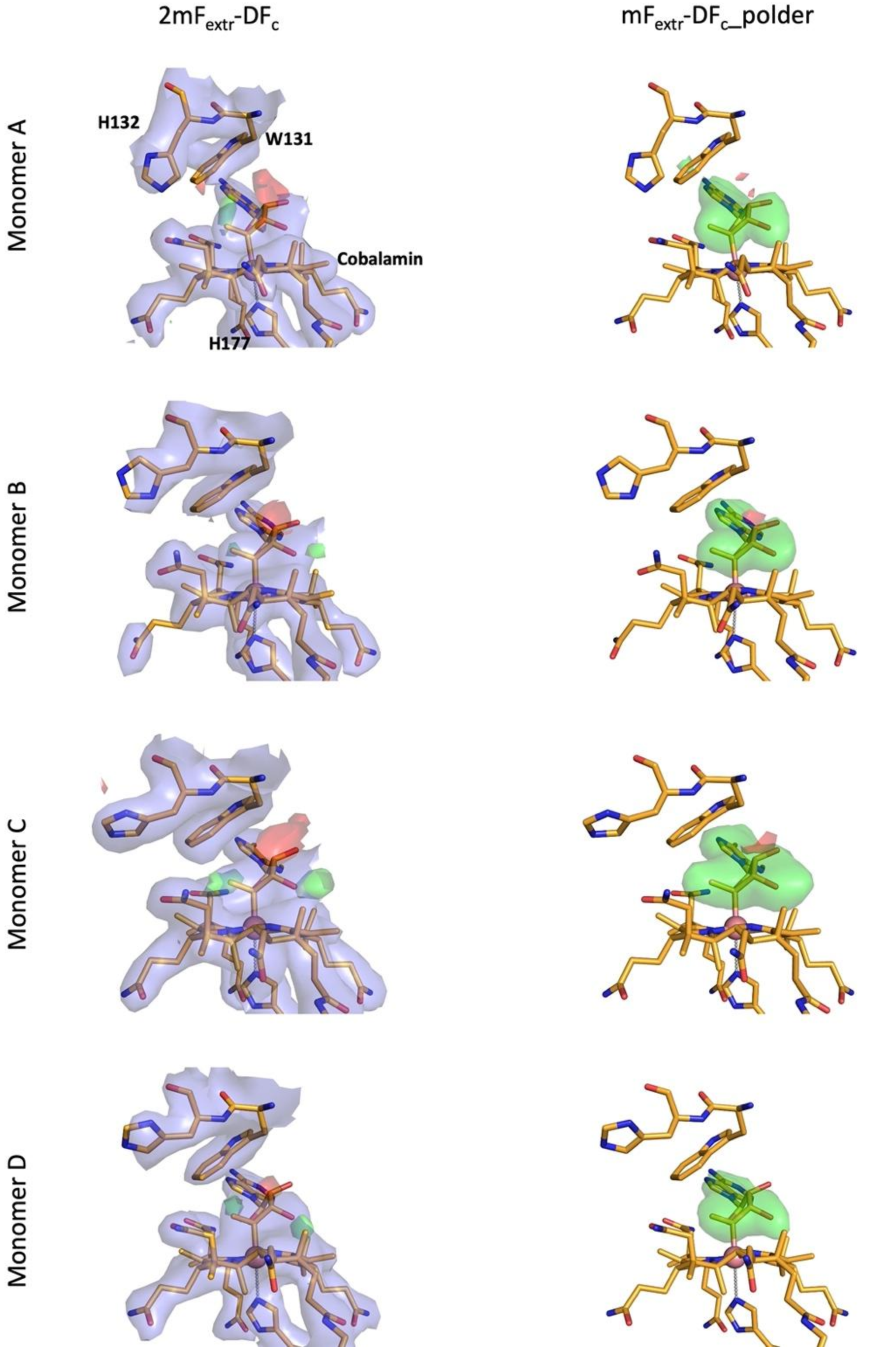


**<u>Supplementary Fig. 14. *3μs_30mJ/cm[2]_SACLA* intermediate-state model and $2mF_{extr}$-$DF_c$ electron density and $mF_{extr}$-$DF_c$ polder maps (omitted adenosyl moiety).</u>**

Close-up view of the extrapolated electron density map in the chromophore region and $mF_{extr}$-$DF_c$_polder map omitting the adenosyl group for each monomer. The 3 μs intermediate-state model is represented in orange sticks. Extrapolated structure factor amplitudes were calculated with an $\alpha$ value of 3.6 (28% occupancy). $2mF_{extr}$-$DF_c$ contoured at +1 σ, and $mF_{extr}$-$DF_c$ at +3.5 (green) and -3.5 σ (red). $mF_{extr}$-$DF_c$_polder map is shown at +3.5 (green) and -3.5 σ (red) sigma.

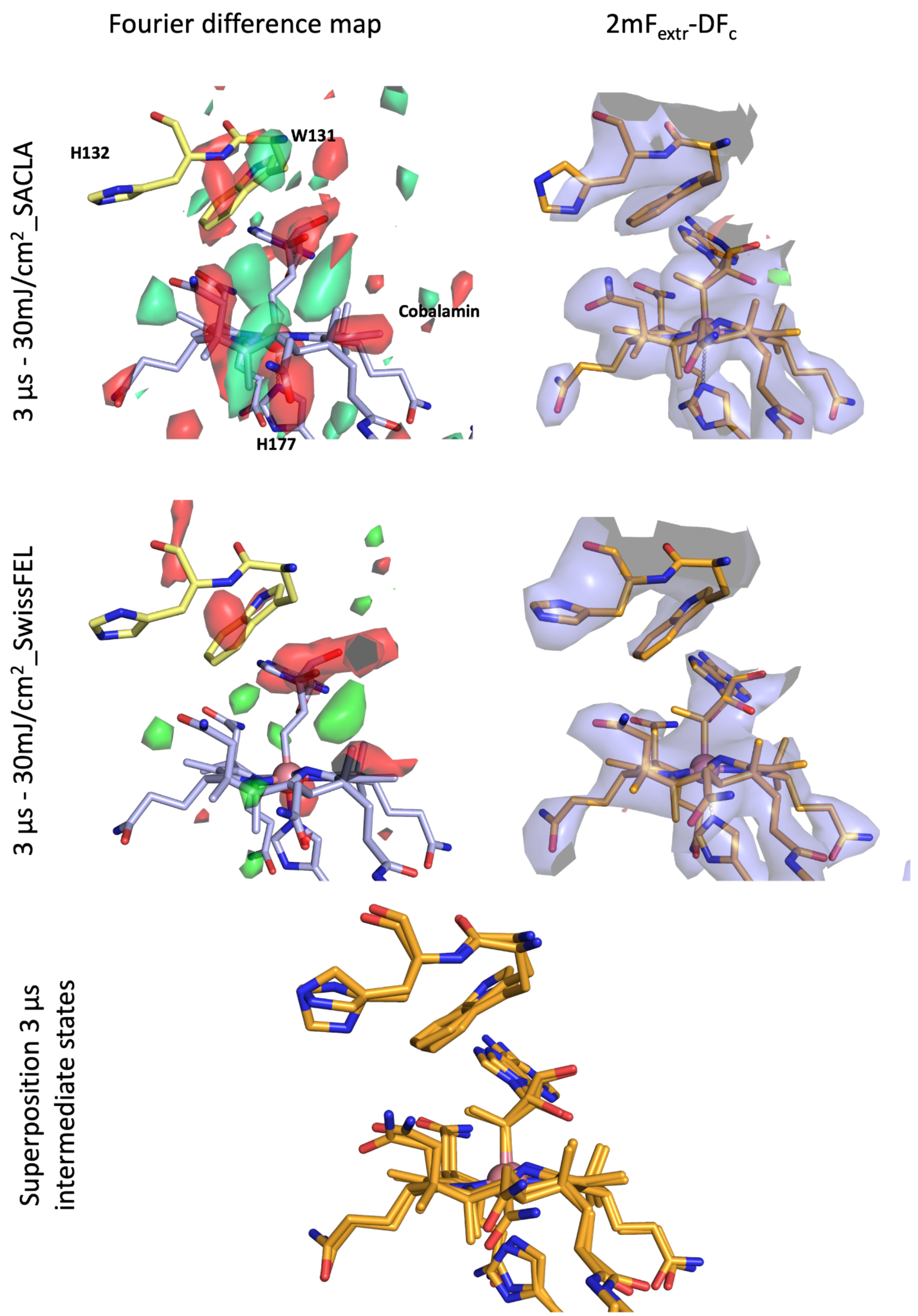


**Supplementary Fig. 15. Comparison of Fourier difference maps, intermediate-state models and 2mF$_{extr}$-DF$_c$ electron density maps of *3µs_30mJ/cm²_SACLA* and *3µs_30mJ/cm²_SwissFEL***

Close-up view of the Fourier difference maps $F_o^{3µs\_30mJ/cm2\_SACLA} - F_o^{dark\_ref\_SACLA}$ and $F_o^{3µs\_30mJ/cm2\_SwissFEL} - F_o^{dark\_ref\_SwissFEL}$ in the chromophore region of monomer B (left, contoured at +3.5 σ (green) and -3.5 σ (red)). The extrapolated electron density maps ($2mF_{extr}$-$DF_c$ and $mF_{extr}$-$DF_c$ ) are displayed on the right with the intermediate-state model represented in orange sticks. $2mF_{extr}$-$DF_c$ is contoured at +1 σ, and $mqF_{extr}$-$DF_c$ at +3.5 σ (green) and -3.5 σ (red). Bottom, SwissFEL intermediate-state model has been superimposed onto SACLA intermediate-state model using Cα atoms of monomer B (RSMD of 0.340 Å). The conformations of the adenosyl group in the two models are very similar.

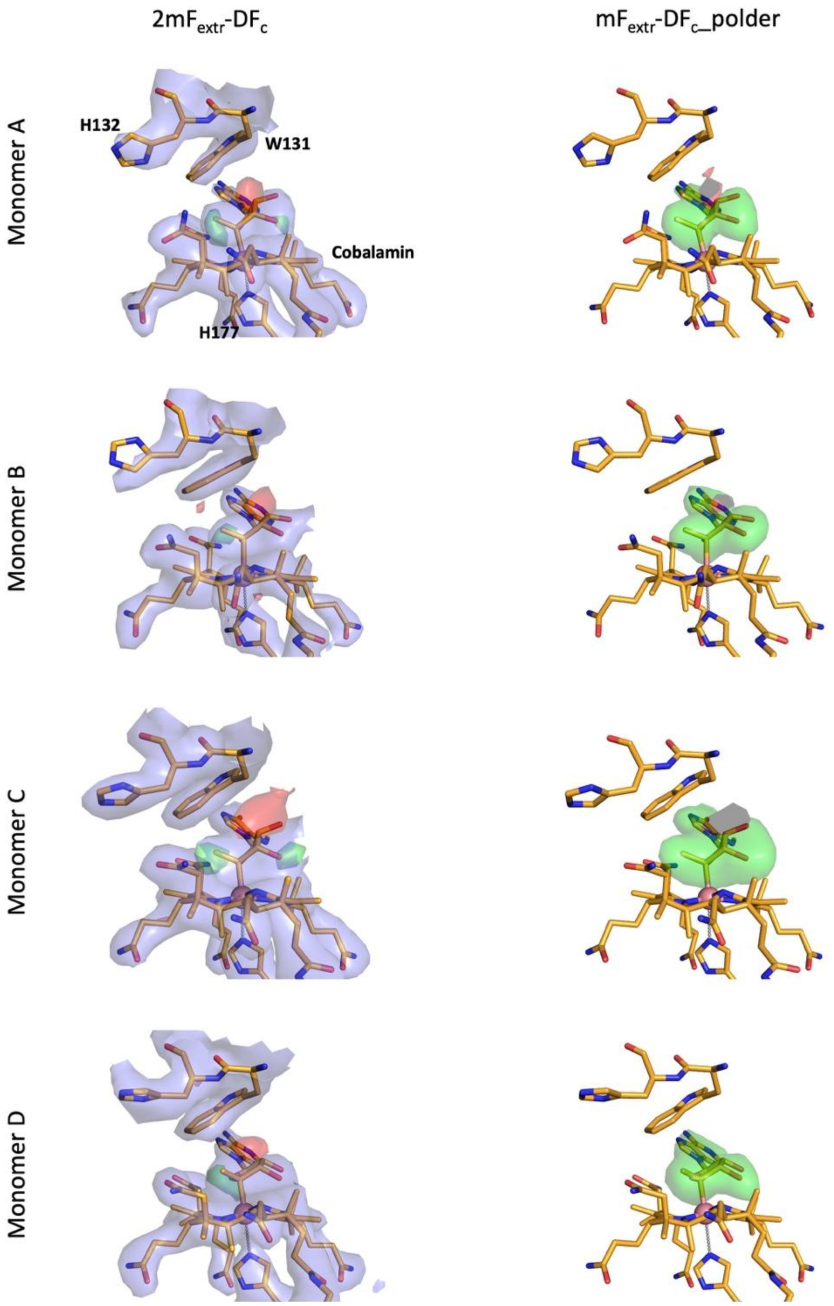


**Supplementary Fig. 16.** ***100µs_30mJ/cm²_SACLA*** **intermediate-state model and $2mF_{extr}$-$DF_c$ electron density and $mF_{extr}$-$DF_c$ polder maps (omitted adenosyl moiety).**

Close-up view of the extrapolated electron density map in the chromophore region and $mF_{extr}$-$DF_c$ polder map omitting the adenosyl group for each monomer. The 100 µs intermediate-state model is represented in orange sticks. Extrapolated structure factor amplitudes were calculated with an $\alpha$ value of 3.3 (30% occupancy). $2mF_{extr}$-$DF_c$ contoured at +1 σ, and $mF_{extr}$-$DF_c$ at +3.5 (green) and -3.5 σ (red). $mF_{extr}$-$DF_c$_polder map is shown at +3.5 (green) and -3.5 σ (red).

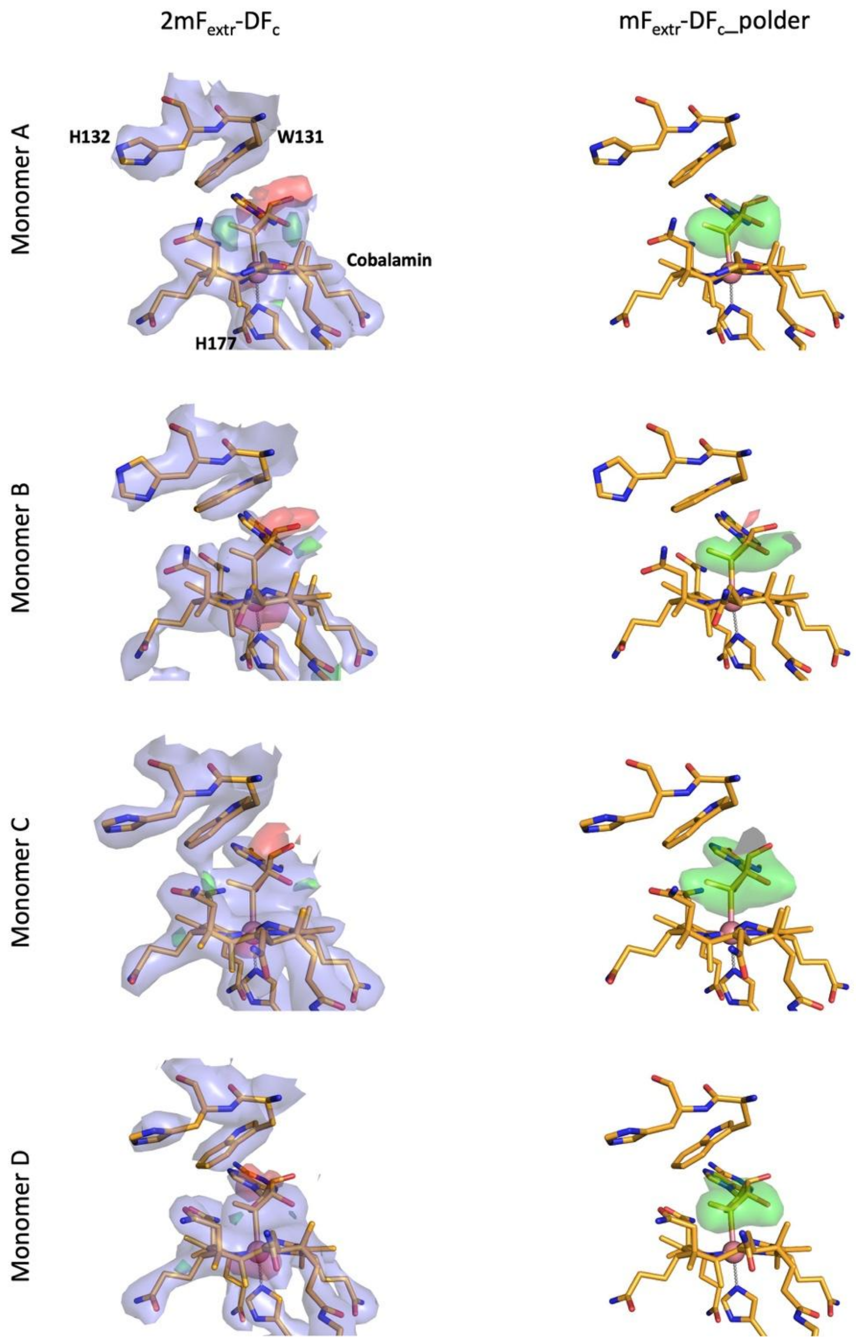


**Supplementary Fig. 17.** ***3ms_30mJ/cm²_SACLA*** **intermediate-state model and $2mF_{extr}$-$DF_c$ electron density and $mF_{extr}$-$DF_c$ polder maps (omitted adenosyl moiety).**

Close-up view of the extrapolated electron density maps (left, $2mF_{extr}$-$DF_c$ and $mF_{extr}$-$DF_c$ ) in the chromophore region and $mF_{extr}$-$DF_c$_polder map omitting the adenosyl group for each monomer. The 3 ms intermediate-state model is represented in orange sticks. Extrapolated structure factor amplitudes were calculated with an $\alpha$ value of 3.3 (30% occupancy). $2mF_{extr}$-$DF_c$ contoured at +1 σ, and $mF_{extr}$-$DF_c$ at +3.5 (green) and -3.5 σ (red). $mF_{extr}$-$DF_c$_polder map is shown at +3.5 (green) and -3.5 σ (red).

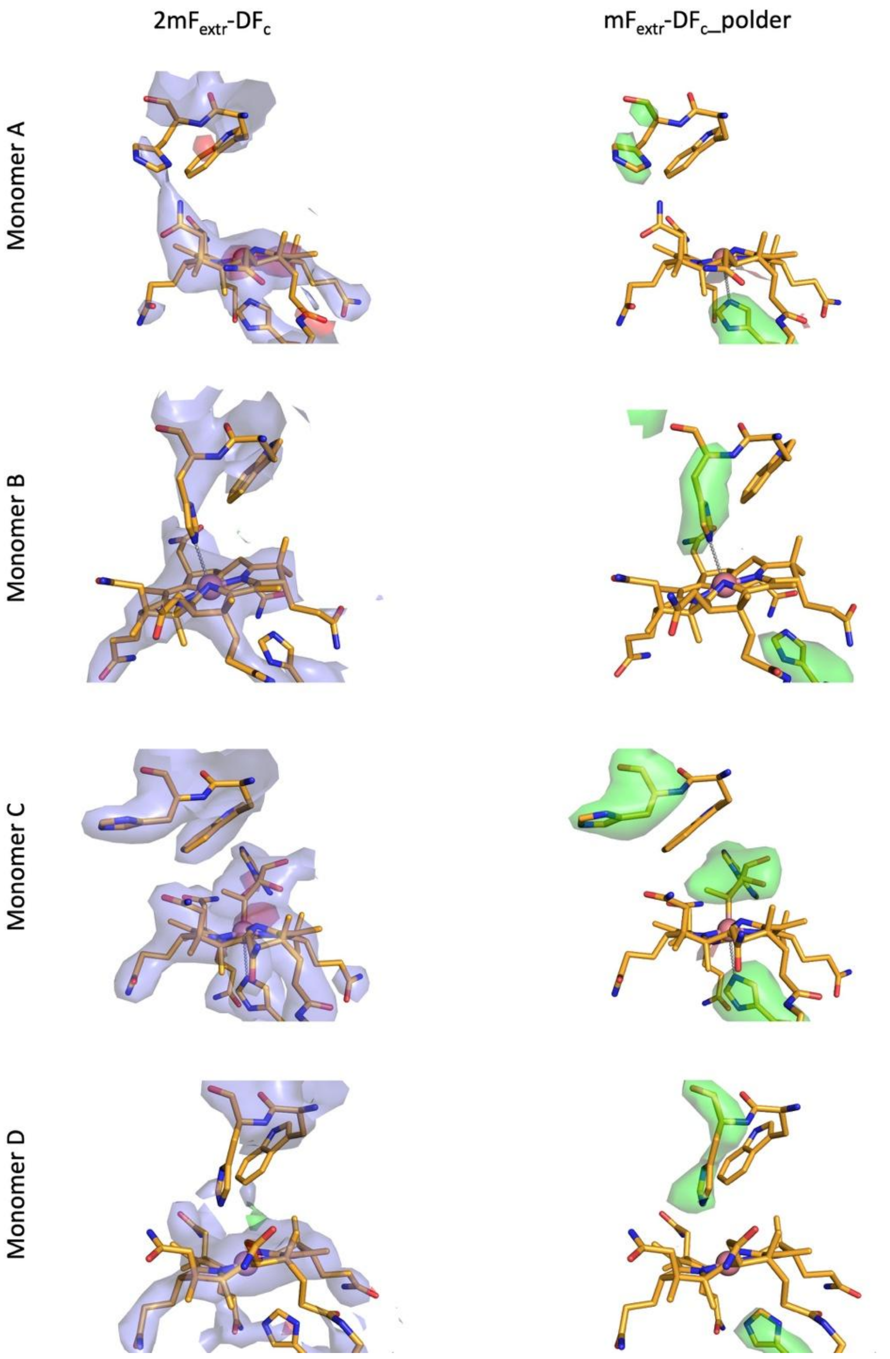


**Supplementary Fig. 18.** ***10ms_30 mJ/cm²_SwissFEL*** **intermediate-state model and $2mF_{extr}$-$DF_c$ electron density and $mF_{extr}$-$DF_c$ polder maps (adenosyl moiety omitted).**

Close-up view of the extrapolated electron density maps (left, $2mF_{extr}$-$DF_c$ and $mF_{extr}$-$DF_c$) in the chromophore region and $mF_{extr}$-$DF_c$_polder map omitting the adenosyl group for each monomer (right). The 10 ms intermediate-state model is represented in orange sticks. Extrapolated structure factor amplitudes were calculated with an $\alpha$ value of 3.3 (30% occupancy). $2mF_{extr}$-$DF_c$ contoured at +1 σ, and $mF_{extr}$-$DF_c$ at +3.5 (green) and -3.5 σ (red). $mF_{extr}$-$DF_c$_polder map is shown at +3.5 (green) and -3.5 σ (red).

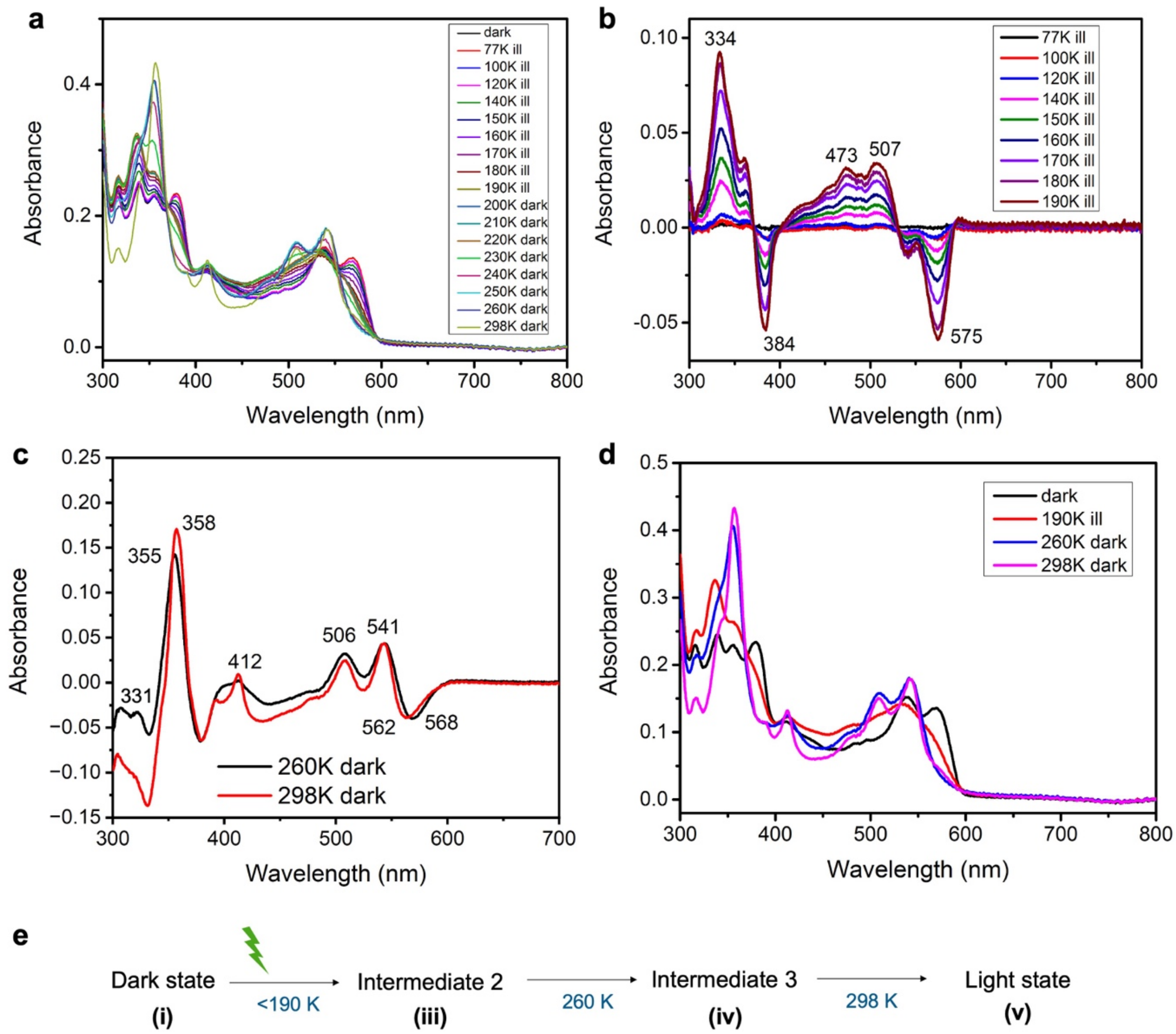


**<u>Supplementary Fig. 19. Cryotrapping of intermediate states in the photochemical reaction of *Tt*CBD in solution as monitored by temperature-resolved absorption spectroscopy measurements.</u>**

(**a**) 77 K absorbance spectra of *Tt*CBD in solution (50 µM) after illumination (530 nm) for 15 min at different temperatures ranging from 77 K to 190 K or after illumination at 190 K for 15 min and incubation in the dark for 15 min at increasing temperatures. (**b**) Difference spectra after successive 15 min illumination at increasing temperatures from 77 to 190 K. The non-illuminated sample was used as a baseline. (**c**) Difference spectra after illumination at 190 K for 15 min and incubation in the dark for 15 min at either 260 or 298 K. (**d**) 77 K absorbance spectra of the dark, light (298 K dark) and intermediate states (190 K and 260 K) identified by cryogenic absorbance measurements at 77 K. The wavelengths of the main spectral changes are labelled accordingly in panels B and C. (**e**) The temperature-resolved absorption spectroscopy data yield the reaction scheme shown here, indicating the temperatures at which each of the intermediate states is formed. The i, iii, iv and v labels illustrate how these species relate to those depicted in Fig. 1 of the main text.

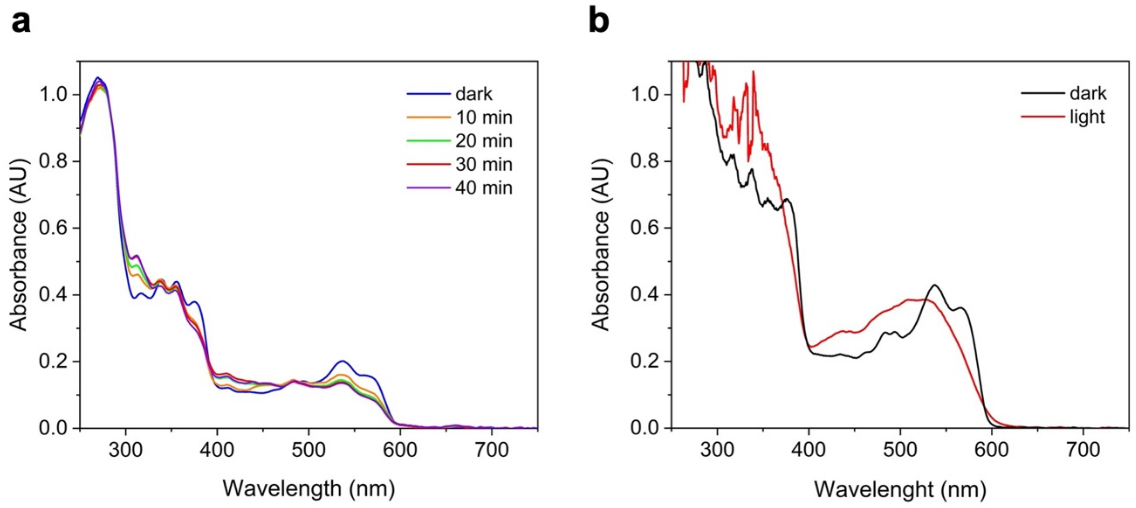


**Supplementary Fig. 20. *In crystallo* UV-vis spectroscopy on *Tt*CBD CarH at BM07-FIP2 beamline (ESRF).**

(**a**) Absorption spectra (normalized at 280 nm) of crystalline *Tt*CBD after consecutive illumination periods of 10 min at 100 K performed with an optical fibre-coupled LED at 530 nm. (**b**) Photoconversion of a *Tt*CBD crystal shown by comparison of the absorption spectra (normalized at 280 nm) of the dark state (black) and of the crystal after 6-min illumination performed at 180 K (red).

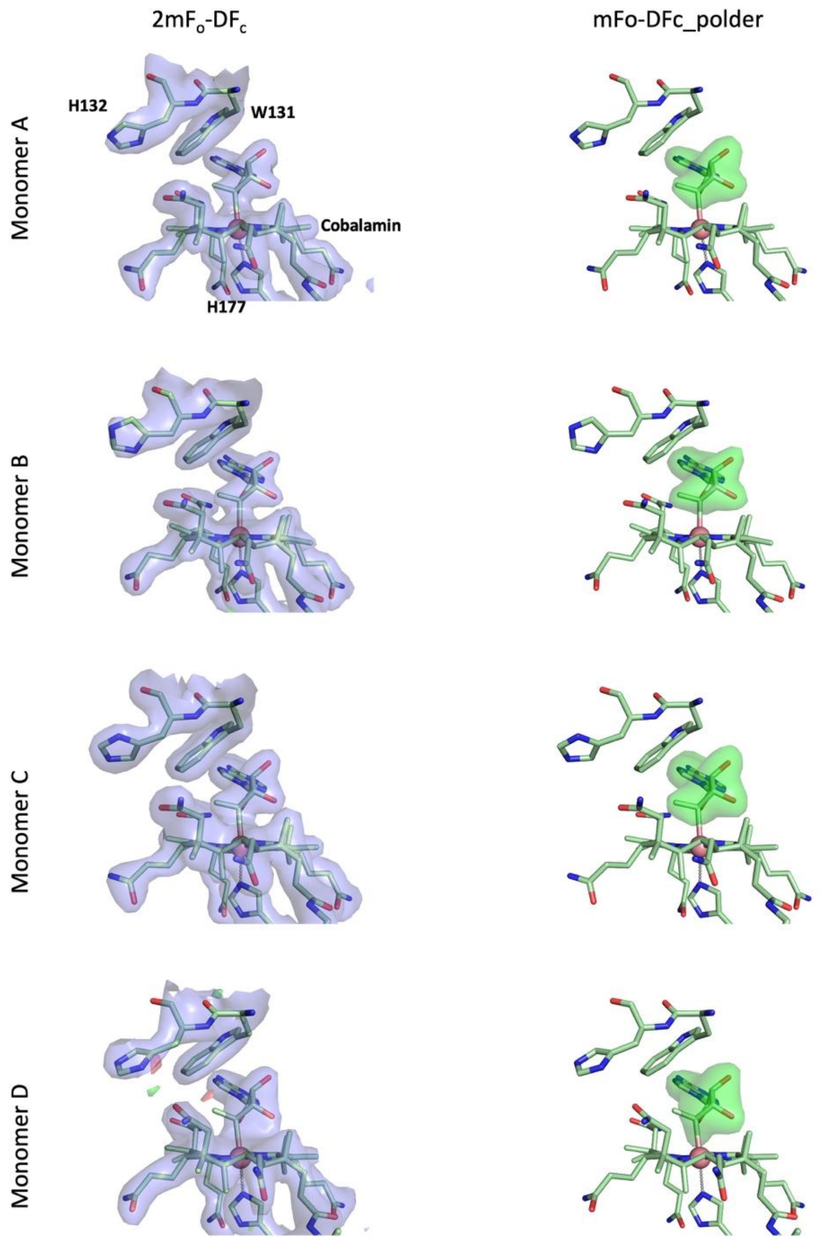


**<u>Supplementary Fig. 21. *Cryo_light_180K* model and $2mF_o$-$DF_c$ electron density and $mF_o$-$DF_c$ polder maps (adenosyl moiety omitted).</u>**

Close-up view of the electron density maps (left, $2mF_o$-DF and $mF_o$-$DF_c$) in the chromophore region and $mF_o$-$DF_c$_polder map omitting the adenosyl group for each monomer (right). The cryo-trapped intermediate state model is represented in pale green sticks. $2mF_o$-$DF_c$ is contoured at +1 σ, and $mF_o$-$DF_c$ at +3.5 σ (green) and -3.5 σ (red). $mF_o$-$DF_c$_polder map is shown at +3.5 σ (green) and -3.5 σ (red).

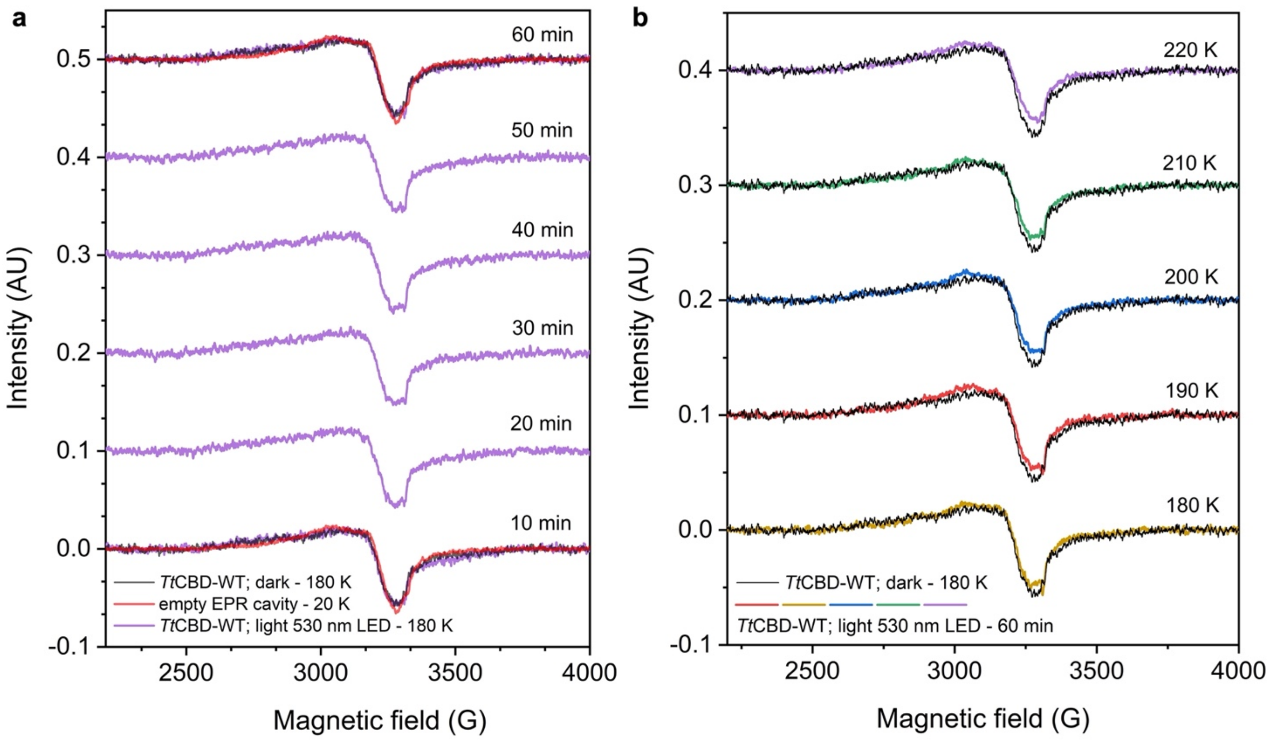


**Supplementary Fig. 22. Characterization of intermediate 2 state in the photochemical reaction of aerobic *Tt*CBD in solution as monitored by temperature-resolved EPR spectroscopy measurements**

(**a**) *In-situ* illumination of aerobic *Tt*CBD in solution at 180 K coupled with cw-EPR data collection at 20 K. The spectra show no paramagnetic EPR signals over the illumination period of 0-60 min, implying Co(III) redox-state for this species. (**b**) EPR spectra recorded at a range of temperatures between 180 K and 220 of the *Tt*CBD sample following illumination at 180 K for 60 mins in solution. No change in the Co redox state is observed, when the spectra were measured between 180 and 220 K, suggesting that no coupled Co(II)-radical species are present.

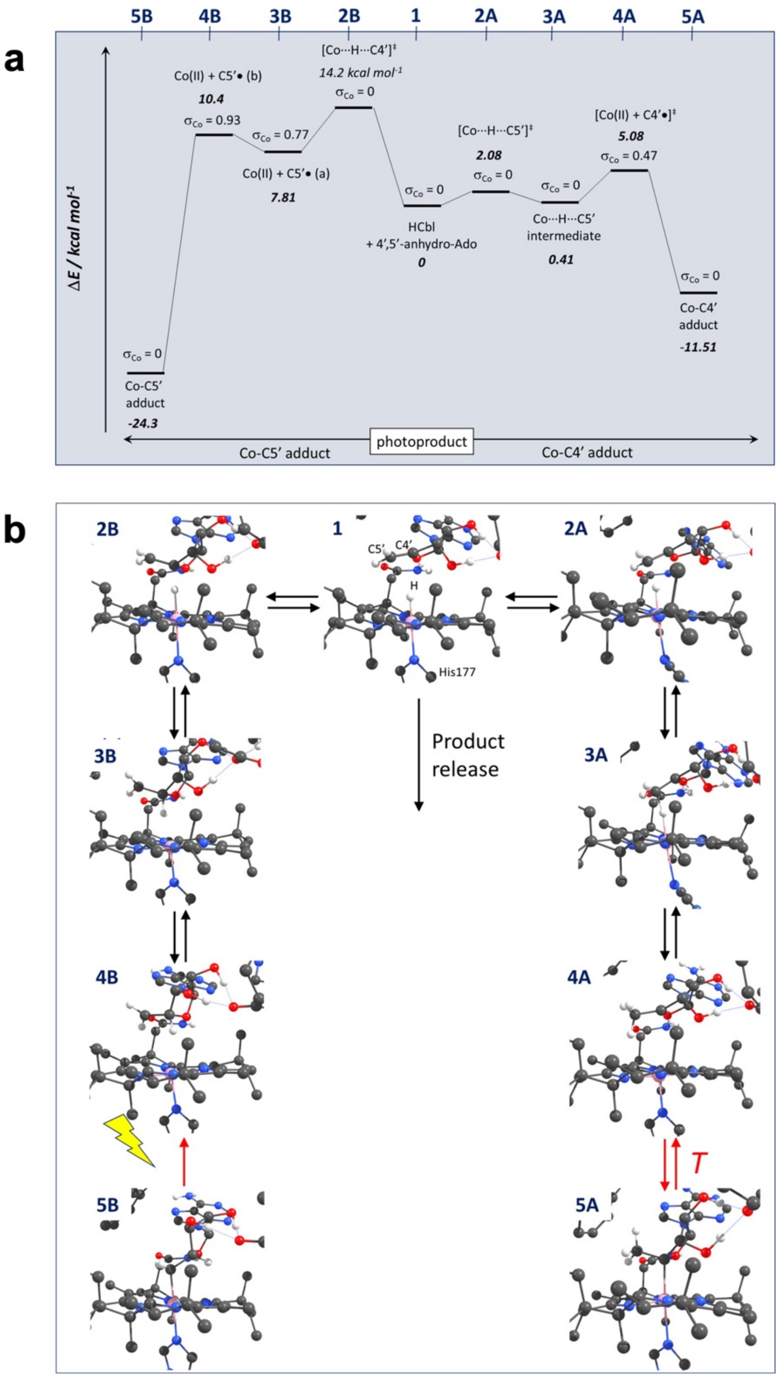


**Supplementary Fig. 23. Proposed mechanism for the interconversion of the Co–C4′ and Co–C5′ adducts via the photoproduct based on cluster model calculations.**

(**a**) Potential energy profile with energy in kcal mol$^{-1}$ for each species (bold) and the spin density on the Co, $\sigma_{Co}$. (**b**) Structures along the mechanism indicating that Co–C5′ bond cleavage in the dark state (5B) requires photoactivation while Co–C4′ bond cleavage in the C4′ adduct (5A) may occur thermally. Release of photoproduct 1 from the binding pocket shifts the equilibrium away from the C4′ adduct (5A).

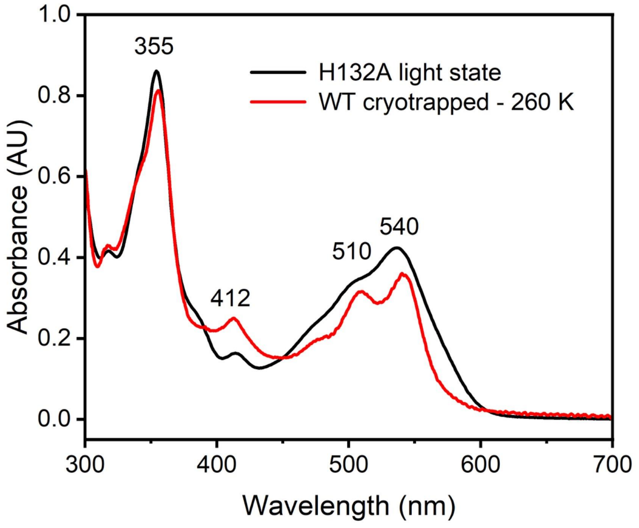


**Supplementary Fig. 24. Absorption solution spectra of the *Tt*CBD H132A mutant light state at room temperature and of the *Tt*CBD intermediate 2 cryotrapped at 260 K.**
Spectroscopic intermediate measured at 77 K after illumination with a 530 nm LED at 190 K for 15 min and incubation in the dark for 15 min at 260 K (red curve) and spectrum of the *Tt*CBD H132A mutant light state measured at room temperature after illumination with a 530 nm LED until complete photoconversion (black curve).

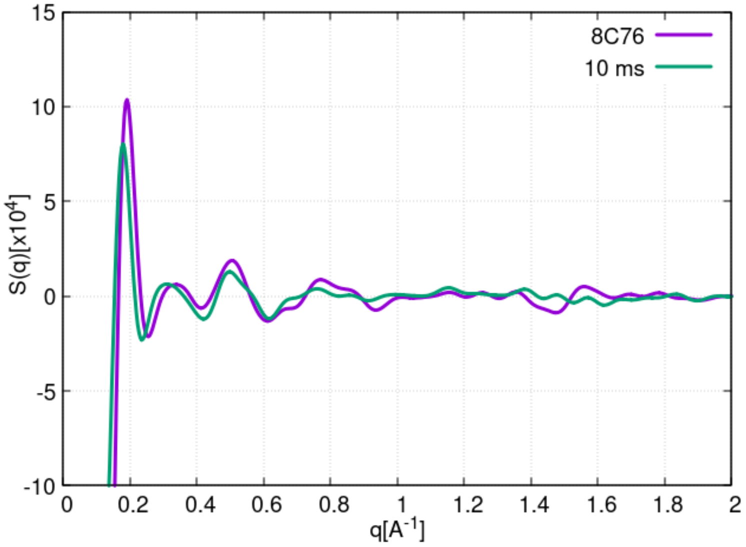


**Supplementary Fig. 25. Comparison of the XSS signals calculated using the cryotrapped 8C76 structure and the SFX 10 ms intermediate.**

The atomic coordinates of the previously reported corrin-ring-shifted *Tt*CBD structure (PDB entry code: 8C76) and the dark-state *Tt*CBD structure (PDB entry code: 8C73[2]) were used to compute the XSS difference signal (shown in purple). A second XSS difference signal (shown in green) was computed using the atomic coordinates of the 10 ms model from SwissFEL (10ms_30mJ/cm²_SwissFEL) and the corresponding dark reference model (dark_ref_SwissFEL). The comparison reveals that both signals contain peaks at similar positions up to ~0.7 $Å^{-1}$, while other features at higher q values (e.g. the positive peak around 0.8 $Å^{-1}$ and the negative one around 1.5 $Å^{-1}$) are lost in the 10 ms signal.

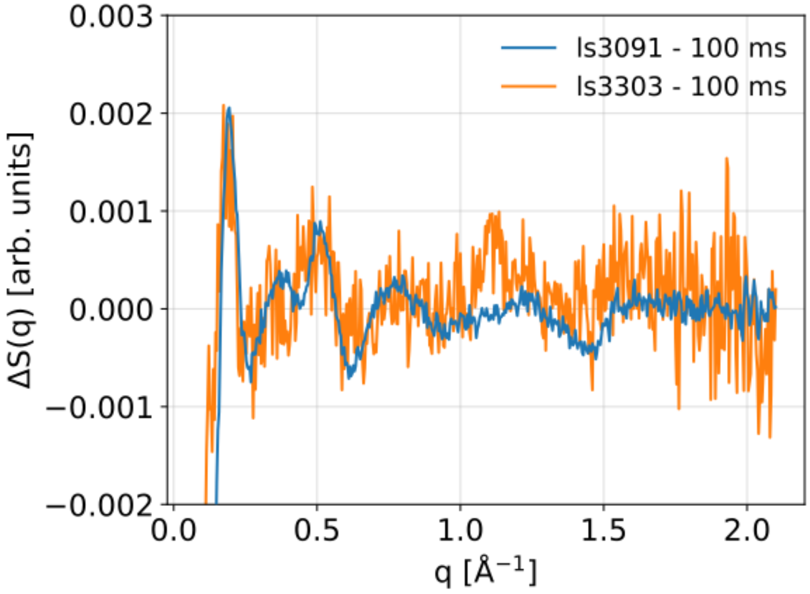


**Supplementary Fig. 26. Comparison of the 100 ms TR-XSS patterns from different data sets.** The TR-XSS pattern at 100 ms from the 30 µs - 100 ms data set (experiment ls3091, blue line) is superimposed on the 100 ms pattern from the 100 ms - 3 s data set (experiment ls3303, orange line). The two data sets were obtained at different protein concentrations and with different data collection protocols, as described in the Material and Methods (see section Time-resolved X-ray solution scattering (TR-XSS): data collection and reduction). The 100 ms overlap time point was measured with both data collection protocols in order to verify that the two data sets yielded TR-XSS patterns with the same shape. Note that the two patterns in the figure have different signal-to-noise ratios due to different data collection parameters (in particular protein concentration, as discussed in the Material and Methods).

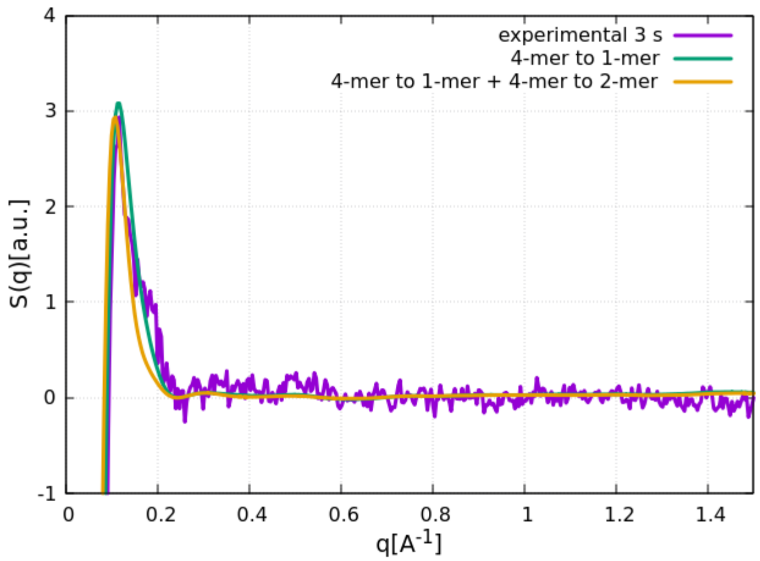


**Supplementary Fig. 27. Effect of tetramer-dimer and tetramer-monomer transitions on the XSS signal.**

A comparison between the XSS difference signals expected for a tetramer-to-monomer transition and for a combination of tetramer-to-monomer (4-mer to 1-mer) and tetramer-to-dimer (4-mer to 2-mer) transitions calculated using the atomic coordinates from the 8C73 PDB and modified versions of the same PDB file, as described in the Material and Methods. The comparison of the two XSS difference signals reveals that both processes induce similar lineshapes, with the tetramer-to-monomer transition being closer to experimental data.

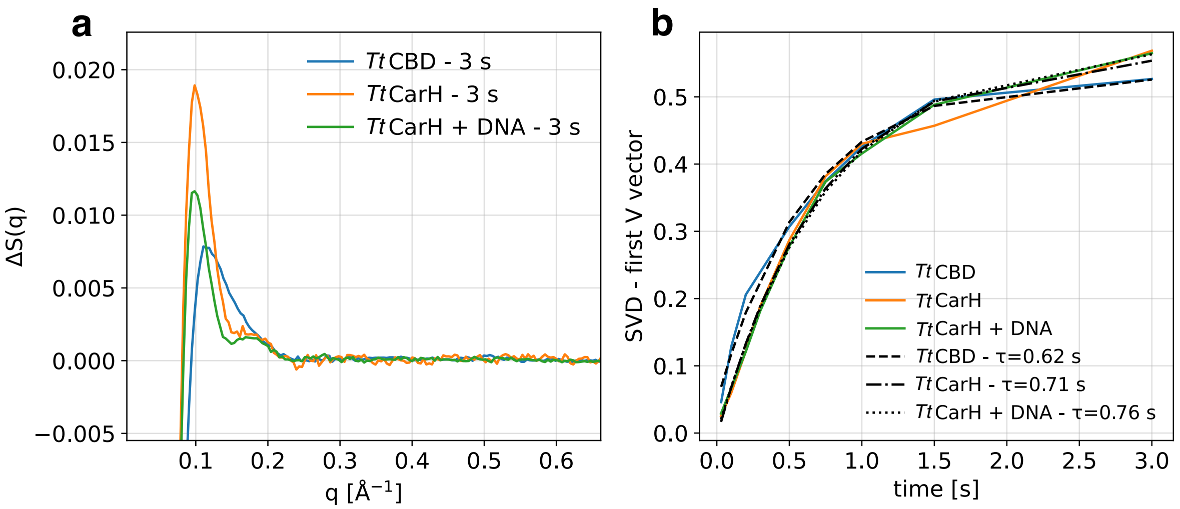


**Supplementary Fig. 28. Longest TR-XSS signal and SVD analysis of TR-XSS data in the 100 ms - 3 s time range.**

(**a**) Experimental scattering patterns of *Tt*CBD (blue), *Tt*CarH (orange) and *Tt*CarH bound to DNA (green) as measured with a laser/X-ray time delay of 3 s. (**b**) Time course of the SVD rSV amplitude from the ms-to-s TR-XSS data sets in *Tt*CBD (blue), *Tt*CarH (orange) and *Tt*CarH bound to DNA (green). The time dependence was fitted using a single exponential model and the result was plotted with the relaxation time given in the legend for *Tt*CBD (dashed line), *Tt*CarH (dash-dotted line) and *Tt*CarH bound to DNA (dotted line).

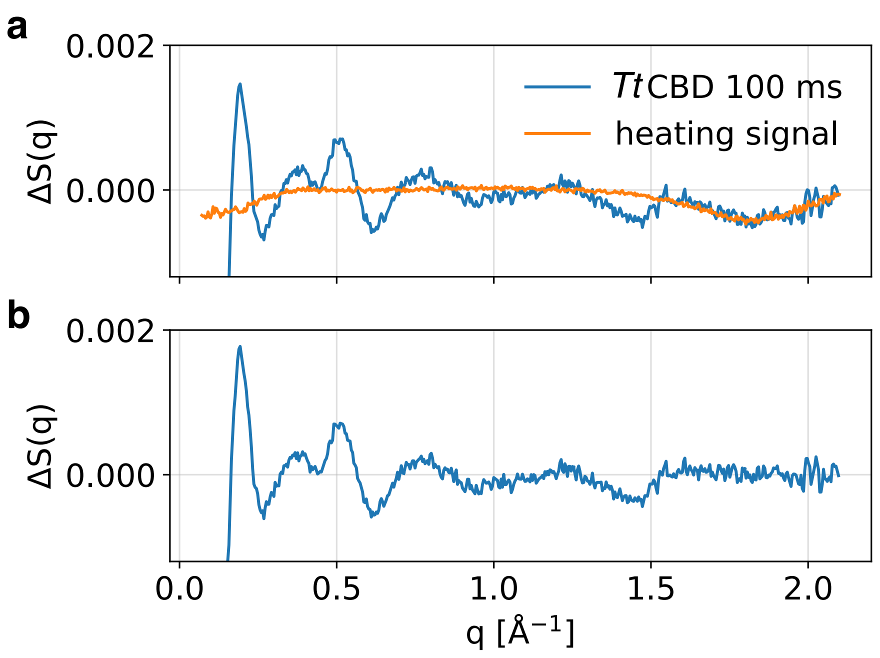


**Supplementary Fig. 29. Removal of solvent heating signal from TR-XSS data.**

(**a**) TR-XSS signal from the *Tt*CBD sample at 100 ms time delay (blue line), and scaled heating signal obtained with a Fast Yellow dye (orange line). (**b**) Resulting signal after subtraction of the scaled heating signal.

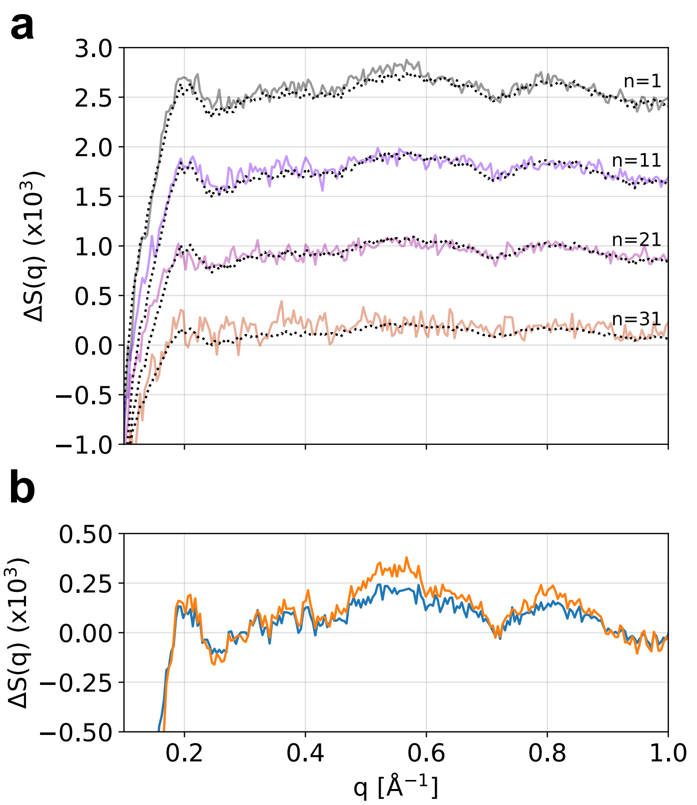


**Supplementary Fig. 30. Correction of TR-XSS data for the irreversible photoconversion kinetics.**

(**a**) Unaveraged TR-XSS difference patterns (coloured continuous lines) measured at the same time delay (1 ms) after 1, 11, 21, or 31 laser/X-ray pulse pairs. The total signal amplitude decays due to the increase of irreversibly photoconverted sample. The SVD reconstructed patterns (using only a single SVD component) are overlaid (black dotted lines). (**b**) Comparison of the average over the TR-XSS difference patterns of panel A before (blue line) and after (orange line) correction for irreversible photoconversion kinetics. A single exponential with a time constant of 1 min was enough to describe the observed amplitude decay as a function of the data collection time.

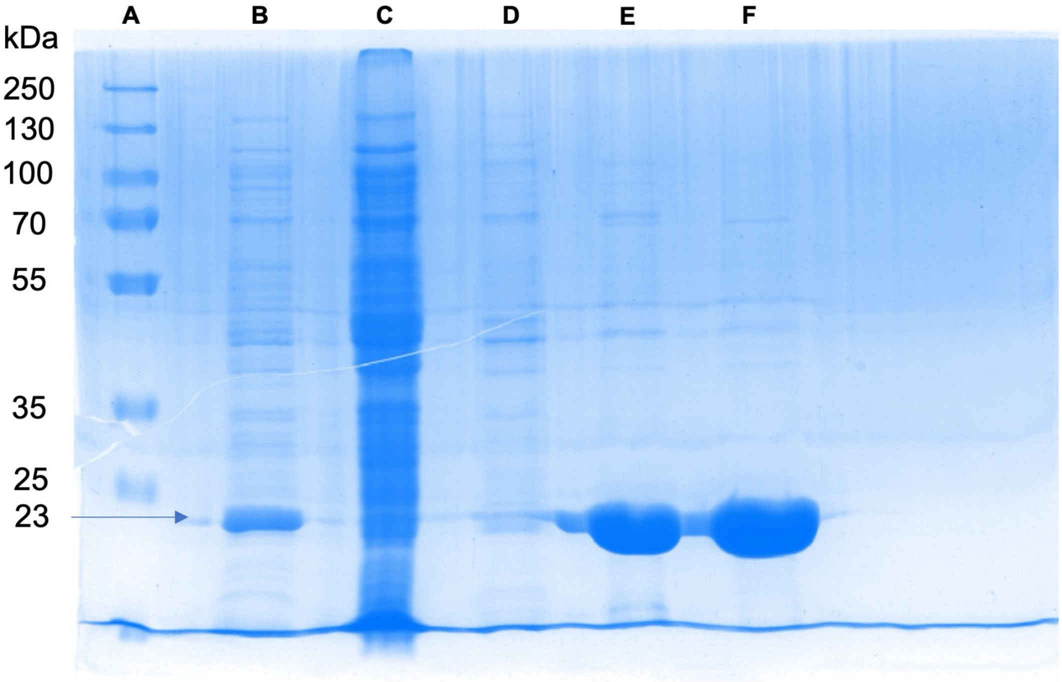


**Supplementary Fig. 31. Purification of *Tt*CBD assessed by SDS-PAGE**

SDS polyacrylamide gel electrophoresis (PAGE) was used to track *Tt*CBD during the purification steps and assess its purity. A: molecular weight markers; B: supernatant after centrifugation; C: flow-through of the Ni-NTA purification step; D: flow-through washing buffer; E: elution of the protein from the Ni-NTA column; F: protein after the SEC purification step.

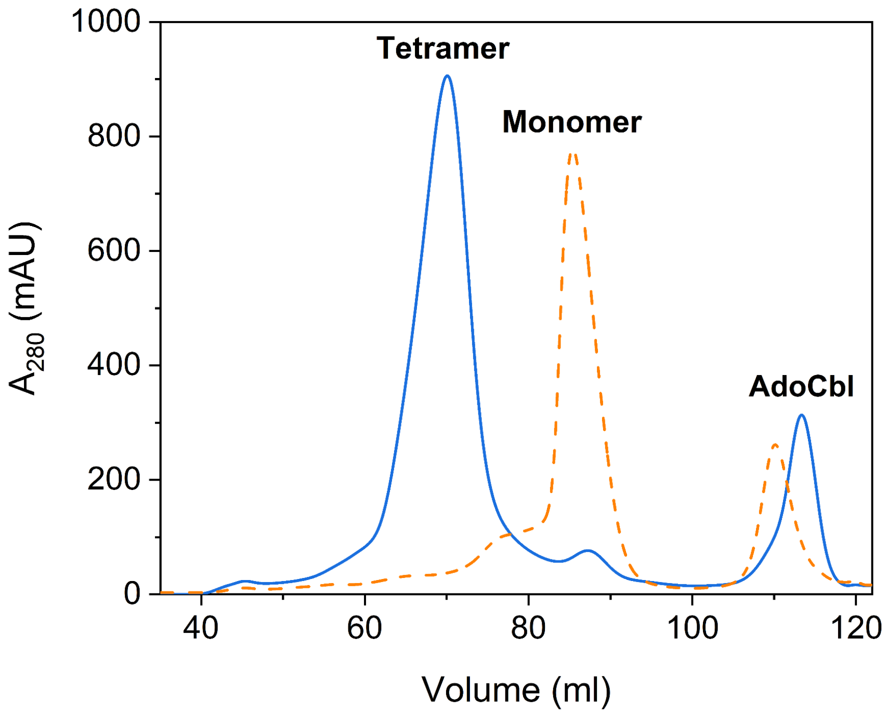


**<u>Supplementary Fig. 32. Size exclusion chromatogram of *Tt*CBD.</u>**

The SEC trace (from Superdex 200 Hiload 16/600 prep grade column) in blue is indicative of *Tt*CDB in the dark state, with the main elution peak at 70 ml corresponding to the tetrameric state of the protein. Residual traces of monomeric *Tt*CBD and excess of AdoCbl are found at higher elution volumes, ~ 87 ml and 114 ml respectively. The dashed orange line shows *Tt*CBD in the light-adapted state, i.e. monomeric. The light-adapted state was obtained upon illumination of the sample after the Ni-NTA purification.

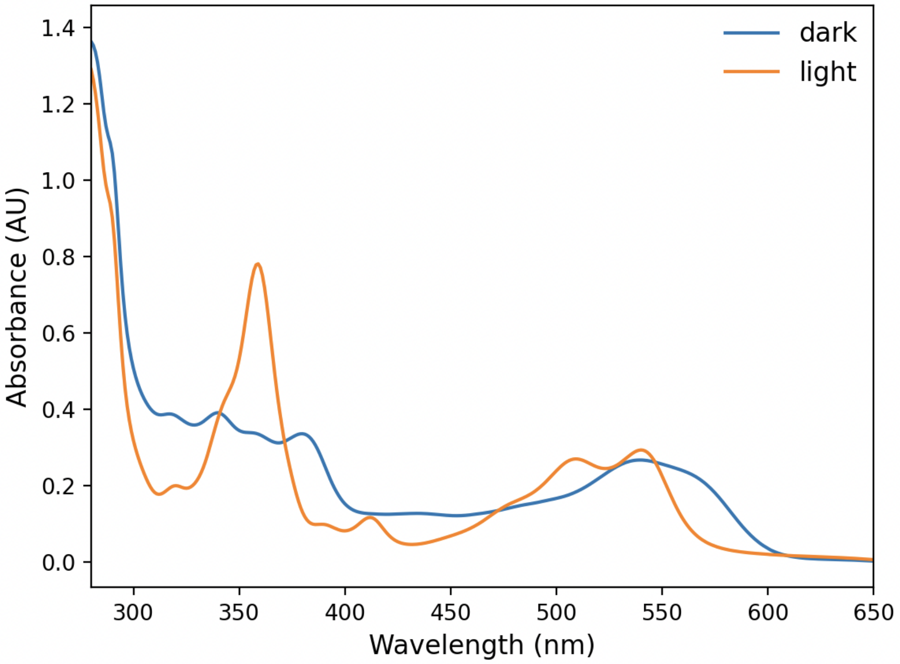


**<u>Supplementary Fig. 33. UV-vis absorption spectroscopy of *Tt*CBD in solution</u>**

UV-vis absorption spectra of *Tt*CBD in solution were collected with a JASCO V-570 spectrophotometer at 25 °C in the dark state (blue line) and after 2 min of illumination (orange line) with a 530-nm LED source at ~100 mW/cm² at the sample position.

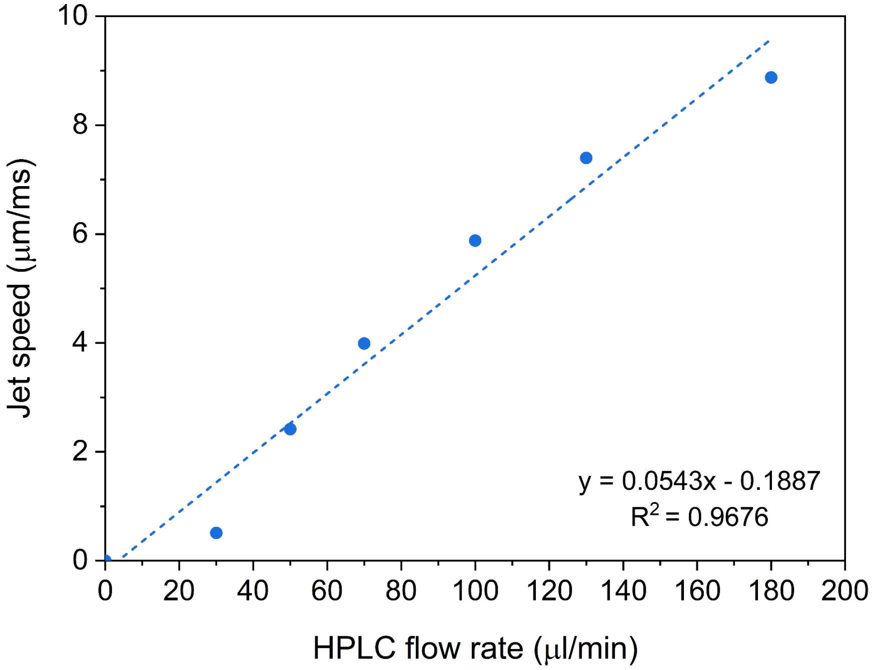


**Supplementary Fig. 34. Jet speed calibration as a function of the HPLC flowrate setting.**

*Tt*CBD crystals were embedded in CMC-5%. Calibration was performed using a HVE injector[8] with a polyamide-coated quartz capillary (inner diameter of 75 μm). Jet speed was monitored with a fast camera (Photron Mini) coupled to a zoom lens system (Navitar 12x).

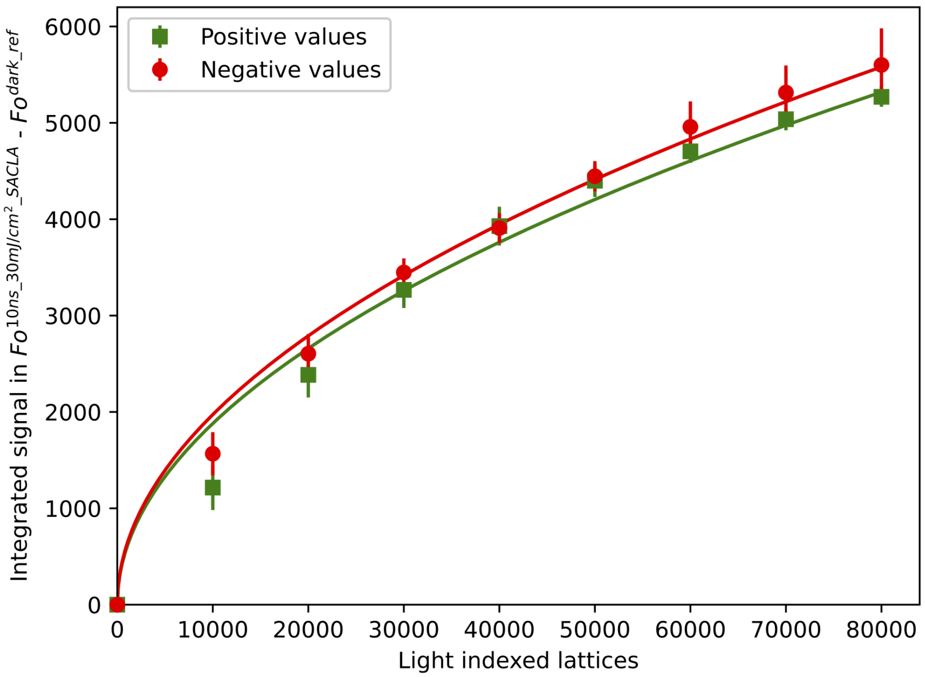


**Supplementary Fig. 35. Dependency of integrated Fourier difference peaks as a function of indexed lattices.**

Data sets based on an increasing number of *light* lattices of the 10 ns time delay (*10ns_30mJ/cm$^2$_SACLA*) were generated, ranging from 10,000 to 80,000 indexed lattices (from a total 86,000). A random selection was applied to compose these data sets, and the signal deviation was estimated by generating 25 different data sets. Fourier difference maps were computed using these data sets and the absolute sum of both positive (green squares) and negative (red circle) peaks within 2 Å of the AdoCbl were calculated. Red and green lines are resulting fits of absolute negative and positive values, respectively, using the following square-root function: $f(x) = a \times \sqrt{x}$, with $a$ the unique parameter used in the fit.

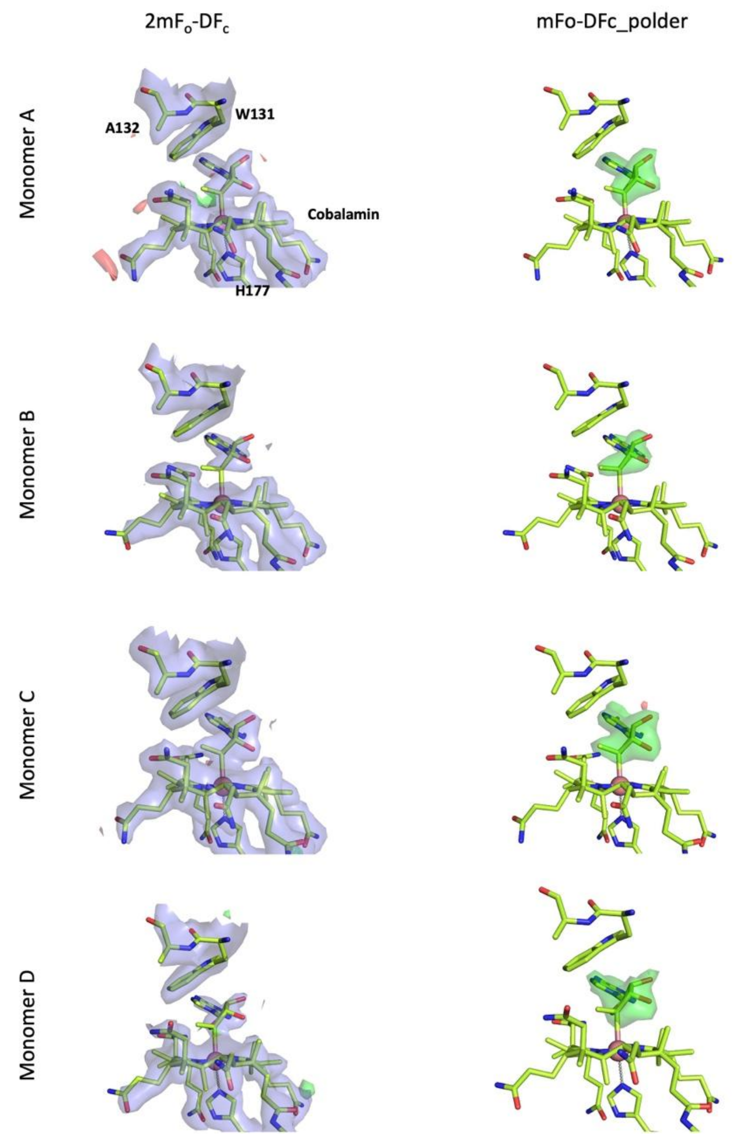


**<u>Supplementary Fig. 36. *illuminated_DLS* intermediate-state model of *Tt*CBD-H132A and 2mF$_o$-DF$_c$ electron density and mF$_o$-DF$_c$ polder maps (adenosyl moiety omitted)</u>**

Close-up view of the electron density maps (left, 2mF$_o$-DF and mF$_o$-DF$_c$) in the chromophore region and mF$_o$-DF$_c$_polder map omitting the adenosyl group for each monomer (right). As described on the MM section, the refined structure has two alternate conformations, one with the dark-state (Alt B) and the other of the intermediate state (Alt A). In this figure we omitted the dark-state (Alt B) and we only show the intermediate state (Alt A). The SSX intermediate state model is represented in lemon green sticks. 2mF$_o$-DF$_c$ is contoured at +1 σ, and mF$_o$-DF$_c$ at +3.5 σ (green) and -3.5 σ (red). mF$_o$-DF$_c$_polder map is shown at +3.5 σ (green) and -3.5 σ (red).

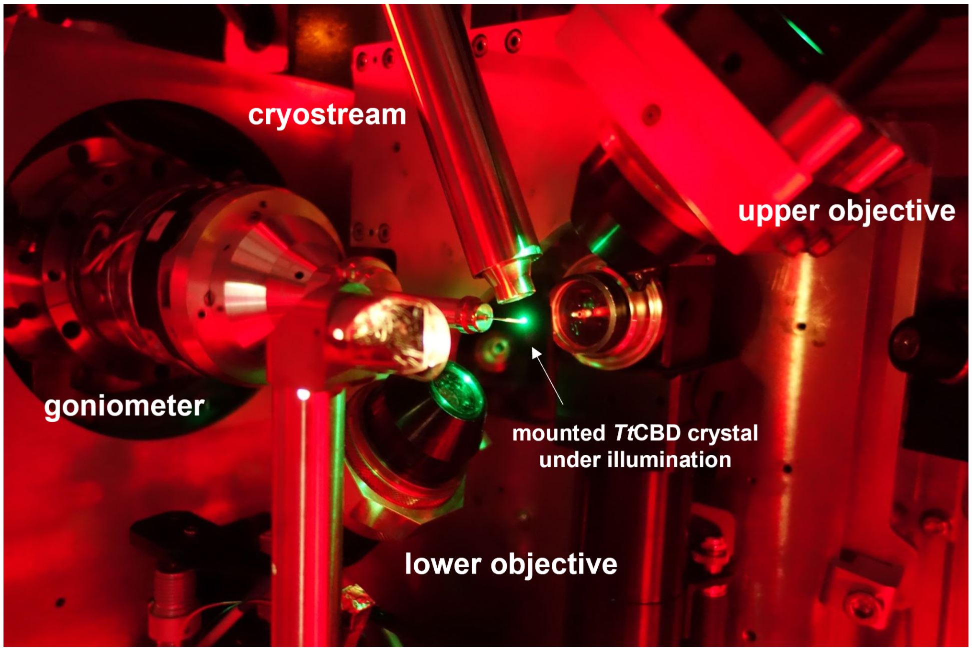


**<u>Supplementary Fig. 37. Online UV-vis microspectrophotometer mounted on the diffractometer of the BM07-FIP2 beamline of the ESRF</u>**

Illumination of a mounted *Tt*CBD crystal at the BM07-FIP2 beamline of the ESRF with the shown set-up: the microspectrophotometer is connected to the lower 4× demagnifying objective with a 910 µm (ID) optical fibre and a deuterium light source (DH2000 BAL light source) connected to the upper 4× demagnifying objective with a 400 µm (ID) optical fibre.

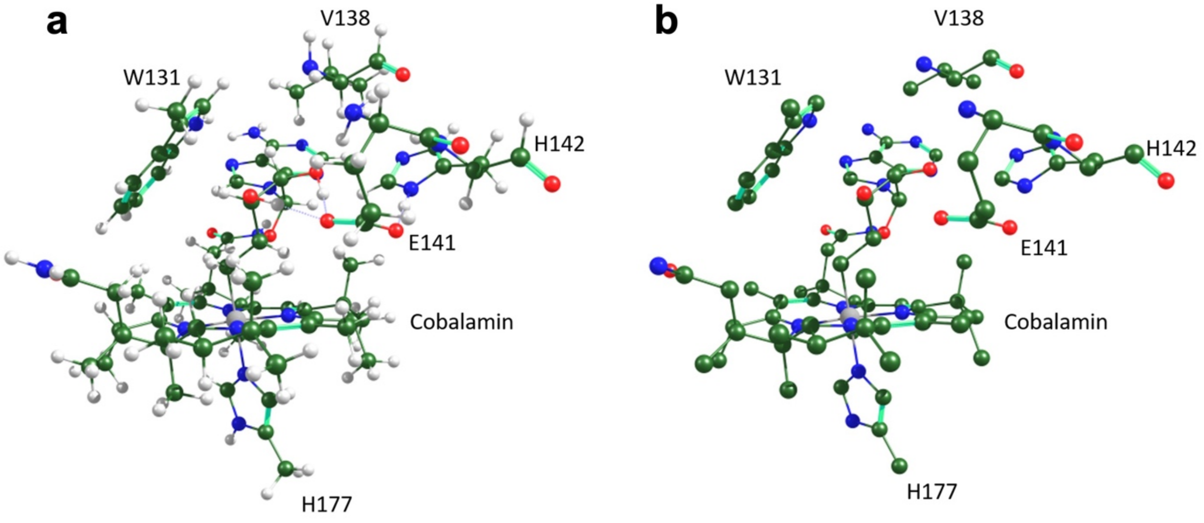


**Supplementary Fig. 38. Cluster model.**

Energy minimized structure of C5′-adduct (**5B** in Supplementary Fig. 23), (**a**) with and (**b**) without hydrogens for clarity.

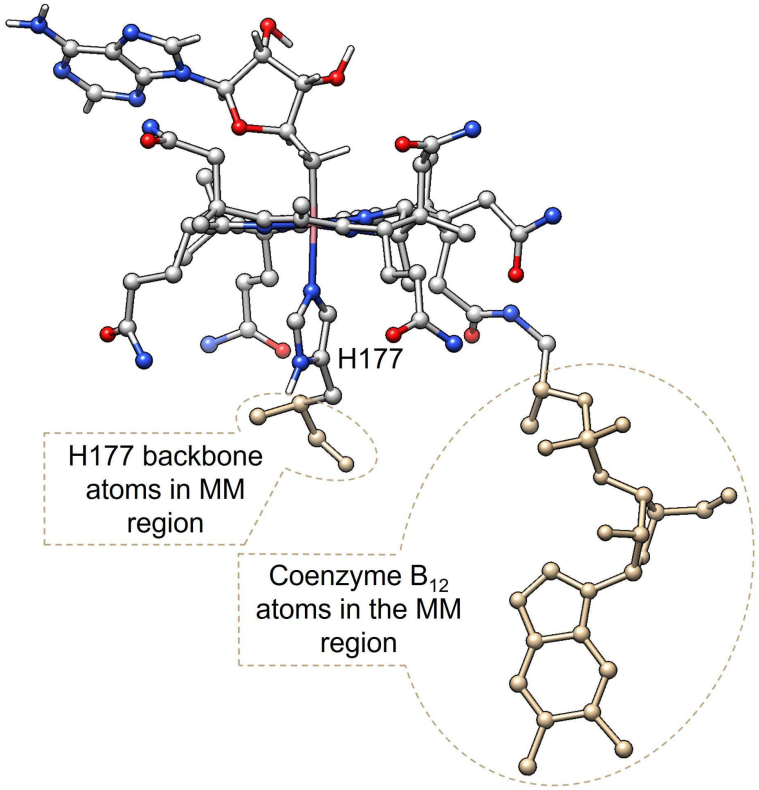


**Supplementary Fig. 39. QM/MM Partitioning.**
The partitioning of the high and low layers for the QM/MM simulations. The atoms of the QM region are shown in grey, red, blue, and pink for carbon, oxygen, nitrogen, and the cobalt ion, respectively. The atoms associated with the MM regions are shown in tan. For clarity of presentation the hydrogens were hidden from the corrin ring and the nucleotide loop. The QM region included the corrin ring, cobalt ion, the corrin ring side chains, and a portion of the nucleotide loop as well as the imidazole portion of the lower axial ligand H177 was protonated at the δ position. The backbone atoms of the H177 amino acid were considered in the MM region.

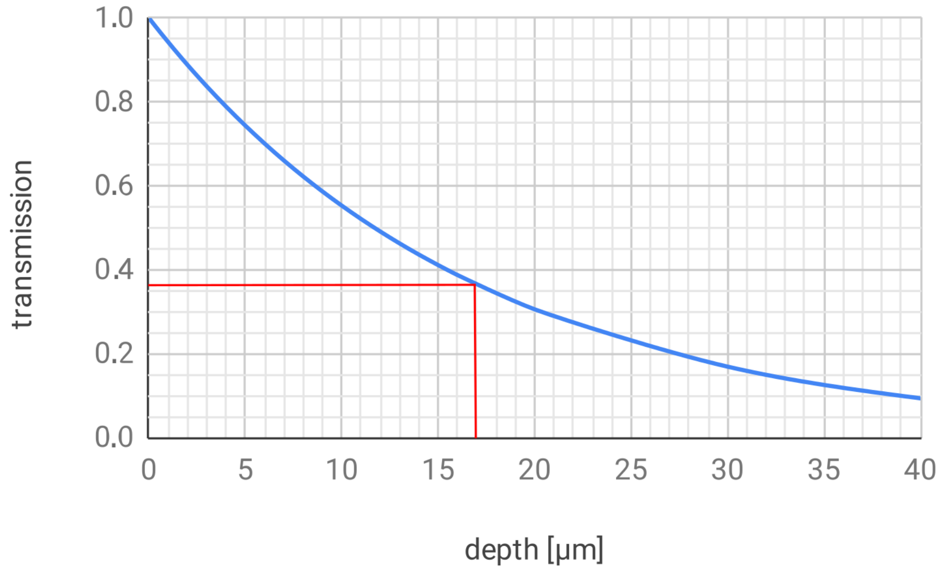


**Supplementary Fig. 40. Penetration of 530 nm light into *Tt*CBD microcrystals.**

Green light (530 nm) transmission through *Tt*CBD microcrystals. More than 70% of the light is transmitted through the average size (5 µm) of the microcrystals used in SFX experiments and the 1/e transmission depth is ~17 µm (indicated by the vertical red line).

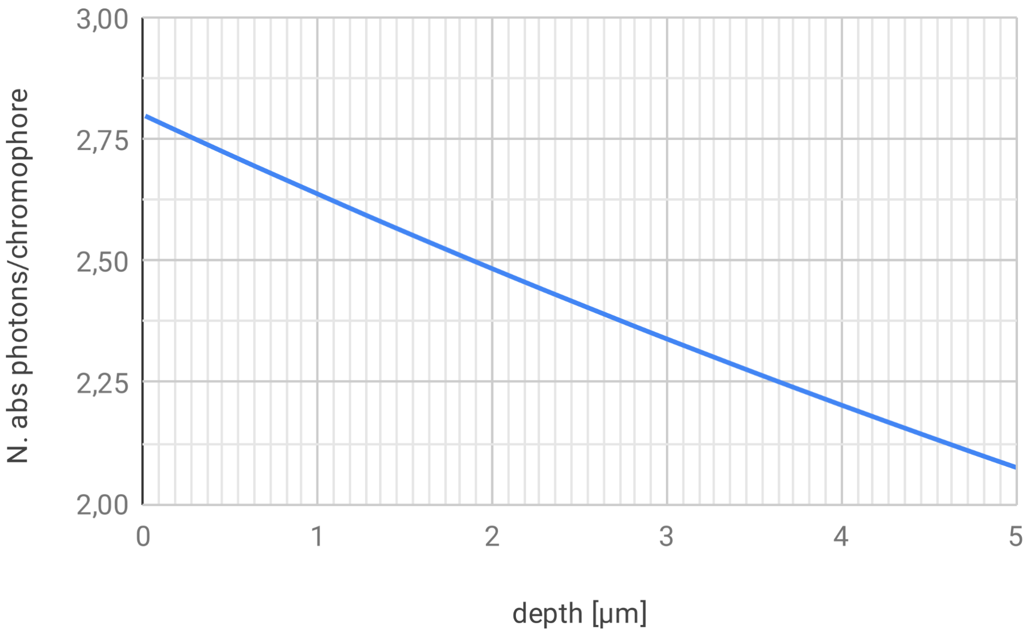


**<u>Supplementary Fig. 41. Absorbed photons *vs.* penetration depth in a *Tt*CBD crystal</u>**

Number of absorbed photons from a *Tt*CBD microcrystal at 530 nm as a function of the penetration depth with a laser fluence of 30 mJ/cm². The number of absorbed photons has been calculated using the Beer-Lambert law with a chromophore concentration of 28.6 mM (based on the TtCBD microcrystal unit cell parameters) and an extinction coefficient of 9010 $M^{-1}cm^{-1}$ at 530 nm Assuming an average microcrystal size of 5 µm, the average number of absorbed photons is 2.4.

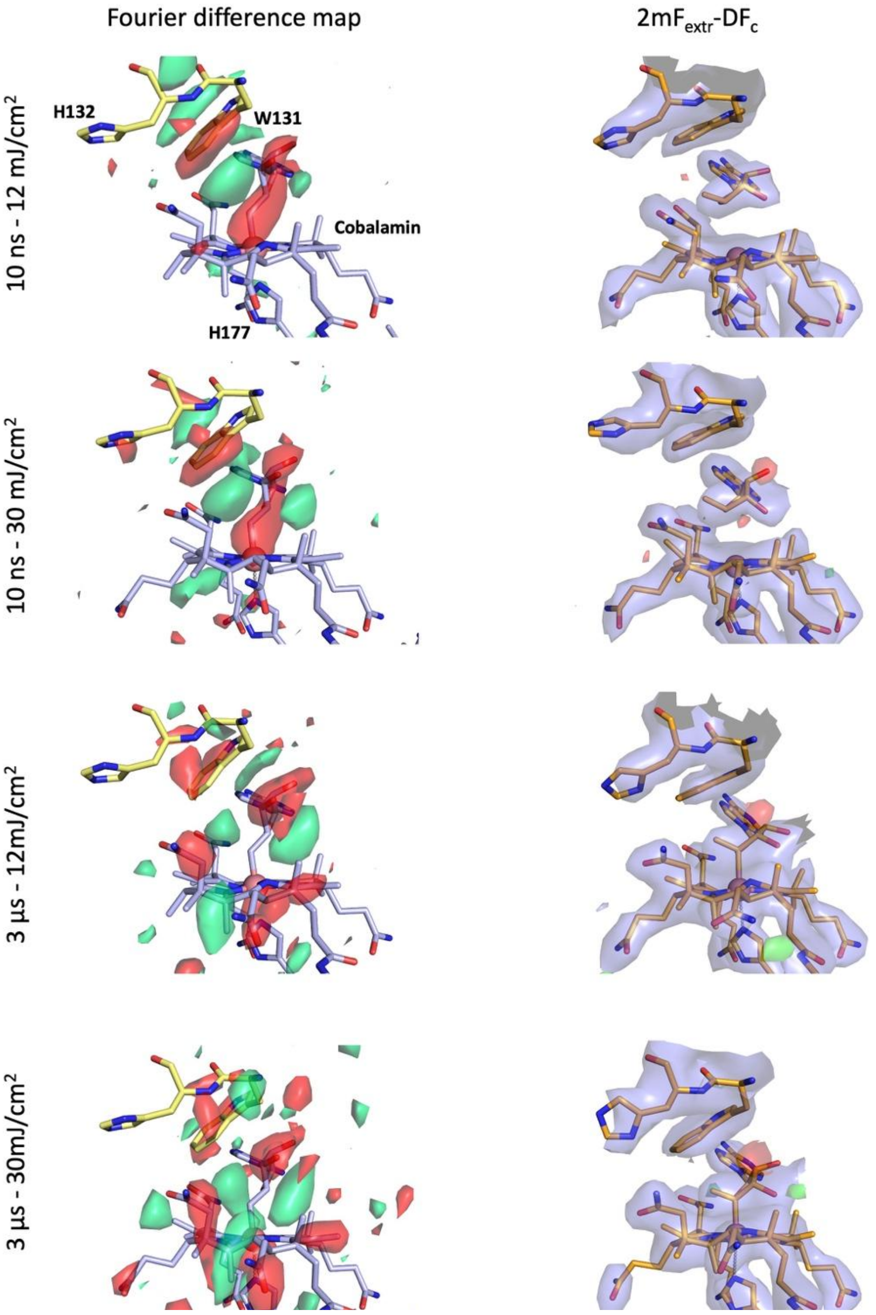


**Supplementary Fig. 42. Comparison of the Fourier difference map and the intermediate-state models at 10 ns and 3 µs time delays and at pump-laser fluences of 12 and 30 mJ/cm$^2$ (SACLA).**

Pump-laser fluences of 12 and 30 mJ/cm$^2$ correspond to 1 and 2.4 nominal absorbed photons per chromophore, respectively. Close-up view of the chromophore region in monomer B. Fourier difference maps $F_o^{\Delta t\_fluence\_SACLA}$ - $F_o^{dark\_ref\_SACLA}$ (**Δ**t: 10 ns and 3 µs; fluence: 12 and 30 mJ/cm$^2$) are contoured at +3.5 σ (green) and -3.5 σ (red) with the *dark_ref_SACLA* model represented in yellow (H132 and W131 of the four-helix bundle) and blue sticks (AdoCbl and H177 of the Rossmann fold) (left). Extrapolated electron density map 2m$F_{extr}$-D$F_c$ (contoured at +1 σ) and

$mF_{extr}$-$DF_c$ at +3.5 σ (green) and -3.5 σ (red) are overlaid with the intermediate-state models represented in orange sticks (right).

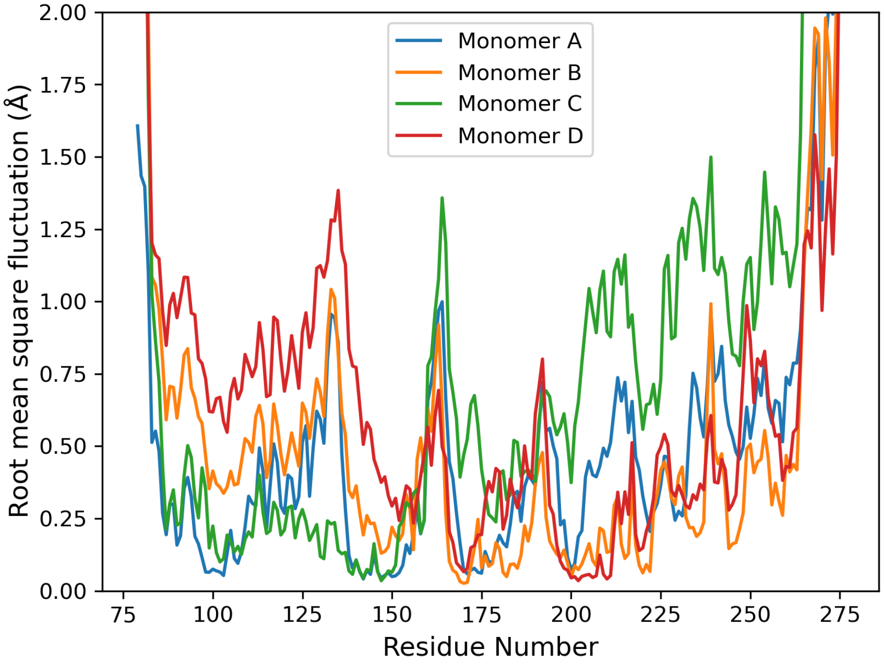


**Supplementary Fig. 43. Root mean square fluctuation (RMSF) obtained from the ensemble refinement of the *dark_ref_SACLA* data set.**

Ensemble refinement performed with *PHENIX* using the *dark_ref_SACLA* data set yielded 100 models, which were used to compute RMSF values of Cα atoms for the four different monomers. RMSF values of monomer A are in blue, monomer B in orange, monomer C in green and monomer D in red.

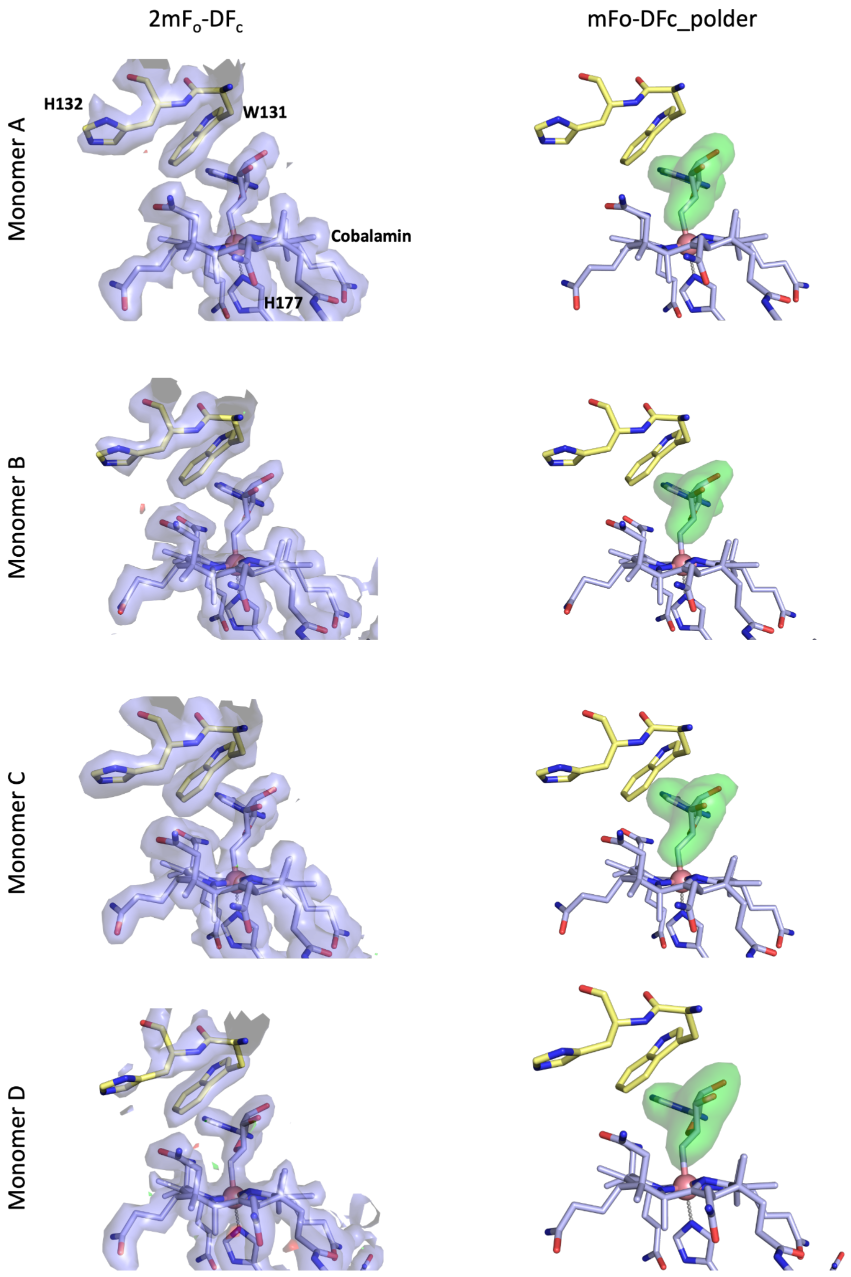


**Supplementary Fig. 44. *Dark_ref_SACLA* model and $2mF_o$-$DF_c$ electron density and $mF_o$-$DF_c$ polder maps (omitted adenosyl moiety).**

Close-up view in the chromophore region of the $2mF_o$-$DF_c$ (contoured at 1 σ) and $mF_o$-$DF_c$ (contoured at + 3.5 σ (green) and -3.5 σ (red)) electron density maps (left) and mFo-DFc_polder_map omitting the adenosyl group (right, contoured at +3.5 σ and coloured in green) for each monomer. The *dark_ref_SACLA* model is represented as sticks in yellow (H132 and W131 of the four-helix bundle) and blue (AdoCbl and H177 of the Rossmann fold).

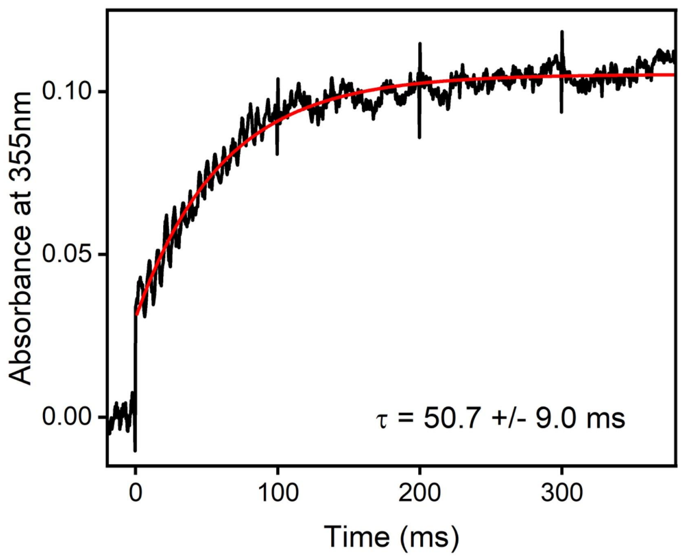


**Supplementary Fig. 45. Time-resolved absorption spectroscopy on the ms time scale in *Tt*CBD H132A mutant in solution after excitation at 530 nm.**

Kinetic transient at 355 nm on the ms timescale of samples containing 50 µM *Tt*CBD H132A mutant. Data were fitted to a single exponential equation (red line) to obtain the time constant.

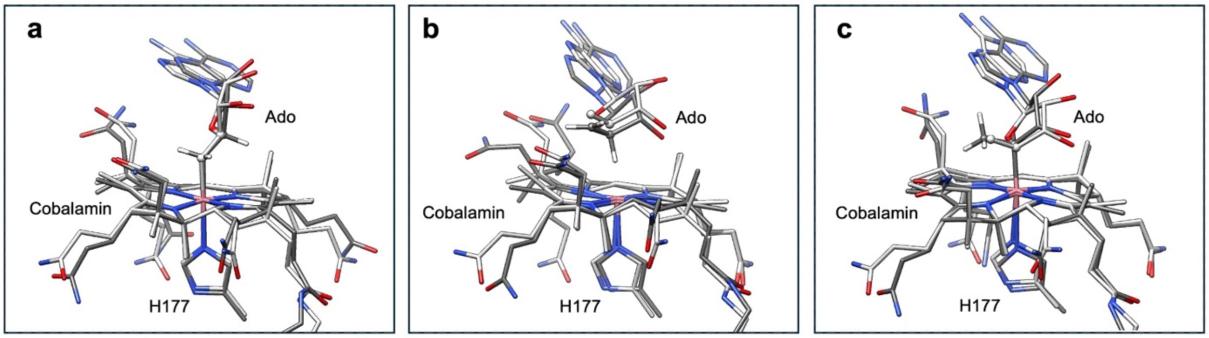


**Supplementary Fig. 46. Comparison of QM/MM optimized models with TR-SFX structures**
AdoCbl moiety with lower axial histidine (H177) with the QM/MM optimized models (dark grey) superimposed on the TR-SFX structures (light grey): (**a**) dark state (*dark_ref_SACLA*), (**b**) 10 ns time delay (*10ns_30mJ/cm$^2$_SACLA)* and (**c**) 3 µs time delay (*3µs_30mJ/cm$^2$_SACLA*). The hydrogens, with the exception of those bound to the C5′ and C4′ of the ribose ring, are hidden for clarity.

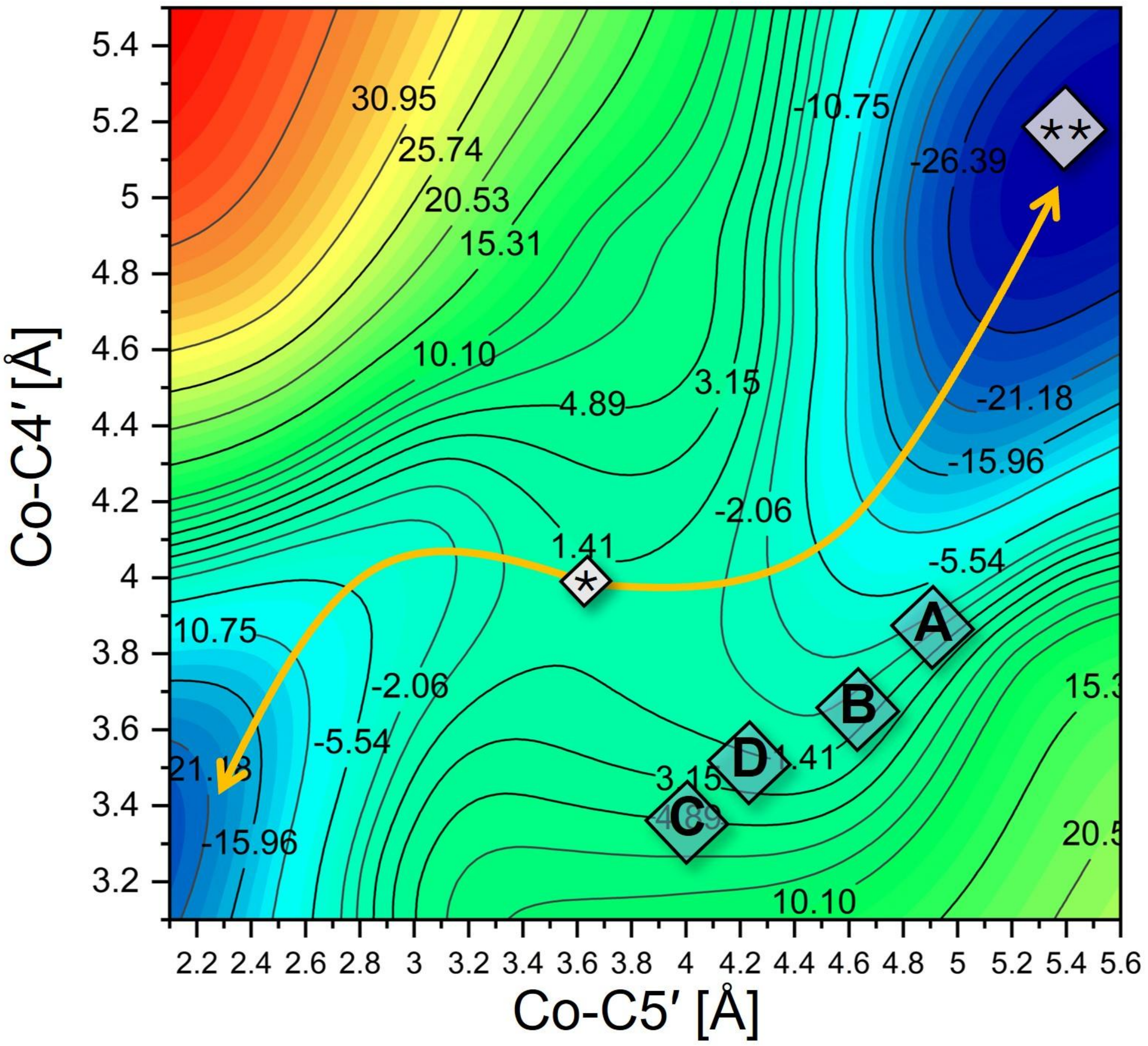


**Supplementary Fig. 47. CarH Diradical Potential Energy Surface.**

The approximate location of each of the monomers (diamonds labelled A - D) of the 10 ns crystal structure (10ns_30mJ/cm$^2$_SACLA) on the Co–C5′/Co–C4′ potential energy surface adopted from[81]. The arrow pointing towards the bottom left corresponds to the return to the AdoCbl CarH dark-state. The arrow pointing towards the top right corresponds to the region where hydridocobalamin is formed with 4′,5′-anhydroadenosine. The diamond containing the asterisk (*) corresponds to the lowest energy QM/MM optimized geometry for CarH where the diradical species involved the Co(II)/C5′• radical pair. The diamond containing two asterisks (**) corresponds to the stable local minimum with the photoproducts 4′,5′-anhydroadenosine and hydridocobalamin.

### Supplementary Tables

#### Supplementary Table 1. AdoCbl geometric parameters at 10 ns.

Selected geometric parameters for AdoCbl from the 10 ns crystal structure (*10ns_30mJ/cm²_SACLA*), the 10ns-diradical-model and the 10ns-photoproduct-model.

| 10 ns crystal structure (10ns_30mJ/cm$^2$_SACLA) | | | | | QM/MM | |
|---|---|---|---|---|---|---|
| | Mono A | Mono B | Mono C | Mono D | 10ns-diradical-model | 10-ns-photoproduct model |
| Bond Lengths/Distances (Å) | | | | | | |
| Co–C5' | 4.71 | 4.43 | 4.05 | 4.10 | 3.89 | 4.10 |
| Co–C4' | 3.62 | 3.54 | 3.29 | 3.38 | 3.08 | 4.48 |
| Co–$H_{C4'}$ | N/A | N/A | N/A | N/A | 1.92 | 1.45 |
| Co–$N_{H177}$ | 2.25 | 2.37 | 2.26 | 2.27 | 2.14 | 2.21 |
| C5'–C4' | 1.51 | 1.51 | 1.51 | 1.52 | 1.44 | 1.34 |
| Dihedrals (°) | | | | | | |
| $\theta_0$ C1'–C2'–C3'–C4' | -37.56 | -37.08 | -37.18 | -21.89 | -41.22 | -40.27 |
| $\theta_1$ C2'–C3'–C4'–O1' | 22.89 | 19.40 | 19.97 | -6.36 | 25.90 | 27.17 |
| $\theta_2$ C3'–C4'–O1'–C1' | 3.08 | 8.62 | 6.85 | 35.65 | 1.83 | -0.97 |
| $\theta_3$ C4'–O1'–C1'–C2' | -28.68 | -33.80 | -31.45 | -50.59 | -29.28 | -25.69 |
| $\theta_4$ O1'–C1'–C2'–C3' | 42.13 | 44.93 | 42.39 | 44.35 | 44.80 | 41.08 |
| Ribose Conformation | | | | | | |
| | 2′-endo | 2′-endo | 2′-endo | 2′-endo | 2′-endo | 2′-endo |

**Supplementary Table 2. AdoCbl geometric parameters at 3 µs.**

Selected geometric parameters for the AdoCbl in each monomer of the 3 µs crystal structure (3µs_30mJ/cm$^2$_SACLA) compared with the four QM/MM optimized models including the photoproducts hydridocobalamin and 4′,5′-anhydroadenosine (HCbl + anhAdo), a primary radical (Co(II)/C5′• RP), a tertiary diradical (Co(II)/C4′• RP), and finally a stable adduct where a bond was formed between the cobalt ion (Co(III)) and the C4 ′.

| 3 µs crystal structure (3µs_30mJ/cm$^2$_SACLA) | | | | | QM/MM | | | |
|---|---|---|---|---|---|---|---|---|
| | Mono A | Mono B | Mono C | Mono D | HCbl + anhAdo | Co(II)/C5′• RP | Co(II)/C4′• RP | Adduct |
| Bond Lengths/Distances (Å) | | | | | | | | |
| Co–C5' | 3.57 | 3.25 | 3.11 | 3.05 | 4.18 | 4.01 | 3.92 | 2.11 |
| Co–C4' | 2.44 | 2.26 | 2.20 | 2.18 | 4.24 | 3.25 | 4.10 | 2.92 |
| Co–$H_{C4'}$ | N/A | N/A | N/A | N/A | 1.45 | 2.17 | N/A | N/A |
| Co–$N_{H177}$ | 2.29 | 2.64 | 2.56 | 2.51 | 2.15 | 2.19 | 2.20 | 2.92 |
| C5'–C4' | 1.52 | 1.51 | 1.52 | 1.50 | 1.34 | 1.47 | 1.49 | 1.52 |
| Dihedrals (°) | | | | | | | | |
| $\theta_0$ C1'–C2'–C3'–C4' | -20.17 | -25.96 | -35.81 | -29.48 | -39.91 | -39.67 | -24.05 | -39.80 |
| $\theta_1$ C2'–C3'–C4'–O1' | 20.56 | 20.64 | 24.98 | 19.81 | 29.35 | 37.47 | 11.99 | 30.38 |
| $\theta_2$ C3'–C4'–O1'–C1' | -11.98 | -6.14 | -3.04 | -1.09 | -5.07 | -20.67 | 6.26 | -8.68 |
| $\theta_3$ C4'–O1'–C1'–C2' | -1.14 | -10.87 | -20.38 | -18.17 | -21.37 | -5.46 | -21.75 | -17.54 |
| $\theta_4$ O1'–C1'–C2'–C3' | 14.01 | 23.36 | 35.55 | 29.78 | 38.44 | 29.41 | 28.16 | 35.91 |
| Ribose Conformation | | | | | | | | |
| | 2′-endo | 2′-endo | 2′-endo | 2′-endo | 2′-endo | 2′-endo | 2′-endo | 2′-endo |

**Supplementary Table 3. Ground state potential energies relative to the photoproduct (HCbl + anhAdo) and selected structural parameters for the cluster model.**

| | *ΔE* (kcal mol)$^{-1}$ | R(Co–C4′) (Å) | R(Co–C5′) (Å) | R(CX′–H)[1] (Å) | R(Co–H) (Å) | R(Co–N(His)) (Å) |
|---|---|---|---|---|---|---|
| HCbl + anhAdo (**1**) | 0 | 3.83 | 3.80 | - | 1.43 | 2.12 |
| [Co···H···C5′]$^{‡}$ (**2A**) | 2.08 | 3.68 | 3.02 | 1.50 (C5) | 1.55 | 2.42 |
| Co···H···C5′ intermediate (**3A**) | 0.41 | 3.68 | 2.92 | 1.28 (C5) | 1.66 | 2.43 |
| [Co(II) + C4′·]$^{‡}$ (**4A**) | 7.30 | 3.56 | 3.56 | 1.10 (C5) | 2.73 | 2.09 |
| Co–C4′ adduct (**5A**) | -11.51 | 2.17 | 3.00 | 1.10 (C5) | 3.09 | 2.43 |
| Co–C4′ adduct (b)[2] | -12.58 | 2.09 | 2.97 | 1.10 (C5) | 3.09 | 2.14 |
| [Co···H···C4′]$^{‡}$ (**2B**) | 14.24 | 3.05 | 3.73 | 1.41 (C4) | 1.69 | 2.35 |
| Cob(II) + C5′· (a) (**3B**) | 7.81 | 3.13 | 3.95 | 1.14 (C4) | 2.06 | 2.08 |
| Cob(II) + C5′· (b) (**4B**) | 10.43 | 3.40 | 3.65 | 1.11 (C4) | 2.70 | 2.09 |
| Co–C5′ adduct (**5B**) | -24.28 | 3.03 | 2.01 | 1.11 (C4) | 3.11 | 2.09 |
| Co–C5′ adduct (b)[2] | -21.03 | 3.04 | 2.05 | 1.11 (C4) | 3.11 | 2.51 |

[1] CX′ = C5′ or C4′ when H is bonded to C5′ or C4′, respectively

[2] The second energy minimized Co–C4′ and Co–C5′ adducts not presented in the potential energy profile

#### Supplementary Table 4. Data collection and refinement statistics of TR-SFX power titration at SACLA

| | 3μs_12mJ/cm$^2$_SACLA | 3μs_30mJ/cm$^2$_SACLA | 3μs_60mJ/cm$^2$_SACLA | 3μs_120mJ/cm$^2$_SACLA |
|---|---|---|---|---|
| | | Power - Titration | | |
| Matrix | CMC 5% | CMC 5% | CMC 5% | CMC 5% |
| PDB Code | 9S0D | N/A | N/A | N/A |
| Facility (Beamline) | | SACLA (BL2-EH3 ) | | |
| Dark/Light | light | light | light | light |
| Pump laser repetition rate (Hz) | 10 | 10 | 30 | 10 |
| Space group | | $P_{21\ 21\ 21}$ | | |
| Unit-cell parameters¶ | | | | |
| a (Å) | 64.76 (0.12) | 64.77 (0.11) | 64.78 (0.11) | 64.77 (0.12) |
| b (Å) | 71.14 (0.16) | 71.09 (0.14) | 71.15 (0.13) | 71.13 (0.14) |
| c (Å) | 207.29 (0.42) | 207.33 (0.42) | 207.36 (0.46) | 207.37 (0.43) |
| *a (°)* | 90.0 (0.2) | 90.0 (0.2) | 90.0 (0.2) | 90.0 (0.2) |
| *β (°)* | 90.0 (0.2) | 90.0 (0.2) | 90.0 (0.2) | 90.03 (0.2) |
| *γ (°)* | 90.0 (0.1) | 90.0 (0.1) | 90.0 (0.1) | 90.01 (0.2) |
| Resolution (Å)# | 50 - 2.25 (2.30 - 2.25) | 50 - 2.25 (2.30 - 2.25) | 50 - 2.25 (2.30 - 2.25) | 50 - 2.25 (2.30 - 2.25) |
| I / σI | 8.42 (0.92) | 8.58 (1.01) | 8.41 (0.95) | 8.63 (0.99) |
| Rsplit (%) | 9.06 (116.81) | 8.96 (108.88) | 9.21 (114.39) | 9.02 (110.32) |
| CC* (%) | 99.89 (81.37) | 99.87 (80.33) | 99.87 (81.94) | 99.87 (81.55) |
| CC 1/2 (%) | 99.53 (49.49) | 99.50 (47.64) | 99.48 (50.55) | 99.51 (49.82) |
| Wilson B-factor (Å$^2$) | 51.22 | 48.86 | 50.57 | 51.01 |
| Completeness (%) | 99.98 (100) | 99.98 (100) | 99.98 (100) | 99.98 (100) |
| # collected images | 131,974 | 133,083 | 146,575 | 136,957 |
| # hits | 47,238 | 52,475 | 56,074 | 48,293 |
| Hit rate (%) | 35.80 | 39.43 | 38.25 | 35.26 |
| # indexed lattices | 66,053 | 66,053 | 66,053 | 66,053 |
| Success rate* (%) | 50.0 | 49.6 | 45.1 | 48.2 |
| # total reflections | 47,726,286 (2,212,796 ) | 48,277,155(2,250,091) | 45,850,784 (2,118,676) | 47802506 (2214728) |
| # unique reflections | 43,537 (2,834 ) | 44,674 (2,930) | 43,563 (2,832) | 43,558 (2,835) |
| Multiplicity | 1,096 (781) | 1,081 (768) | 1,052.51(748) | 1,098 (781) |
| Refinement§ | | | | |
| Rwork (%) | 27.78 (46.77) | | | |
| Rfree (%) | 31.84 (47.12) | | | |
| No. Atoms | 6,794 | | | |
| Protein | 6,084 | | | |

| | |
|---|---|
| Ligands | 436 |
| Solvent | 274 |
| Average B-factor ($Å^2$) | 54.43 |
| Protein | 54.51 |
| Ligands | 53.62 |
| Solvent | 54.11 |
| R.m.s. deviations | |
| Bond lengths (Å) | 0.007 |
| Bond angles (°) | 1.02 |
| Ramachandran | |
| Favoured (%) | 97.6 |
| Allowed (%) | 2.4 |
| Outliers (%) | 0.0 |
| Rotamer outliers (%) | 3.9 |
| Clashscore | 8.5 |

* Success rate: number of indexed lattices/total number of recorded images.

¶ Values in parenthesis correspond to the standard deviation of the fitted normal distribution.

### Values in parenthesis correspond to the highest resolution shell.

§ For the light data sets, refinement statistics are obtained from models refined against extrapolated structure factor amplitudes.

#### Supplementary Table 5. Data collection and refinement statistics of TR-SFX time series at SACLA

| | dark_only_SACLA | dark_ref_SACLA | 10ns_30mJ/cm$^2$_ SACLA | 300ns_30mJ/cm$^2$_ SACLA | 3µs_30mJ/cm$^2$_ SACLA | 100µs_30mJ/cm$^2$_ SACLA | 3ms_30mJ/cm$^2$_ SACLA | 10ns_12mJ/cm$^2$_ SACLA |
|---|---|---|---|---|---|---|---|---|
| Matrix | CMC-5% | CMC-5% | CMC-5% | CMC-5% | CMC-5% | CMC-5% | CMC-5% | CMC-5% |
| PDB Code | N/A | 9S06 | 9S08 | 9S09 | 9S0E | 9S0A | 9S0B | 9S0C |
| Facility (Beamline) | | | | SACLA (BL2-EH3 ) | | | | |
| Dark/Light | dark | dark | light | light | light | light | light | light |
| Pump laser repetition rate (Hz) | 0 | N.A | 15 | 15 | 10 | 15 | 15 | 15 |
| Space group | | | | $P_{21\ 21\ 21}$ | | | | |
| Unit-cell parameters[¶] | | | | | | | | |
| a (Å) | 64.85 (0.13) | 64.80 (0.12) | 64.81 (0.11) | 64.80 (0.11) | 64.77(0.11) | 64.79 (0.12) | 64.74 (0.11) | 64.82 (0.11) |
| b (Å) | 71.11 (0.14) | 71.09 (0.14) | 71.06 (0.14) | 71.06 (0.13) | 71.09 (0.14) | 71.21 (0.16) | 71.28 (0.19) | 71.10 (0.13) |
| c (Å) | 207.37 (0.50) | 207.26 (0.46) | 207.45 (0.55) | 207.23 (0.47) | 207.33 (0.42) | 207.25 (0.47) | 207.21 (0.48) | 207.36 (0.43) |
| *a (°)* | 90.0 (0.2) | 90.0 (0.2) | 90.0 (0.2) | 90.0 (0.2) | 90.0 (0.2) | 90.0 (0.2) | 90.0 (0.2) | 90.0 (0.2) |
| *β (°)* | 90.0 (0.2) | 90.0 (0.2) | 90.0 (0.2) | 90.0 (0.2) | 90.0 (0.2) | 90.0 (0.2) | 90.0 (0.2) | 90.0 (0.2) |
| *ɣ (°)* | 90.0 (0.1) | 90.0 (0.1) | 90.0 (0.1) | 90.0 (0.1) | 90.0 (0.1) | 90.0 (0.1) | 90.0 (0.1) | 90.0 (0.1) |
| Resolution (Å)[#] | 50 - 2.2 (2.25 - 2.2) | 50 - 2.05 (2.10 - 2.05) | 50 - 2.30 (2.35 - 2.30) | 50 - 2.30 (2.35 - 2.30) | 50 - 2.25 (2.30 - 2.25) | 50 - 2.30 (2.35 - 2.30) | 50 - 2.35 (2.40 - 2.35) | 50 - 2.28 (2.33 - 2.28) |
| I / σI | 10.19 (1.04) | 22.83 (1.87) | 10.07 (1.02) | 9.40 (1.03) | 8.73 (0.94) | 8.84 (0.92) | 9.62 (1.06) | 9.25 (1.06) |
| Rsplit (%) | 7.50 (107.48) | 3.04 (59.02) | 7.82 (104.71) | 8.29 (105.62) | 8.63 (114.93) | 8.87 (116.84) | 8.16 (100.10) | 8.34 (102.33) |
| CC* (%) | 99.92 (81.97) | 99.98 (90.64) | 99.90 (83.91) | 99.90 (83.77) | 99.89 (79.23) | 99.88 (81.43) | 99.89 (86.97) | 99.89 (82.65) |
| $CC_{1/2}$ (%) | 99.70 (50.60) | 99.95 (69.73) | 99.63 (54.33) | 99.60 (54.06) | 99.56 (45.74) | 99.55 (49.60) | 99.59 (60.83) | 99.59 (51.87) |
| Wilson B-factor (Å$^2$) | 44.33 | 41.16 | 52.04 | 51.13 | 47.60 | 52.23 | 52.91 | 48.78 |
| Completeness (%) | 100.0 (100.0) | 100.0 (100.0) | 100.0 (100.0) | 100.0 (100.0) | 100.0 (100.0) | 100.0 (100.0) | 100.0 (100.0) | 100.0 (100.0) |
| # collected images | 114,111 | 1,709,013 | 220,172 | 241,824 | 133,083 | 196,693 | 162,035 | 136,607 |
| # hits | 83,999 | 634,317 | 78,689 | 63,544 | 52,475 | 54,455 | 53,502 | 56,291 |
| Hit rate (%) | 73.6 | 37.0 | 35.7 | 26.3 | 39.4 | 27.7 | 33.0 | 41.2 |
| # indexed lattices | 115,060 | 847,084 | 91,159 | 83,486 | 73,296 | 72,429 | 75,958 | 78,864 |

| | | | | | | | | |
|---|---|---|---|---|---|---|---|---|
| Success rate* (%) | 100.8 | 49.6 | 41.4 | 34.5 | 55.0 | 36.8 | 46.9 | 57.3 |
| # total reflections | 89,613,137 (4,178,048 ) | 788,407,590 (36,957,378) | 70,268,336 (3,258,927) | 59,839,102 (2,779,977) | 55,116,389 (2,581,031) | 53,269,867 (2,465,796) | 55,465,226 (2,612,858) | 57,190,963 (2,661,822) |
| # unique reflections | 49,718 (3,258 ) | 61,105 (4,014) | 43,560 (2,842) | 43,554 (2,846 ) | 46,453 (3,053) | 43,560 (2,836) | 40,934 (2,712) | 44,719 (2,928) |
| Multiplicity | 1802 (1,282) | 12903 (9,207) | 1613 (1,147) | 1374 (977) | 1,186 (845) | 1,220 (870) | 1,355 (963) | 1,279 (909) |
| Refinement§ | | | | | | | | |
| Rwork (%) | | 17.43 (42.47) | 25.20 (43.60) | 27.73 (43.89) | 27.08 (44.54) | 28.88 (42.95) | 26.76 (44.87) | 25.28 (46.91) |
| Rfree (%) | | 21.98 (40.34) | 30.84 (47.30) | 33.03 (46.62) | 31.59 (42.23) | 35.07 (43.13) | 31.51 (51.35) | 29.94 (50.21) |
| No. Atoms | | 6,915 | 6,925 | 6,667 | 6,822 | 6,736 | 6,748 | 6,773 |
| Protein | | 6,119 | 6,145 | 6,022 | 6,102 | 6,026 | 6,031 | 6,077 |
| Ligands | | 436 | 436 | 436 | 436 | 436 | 436 | 436 |
| Solvent | | 360 | 344 | 209 | 284 | 274 | 281 | 260 |
| Average B-factor (Å$^2$) | | 56.86 | 54.66 | 56.45 | 62.30 | 51.77 | 54.75 | 64.88 |
| Protein | | 57.05 | 54.88 | 56.80 | 62.57 | 51.87 | 54.41 | 65.17 |
| Ligands | | 51.72 | 51.09 | 53.25 | 58.52 | 50.78 | 58.66 | 61.81 |
| Solvent | | 59.87 | 55.30 | 53.03 | 62.13 | 51.36 | 55.92 | 63.23 |
| R.m.s. deviations | | | | | | | | |
| Bond lengths (Å) | | 0.008 | 0.007 | 0.007 | 0.008 | 0.011 | 0.005 | 0.009 |
| Bond angles (°) | | 1.03 | 0.97 | 1.00 | 1.11 | 1.83 | 0.78 | 1.19 |
| Ramachandran | | | | | | | | |
| Favoured (%) | | 98.0 | 97.4 | 97.7 | 96.3 | 96.8 | 97.8 | 97.3 |
| Allowed (%) | | 2.0 | 2.6 | 2.3 | 3.7 | 3.2 | 2.2 | 2.7 |
| Outliers (%) | | 0.0 | 0.0 | 0.0 | 0.0 | 0.0 | 0.0 | 0.0 |
| Rotamer outliers (%) | | 1.7 | 5.5 | 2.2 | 5.9 | 3.6 | 2.5 | 4.7 |
| Clashscore | | 4.5 | 6.9 | 8.1 | 10.4 | 8.9 | 7.0 | 7.7 |

* Success rate: number of indexed lattices/total number of recorded images.

¶ Values in parenthesis correspond to the standard deviation of the fitted normal distribution.

### Values in parenthesis correspond to the highest resolution shell.

§ For the light data sets, refinement statistics are obtained from models refined against extrapolated structure factor amplitudes.

#### Supplementary Table 6. SwissFEL TR-SFX Data collection and refinement statistics

| | dark_only_SwissFEL | dark_ref_SwissFEL | 3μs_30mJ/cm2_SwissFEL | 10ms_30mJ/cm2_SwissFEL |
|---|---|---|---|---|
| PDB Code | N/A | 9S0F | 9S07 | 9S0J |
| Facility (Beamline) | SwissFEL (Cristallina ) | | | |
| Dark/Light | dark | dark | light | light |
| Pump laser repetition rate (Hz) | 0 | N.A | 50 | 25 |
| Space group | $P_{21\ 21\ 21}$ | | | |
| Unit-cell parameters¶ | | | | |
| a (Å) | 65.52 (0.07) | 65.52 (0.07) | 65.46 (0.07) | 65.44 (0.13) |
| b (Å) | 72.12 (0.08) | 72.13 (0.10) | 72.15 (0.12) | 72.65 (0.30) |
| c (Å) | 209.49 (0.30) | 209.53 (0.35) | 209.56 (0.36) | 209.58 (0.67) |
| *α (°)* | 90.0 (0.1) | 90.0 (0.1) | 90.0 (0.1) | 90.0 (0.2) |
| *β (°)* | 90.0 (0.1) | 90.0 (0.1) | 90.0 (0.1) | 90.0 (0.2) |
| *γ (°)* | 90.0 (0.1) | 90.0 (0.1) | 90.0 (0.1) | 90.0 (0.1) |
| Resolution (Å)# | 50 - 2.5 (2.55 - 2.5) | 50 - 2.35 (2.40 - 2.35) | 50 - 2.60 (2.65 - 2.60) | 50 - 2.90 (2.95 - 2.90) |
| I / σI | 6.5 (1.2) | 8.8 (1.1) | 5.9 (0.8) | 7.8 (1.3) |
| Rsplit (%) | 13.1 (97.2) | 8.4 (98.8) | 13.6 (103.4) | 11.6 (79.5) |
| CC* (%) | 99.69 (79.64) | 99.89 (85.23) | 99.76 (83.80) | 99.76 (82.26) |
| $CC_{1/2}$ (%) | 98.35 (46.45) | 99.57 (57.04) | 98.65 (54.11) | 99.05 (51.14) |
| Wilson B-factor ($Å^2$) | 48.73 | 48.47 | 51.64 | 63.70 |
| Completeness (%) | 100.0 (100.0) | 100.0 (100.0) | 100.0 (100.0) | 100.0 (100.0) |
| # collected images | 131,350 | 368,219 | 65,740 | 275,733 |
| # hits | 107,353 | 262,921 | 44,725 | 175,343 |
| Hit rate (%) | 81.7 | 71.4 | 68.0 | 63.6 |
| # indexed lattices | 68,943 | 150,910 | 33,798 | 47,007 |
| Success rate* (%) | 52.5 | 41.0 | 51.4 | 17.0 |
| # total reflections | 33,321,437 (1,464,685) | 74,356,065 (3,017,709) | 15,774,309 (718,158) | 14,390,586 (659,478) |
| # unique reflections | 35,213 (2,297) | 42,285 (2,791) | 31,460 (2,056) | 22,989 (1,519) |
| Multiplicity | 946 (638) | 1,758 (1,074) | 501 (349) | 625 (434) |
| Refinement§ | | | | |
| Rwork (%) | | 16.11 (30.74) | 26.44 (43.10) | 27.03 (38.83) |
| Rfree (%) | | 21.17 (38.64) | 32.84 (46.74) | 33.27 (49.23) |
| No. Atoms | | 6,899 | 6,586 | 6,447 |
| Protein | | 6,110 | 6,013 | 6,019 |
| Ligands | | 436 | 436 | 382 |
| Solvent | | 353 | 137 | 46 |
| Average B-factor ($Å^2$) | | 58.81 | 66.10 | 73.07 |

| | | | |
|---|---|---|---|
| Protein | 59.07 | 65.91 | 72.28 |
| Ligands | 52.97 | 68.14 | 87.20 |
| Solvent | 61.62 | 68.09 | 59.01 |
| R.m.s. deviations | | | |
| Bond lengths (Å) | 0.009 | 0.013 | 0.004 |
| Bond angles (°) | 1.08 | 1.60 | 0.8 |
| Ramachandran | | | |
| Favoured (%) | 98.1 | 95.6 | 97.2 |
| Allowed (%) | 1.9 | 4.4 | 2.7 |
| Outliers (%) | 0.0 | 0.0 | 0.1 |
| Rotamer outliers (%) | 3.0 | 2.0 | 3.5 |
| Clashscore | 3.6 | 12.1 | 10.1 |

* Success rate: number of indexed lattices/total number of images.

¶ Values in parenthesis correspond to the standard deviation of the fitted normal distribution.

### Values in parenthesis correspond to the highest resolution shell.

§ For the light data sets, refinement statistics are obtained from models refined against extrapolated structure factor amplitudes

#### Supplementary Table 7. Diamond SSX Data collection and refinement statistics of *Tt*CBD mutant - H132A

| | dark_DLS | illuminated_DLS |
|---|---|---|
| **PDB Code** | 9S0H | 9S0I |
| **Data collection** | DLS (I24) | DLS (I24) |
| **Space group** | $P_{21\ 21\ 21}$ | $P_{21\ 21\ 21}$ |
| **Unit-cell parameters**[¶] | | |
| **a, b, c (Å)** | 64.49 (0.13), 71.47 (0.14), 206.50 (0.28) | 64.47 (0.23), 71.44 (0.33), 206.76 (0.41) |
| ***a, β, ɣ (°)*** | 90.00, 90.00, 90.00 | 90.00, 90.00, 90.00 |
| **Resolution (Å)** | 103.19 - 2.00 (2.03-2.00) | 103.38 - 2.20 (2.24-2.20) |
| ***I / σI*** | 7.0 (1.1) | 6.9 (0.8) |
| ***Rsplit (%)*** | 12.5 (90.5) | 13.9 (114.1) |
| ***CC 1/2 (%)*** | 99.1 (40.8) | 99.2 (34.3) |
| **Wilson B-factor (Å$^2$)** | 27.49 | 25.66 |
| **Completeness (%)** | 100 (100) | 99.9 (99.2) |
| **Refinement**[#] | | |
| **Rwork (%)** | 20.10 (34.08) | 20.01 (31.62) |
| **Rfree (%)** | 24.05 (36.30) | 24.92 (33.37) |
| **No. Atoms** | 6,737 | 12,906 |
| **Protein** | 5,973 | 11,840 |
| **Ligand/ion** | 436 | 872 |
| **Water** | 328 | 194 |
| **Average B-factor (Å$^2$)** | 33.98 | 45.68 |
| **Protein** | 34.05 | 45.18 |
| **Ligands/ion** | 30.44 | 52.63 |
| **Solvent** | 37.41 | 45.13 |
| **R.m.s. deviations** | | |
| **Bond lengths (Å)** | 0.008 | 0.007 |
| **Bond angles (°)** | 1.10 | 1.03 |
| **Ramachandran** | | |
| **Favoured (%)** | 97.6 | 97.6 |
| **Allowed (%)** | 2.4 | 2.4 |
| **Outliers (%)** | 0 | 0.0 |
| **Rotamer outliers (%)** | 2.9 | 2.3 |
| **Clashscore** | 5.9 | 6.0 |

[#] Values in parenthesis correspond to the highest resolution shell.

[¶] Values in parenthesis correspond to the standard deviation of the fitted normal distribution. Statistics for the highest-resolution shell are shown in parentheses.

#### Supplementary Table 8. Cryo-T dependent MX Data collection and refinement statistics

| | **cryo_light_180K** |
|---|---|
| **PDB Code** | 9S0G |
| **Data collection** | ESRF (BM07) |
| **Space group** | $P_{21\ 21\ 21}$ |
| **Unit-cell parameters** | |
| **a, b, c (Å)** | 63.99, 70.13, 206.27 |
| ***a, β, γ (°)*** | 90.00, 90.00, 90.00 |
| **Resolution (Å)** | 41.55 - 1.90<br>(1.93 - 1.90) |
| ***I / σI*** | 14.3 (1.0) |
| ***Rmeas (%)*** | 8.8 (246.7) |
| ***CC* (%)*** | 99.97 (79.8) |
| ***CC 1/2 (%)*** | 99.9 (46.7) |
| **Wilson B-factor (Å$^2$)** | 41.17 |
| **Completeness (%)** | 98.8 (99.5) |
| **Refinement[#]** | |
| **Rwork (%)** | 20.50 (36.24) |
| **Rfree (%)** | 23.94 (40.25) |
| **No. Atoms** | 6,685 |
| **Protein** | 5,915 |
| **Ligand/ion** | 442 |
| **Water** | 355 |
| **Average B-factor (Å$^2$)** | 47.60 |
| **Protein** | 47.75 |
| **Ligands/ion** | 45.79 |
| **Solvent** | 47.36 |
| **R.m.s. deviations** | |
| **Bond lengths (Å)** | 0.008 |
| **Bond angles (°)** | 1.05 |
| **Ramachandran** | |
| **Favoured (%)** | 96.5 |
| **Allowed (%)** | 3.2 |
| **Outliers (%)** | 0.3 |
| **Rotamer outliers (%)** | 0.9 |
| **Clashscore** | 4.4 |

[#] Values in parenthesis correspond to the highest resolution shell.

Statistics for the highest-resolution shell are shown in parentheses.

**Supplementary Table 9. Comparison of Co–C5′ and Co–C4′ distances in TR-SFX structures at 10 ns and 3 µs at two pump-laser fluences.**

| 10ns_12mJ/cm²_SACLA | | | | | 10ns _30mJ/cm²_SACLA | | | | |
|---|---|---|---|---|---|---|---|---|---|
| | Mono A | Mono B | Mono C | Mono D | | Mono A | Mono B | Mono C | Mono D |
| Bond lengths (Å) | | | | | Bond lengths (Å) | | | | |
| Co–C5′ | 4.78±0.1 | 4.56±0.1 | 4.01±0.1 | 4.22±0.1 | Co–C5′ | 4.77±0.1 | 4.54±0.1 | 4.15±0.1 | 4.14±0.1 |
| Co–C4′ | 3.65±0.1 | 3.64±0.1 | 3.27±0.1 | 3.40±0.1 | Co–C4′ | 3.62±0.1 | 3.54±0.1 | 3.29±0.1 | 3.38±0.1 |
| **3µs_12mJ/cm²_SACLA** | | | | | **3µs _30mJ/cm²_SACLA** | | | | |
| | Mono A | Mono B | Mono C | Mono D | | Mono A | Mono B | Mono C | Mono D |
| Bond lengths (Å) | | | | | Bond lengths (Å) | | | | |
| Co–C5′ | 3.69±0.1 | 3.30±0.1 | 3.20±0.1 | 3.19±0.1 | Co–C5′ | 3.57±0.1 | 3.16±0.1 | 3.04±0.1 | 3.01±0.1 |
| Co–C4′ | 2.53±0.1 | 2.32±0.1 | 2.22±0.1 | 2.29±0.1 | Co–C4′ | 2.44±0.1 | 2.18±0.1 | 2.12±0.1 | 2.12±0.1 |

**Supplementary Table 10. Atomic charges and spin densities for selected atoms in the cluster model.**

| | Natural charge | | | | Spin density | | |
|---|---|---|---|---|---|---|---|
| | Co | C4′ | C5′ | Ribose[1] | Co | C4′ | C5′ |
| HCbl + anhAdo (**1**) | -0.243 | 0.288 | -0.535 | 0.455 | 0 | 0 | 0 |
| [Co···H···C5′]$^{‡}$ (**2A**) | -0.031 | 0.367 | -0.625 | 0.517 | 0 | 0 | 0 |
| Co···H···C5′ intermediate (**3A**) | 0.046 | 0.395 | -0.678 | 0.522 | 0 | 0 | 0 |
| [Co(II) + C4′·]$^{‡}$ (**4A**) | 0.271 | 0.318 | -0.77 | 0.198 | 0.909 | -0.78 | 0.072 |
| Co–C4′ adduct (**5A**) | 0.107 | 0.298 | -0.726 | 0.200 | 0 | 0 | 0 |
| Co–C4′ adduct (b)[2] | -0.001 | 0.277 | -0.721 | 0.144 | 0 | 0 | 0 |
| [Co···H···C4′]$^{‡}$ (**2B**) | 0.029 | 0.109 | -0.424 | 0.177 | 0 | 0 | 0 |
| Cob(II) + C5′· (a) (**3B**) | 0.166 | 0.028 | -0.416 | 0.207 | -0.772 | -0.086 | 0.860 |
| Cob(II) + C5′· (b) (**4B**) | 0.274 | -0.04 | -0.384 | 0.166 | -0.925 | -0.055 | 0.998 |
| Co–C5′ adduct (**5B**) | -0.044 | 0.029 | -0.457 | 0.165 | 0 | 0 | 0 |
| Co–C5′ adduct (b)[2] | 0.103 | 0.022 | -0.430 | 0.211 | 0 | 0 | 0 |

[1]The ribose moiety was defined as including both C4′ and C5′

**Supplementary Table 11. AdoCbl geometric parameters in the dark state.**

Selected geometric parameters for AdoCbl in the dark state form of the crystal structure and the QM/MM optimized model. The conformation of the ribose portion of the adenosyl ligand is listed.

| | Monomer A | Monomer B | Monomer C | Monomer D | QM/MM |
|---|---|---|---|---|---|
| Bond Lengths/Distances (Å) | | | | | |
| Co–C5' | 2.02 | 2.03 | 2.02 | 2.03 | 2.01 |
| Co–C4' | 2.94 | 3.05 | 3.00 | 3.00 | 3.09 |
| Co–$H_{C4'}$ | N/A | N/A | N/A | N/A | 3.21 |
| Co–$N_{H177}$ | 2.18 | 2.27 | 2.19 | 2.24 | 2.33 |
| C5'–C4' | 1.52 | 1.52 | 1.52 | 1.52 | 1.52 |
| Dihedrals (°) | | | | | |
| $\theta_0$ C1'–C2'–C3'–C4' | 12.77 | 14.59 | 10.63 | 8.03 | -2.63 |
| $\theta_1$ C2'–C3'–C4'–O1' | -26.67 | -30.77 | -26.87 | -26.78 | -22.23 |
| $\theta_2$ C3'–C4'–O1'–C1' | 31.63 | 36.74 | 34.05 | 37.24 | 40.95 |
| $\theta_3$ C4'–O1'–C1'–C2' | -23.32 | -26.92 | -27.27 | -31.95 | -43.07 |
| $\theta_4$ O1'–C1'–C2'–C3' | 5.33 | 6.55 | 9.18 | 13.48 | 27.20 |
| Ribose Conformation | | | | | |
| | 3'-endo | 3'-endo | 3'-endo | 3'-endo | 3'-endo |
